# Demographic history shaped geographical patterns of deleterious mutation load in a broadly distributed Pacific Salmon

**DOI:** 10.1101/732750

**Authors:** Quentin Rougemont, Jean-Sébastien Moore, Thibault Leroy, Eric Normandeau, Eric B. Rondeau, Ruth E. Withler, Donald M. Van Doornik, Penelope A. Crane, Kerry A. Naish, John Carlos Garza, Terry D. Beacham, Ben F. Koop, Louis Bernatchez

## Abstract

A thorough reconstruction of historical processes is essential for a comprehensive understanding the mechanisms shaping patterns of genetic diversity. Indeed, past and current conditions influencing effective population size have important evolutionary implications for the efficacy of selection, increased accumulation of deleterious mutations, and loss of adaptive potential. Here, we gather extensive genome-wide data that represent the extant diversity of the Coho salmon (*Oncorhynchus kisutch*) to address two objectives. We demonstrate that a single glacial refugium is the source of most of the present-day genetic diversity, with detectable inputs from a putative secondary micro-refugium. We found statistical support for a scenario whereby ancestral populations located south of the ice sheets expanded in postglacial time, swamping out most of the diversity from other putative micro-refugia. Demographic inferences revealed that genetic diversity was also affected by linked selection in large parts of the genome. Moreover, we demonstrate that the recent demographic history of this species generated regional differences in the load of deleterious mutations among populations, a finding that mirrors recent results from human populations and provides increased support for models of expansion load. We propose that insights from these historical inferences should be better integrated in conservation planning of wild organisms, which currently focuses largely on neutral genetic diversity and local adaptation, with the role of potentially maladaptive variation being generally ignored.

## Introduction

Historical climate variation has had a major influence on the current distribution of species genetic diversity [1]. The Pleistocene glaciations, in particular, resulted in major contractions in the geographical distributions of many species into refugia that persisted in unglaciated areas [2–4]. Postglacial range expansions often led to contact between ancestral populations previously segregated in different refugia [2,4]. The effects of long-term climate change combined with recent human-induced population declines in wild animal populations can foster genetic changes including a loss of genetic diversity, increased inbreeding, increased load of deleterious mutations, and a loss of local adaptation [5].

In this context, understanding the impacts of historical climatic oscillations on the demographic history of a given species is crucial. Namely, it is important to understand how historical demographic events influenced the present-day geographic distribution of the within-species genetic diversity across its range. Here it is noteworthy that while the majority of studies on wild populations have focused on patterns of neutral and potentially adaptive genetic diversity, the importance of potentially maladaptive variation (e.g. patterns of deleterious mutation load) has generally been ignored. By disentangling past and current drivers of range-wide genomic diversity, this information can inform management and conservation decisions [6]. Beyond conservation implications, such context provides a unique opportunity to address outstanding questions in evolutionary biology: How is the efficacy of selection affected by historical processes that resulted in population expansion or bottleneck [7,8]? What are the demographic conditions required to generate substantial differences in deleterious load among populations [7–9]?

A major challenge in understanding drivers of genome-wide patterns of diversity is that different demographic processes can lead to similar contemporary genomic footprints [10]. As populations diverge, they accumulate genetic incompatibilities forming barriers to gene flow [11], while the rest of the genome may continue to be freely exchanged. Consequently, the genomic landscape of divergence is expected to vary, with greater differences along the genome at genomic barriers as compared to genomic regions exhibiting ongoing gene flow. However, similar patterns of heterogeneous genome-wide divergence can also be due to genetic hitchhiking of neutral alleles linked to selective sweeps [12,13] or to background selection (BGS;[12,14,15]]). These two processes, collectively referred to as linked selection, result in a reduction in nucleotide diversity in the vicinity of the sites targeted by positive or purifying selection. Due to its diversity-reducing effect, linked selection can be modeled as a local reduction of effective population size (*Ne)* [15]. It is now recognized that neglecting linked selection can bias demographic inferences [16,17] or lead to false adaptive interpretations [18]. Inference methods that incorporate variation in local effective population size and migration rate can help better understand how demography unfolds through time.

An understanding of historical demography is also essential for a sound interpretation of patterns of deleterious mutation load observed among contemporary populations [7,9]. Population bottlenecks are predicted to reduce potential for local adaptation, but also to reduce standing genetic variation and the efficacy of selection [8,19]. In turn, a reduced efficacy of purifying selection leads to an increase in the number of deleterious variants segregating in a population, thus reducing its fitness, and can result in maladaptation and/or reduced adaptive potential in bottlenecked populations. From a conservation standpoint, populations harboring an elevated number of deleterious variants might need to be monitored more closely to minimize extinction risks [20].

Combining population genomics data with demographic modelling represents a powerful strategy to test alternative hypotheses about historical drivers of existing genomic diversity. Previous studies employing a similar approach have focused mostly on species with a narrow geographic range, such as those inhabiting small islands [21,22], on the verge of extinction [23–26], or that experienced a strong bottleneck [27]. Few studies, however, have used demographic modelling to understand how historical processes have shaped the geographical patterns in the distribution of genomic diversity in a more broadly distributed species. An exception to this observation is the vast literature on demographic reconstructions of human populations. Long-lasting debates in this literature regarding the role of demography in generating mutation load differences among populations [9,28,29] could benefit from studies of species displaying similarly complex demographic histories and broad geographic distributions.

Salmonid fishes are economically important species that have suffered recent demographic declines [30,31]. This is particularly the case for Coho salmon (*Oncorhynchus kisutch*), one of the five anadromous species of Pacific salmon that supports important recreational and indigenous subsistence fisheries, and which has suffered dramatic population declines (> 90%) over the last three decades in parts of its range [32]. A previous study investigated the range-wide population structure and demographic history of the species and found a cline of decreasing diversity from south to north, as well as some endemic diversity in small putative refugia [33] (see also [34]). This study indicated that Coho salmon may have survived the last glacial maximum (LGM, i.e. the Fraser Glaciation in British Columbia, and the McConnell/McCauley Glaciation in Yukon and Alaska; 23 to 18 Ky ago) in unglaciated areas of Haida Gwaii and Beringia in addition to areas south of the ice sheets. This study, however, predates the genomic era and could not eliminate alternative hypotheses regarding the origin and number of glacial refugia during the LGM. In North America, the species is currently distributed from California to Alaska [35]. Unglaciated areas that could potentially serve as glacial refugia persisted both north (e.g. the Beringian refugium in Alaska, the Yukon Territory of Canada and areas of Asia and the Bering Land Bridge) and south (e.g. all of the deglaciated area south of British Columbia, Canada) of the ice sheets [35–37]. Other unglaciated areas (e.g. Haida Gwaii in British Columbia) could also have been micro-refugia [38,39]. In this context, distinct demographic scenarios can be tested. Under a first scenario whereby populations expanded north from a single southern refugium, we predict: *i*) a latitudinal decrease in genetic diversity from south to north along with a pattern of isolation-by-distance (IBD), and *ii*) ancestral populations located in areas south of the ice sheets. Under a second scenario, populations expanded south from a single northern refugium, and we predict the opposite geographic pattern. The third scenario corresponds to the survival of populations in different refugia where we predict: *i)* the existence of clearly distinct genetic clusters, and *ii)* postglacial gene flow with signatures of secondary contacts, with contact zones displaying higher genetic diversity through postglacial admixture between different genomic backgrounds.

*In order to test these alternative scenarios, we generated genome-wide data from nearly 2,000 Coho salmon from California to Alaska, one of the most extensive genomic datasets for a non-model vertebrate species to date. First, to resolve the species demographic history, we used a modelling approach that accounts both for barriers to gene flow affecting migration locally*, and for linked selection affecting the rate of drift. Second, we hypothesized that demographic history shaped the pattern of deleterious mutation load, both within and among populations. We hypothesized that postglacial re-colonisation influenced levels of standing genetic variation and favoured the accumulation of deleterious mutations at the expansion front. In these conditions, we predicted that neutral genetic diversity should decrease as a function of the distance from the ancestral populations.

## Results & Discussion

### Latitudinal gradient in nucleotide diversity

To investigate the distribution of genetic diversity and population differentiation, we sampled 1,957 Coho salmon from 58 sampling locations throughout their entire North American range from California to Alaska (mean n = 34 fish per location, **Fig. 1a, S1 Table**). Samples were genotyped using a genotyping-by-sequencing (GBS) method that generated 82,772 high quality filtered single nucleotide polymorphisms (SNPs). Consistent with earlier work based on non-genomic data, our analyses indicate that the southern-most populations are the most ancestral and suggest that a single southern refugium explains most of the observed patterns in the distribution of genetic diversity. Under a single refugial expansion scenario, one would expect genetic diversity to be highest in the ancestral refugium and a linear decrease as a function of the distance from the source, as documented in humans [40]. To test this prediction, we plotted the distribution of genetic diversity as a function of the distance to the southernmost sample in our data. As expected, levels of genetic diversity (observed and expected heterozygosity, π_SNP_) were highest in formerly deglaciated refugial areas in the south (California, Cascadia, **Fig. 2a, Fig 1b**) and decreased as a function of distance from the southernmost site up to Alaska (r = 0.64, p < 0.0001**, Fig. 2a, S1 Fig, S2 Fig.**). Two populations however deviated from this linear pattern (Clackamas R in Cascadia and Upper Pitt R. in BC) and displayed higher genetic diversity than the Southern samples. We hypothesized that these two rivers may have undergone recent admixture from differentiated genetic clusters, locally increasing genetic diversity. Recent admixture between populations from several refugia can produce increased genetic variation [41], which could explain why most ancestral populations (relict populations from the refugia) are not systematically those displaying the highest genetic diversity. The Thompson River watershed (Thompson R. hereafter) in southern British Columbia was an exception to this latitudinal pattern and displayed the lowest average level of regional genetic diversity of all sampling locations, which we hypothesized resulted from bottlenecks (see below). The remaining samples from British Columbia were intermediate in genetic diversity.

**Figure 1).**
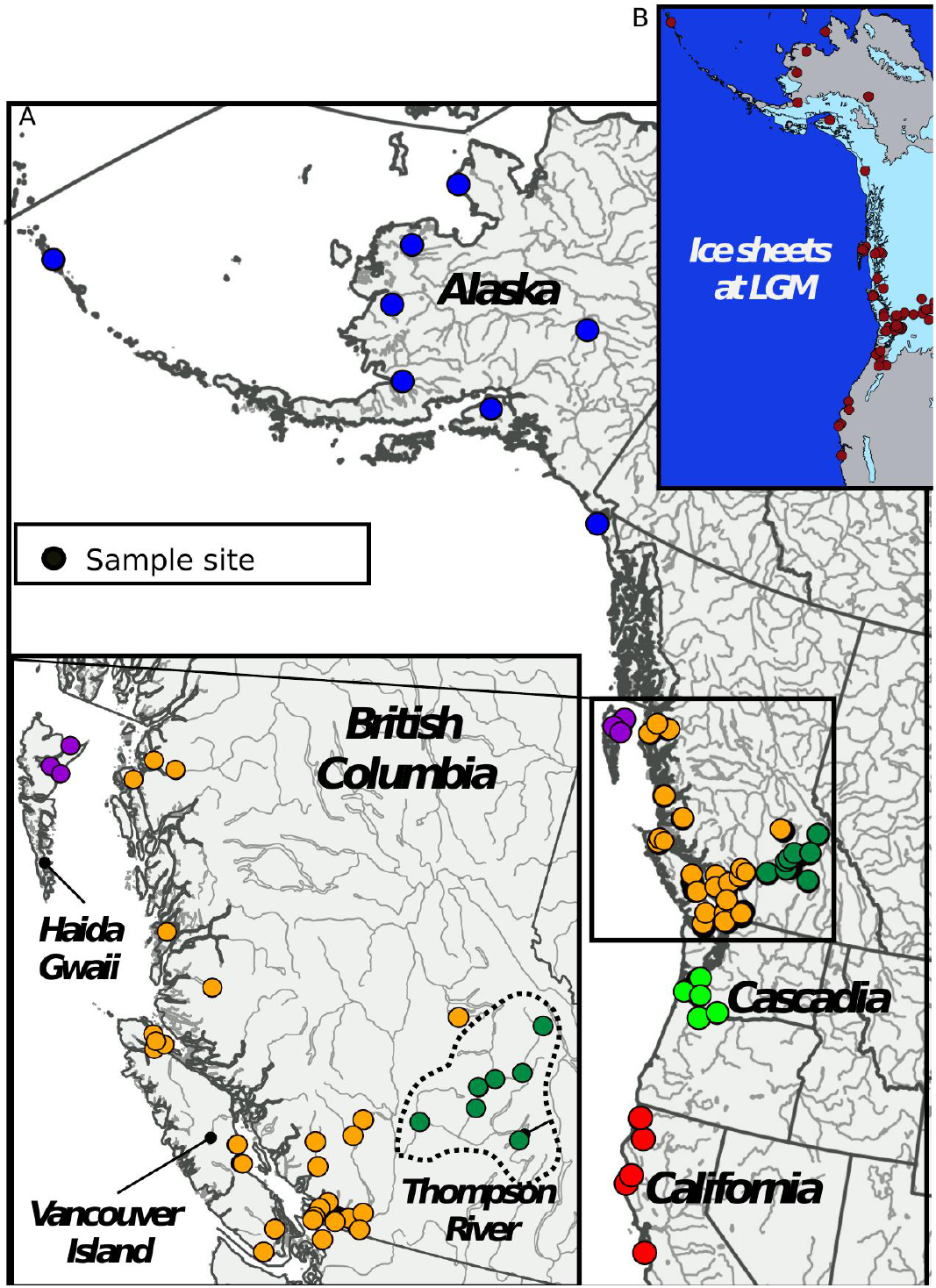
Sampling design. Sampling locations of 58 Coho salmon populations distributed across the species’ North American range. Each dot represents a sampling location. Inset: Map showing the extent of ice-sheet during the Last Glacial Maximum 13 Kya. Data modified from [94].

**Figure 2).**
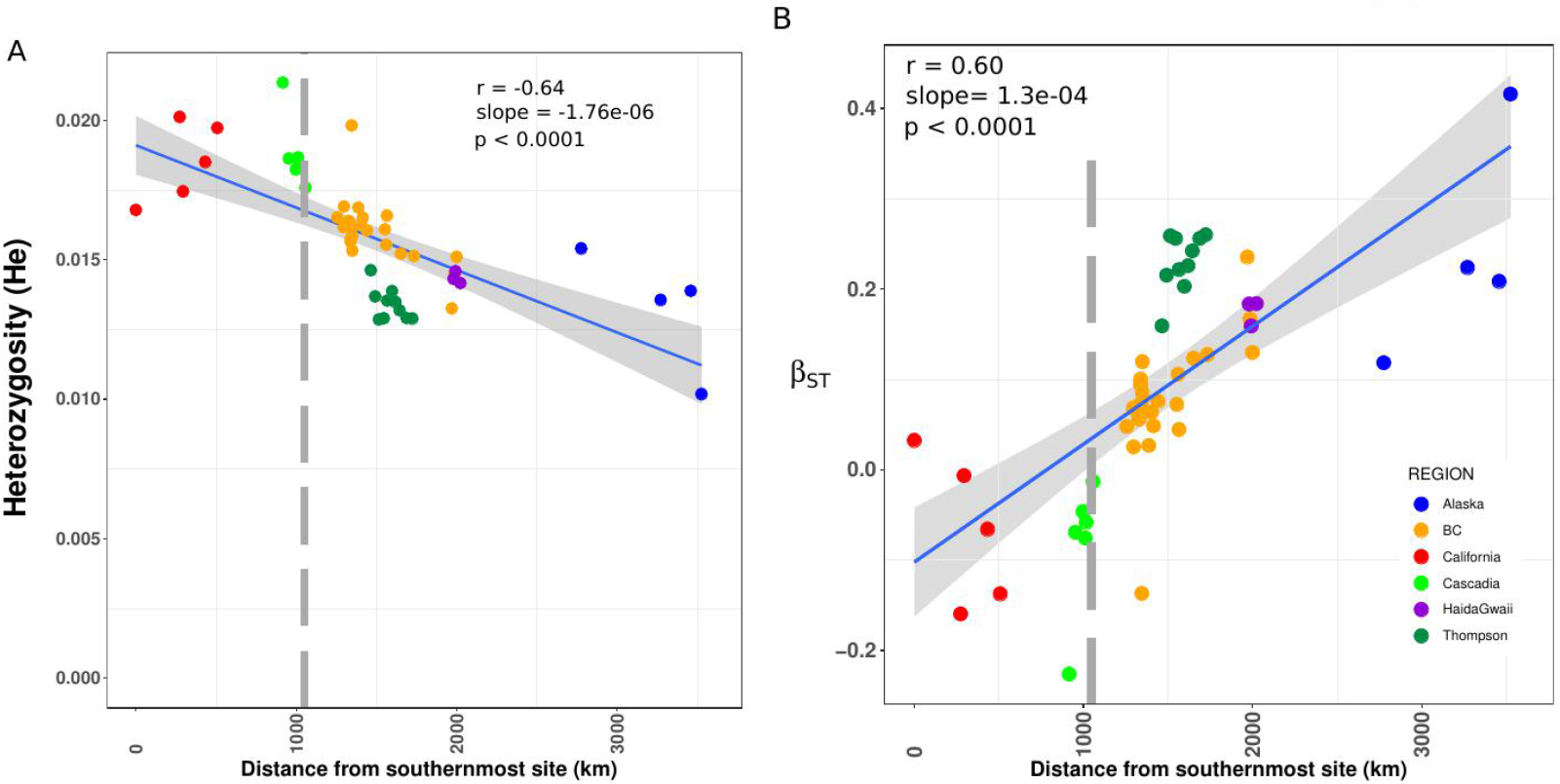
Genetic diversity and differentiation. A) Linear relationship between expected heterozygosity and distance from the southernmost population located in California. B) Linear increase in genetic differentiation as measured by β_ST_ as a function of the distance from the southernmost population located in California. Negative values indicate the most likely ancestral population. The relationship in A and B was tested using linear models. The grey vertical bar in panel A and B indicates the approximate location of the southern limit of the ice-sheet at the end of the last glacial maximum. The coefficient of correlation (r) is indicated for each plot. The grey shared area along the regression line corresponds to the 95% confidence level interval obtained from the linear models applied to each dataset.

### Present-day southern populations are the most ancestral

To help discriminate a single versus two population refugia, we aimed to identify the most ancestral populations among all samples. Under a single scenario refugium, the ancestral populations should be located preferentially in one area (e.g., South or North), whereas under a two-population scenario, we expected ancestral populations to occur in different areas located at both edges of the species’ distribution range (Beringia covering most of Alaska and the Yukon being known as an important glacial refugium for many species (reviewed in [2–3])). To discriminate between these alternative scenarios, we used two simple summary statistics, namely (1) the distribution of singletons among populations, and (2) the βst index recently developed in [42]. It was previously shown that older populations are expected to have accumulated a higher density of singletons [43]. Therefore, counting the number of singletons by sampling site after correcting for sample size (see methods) and averaging by regional groups may help identifying older populations. Alaska and Cascadian samples contained the highest number of singletons, with respective mean values of 753 (sd=116) and 650 (sd=152) singletons per site, consistent with the hypothesis of a refugium being present in these regions. Thompson and Californian samples had the fewest number of singletons (n_MEAN_ = 131 and 131 respectively). All regions differed significantly in their distribution of singleton (p < 0.0001) except the comparison between Thompson and California (S3 Fig) The observed signal in the Thompson was consistent with our above observations of reduced diversity. Similarly, we hypothesized that such reduced number of singletons in a few Californian samples reflects recent and strong population declines in some of these populations [44,45] (see also Supplementary Note S1). The similarity of the singleton distributions for both Casciadia and Alaska further emphasized the need for additional analyses (i.e. modeling approach implemented below) to help disentangle the scenarios of one versus two refugia.

Next, we used the β_ST_ coefficient to identify ancestral populations [42]. Unlike *F*_ST_ estimates [46], β_ST_ can account for the non-independence among populations and negative values are indicative of ancestral populations [42]. More specifically, the β_ST_ coefficient will be negative when a population has accumulated many private alleles at low to intermediate frequencies and the intensity of this value is dependent on the number of sampled populations [42]. As for singletons, older populations and populations of larger effective size have more time to accumulate rare alleles, resulting in negative values, as illustrated in human samples from Africa (see details in 42). Here, the β_ST_ indicated that the likely ancestral populations were in previously unglaciated areas corresponding to Cascadia (n= 5 localities), California (n = 3 localities) as well as one site from southern British Columbia **(Fig. 2b, S2 Table**). A linear decrease in β_ST_ with distance from the southernmost site was observed (r = 0.60, slope = 1.03e-04, *p <0.0001*) as expected under isolation-by-distance (IBD). Support for this IBD pattern was also observed using *F*_ST_ (**S4 Fig**, r = 0.66, slope= 4e-05, *p < 0.0001*) as well as Mantel tests (r = 0.64, *p < 0.0001*; and r = 0.72, *p < 0.0001* when removing Thompson R. populations). As for genetic diversity, the river CLA in Cascadia and UPT in British Columbia departed from the pattern and displayed the most negative coefficients (**Fig. 2b**). Finally, average pairwise *F*_ST_ across all populations was 0.095 and varied from 0.002 to 0.334 (**S5 Fig**), typical of anadromous species connected by gene flow [47].

To further confirm the number of putative refugia and better understand the extent of divergence among populations, we performed a principal components analysis (PCA). PCA can be used to summarize the distribution of genetic diversity with different expectations under discrete models of population genetic structure (as expected with different refugia) versus IBD under a one- or two-dimensional habitat model. Assuming K clusters, we expect that the top K-1 PCA axes should discriminate all these clusters [48]. On the contrary, under models of IBD, the top two axes should be correlated with geographical axes such as the longitude and latitude of populations’ locations [49]. Consistent with this latter hypothesis, we observed that the first axis was strongly correlated with latitude (r = 0.77, *p = 2e-16*), whereas the second axis was strongly correlated with longitude (r = −0.54, p = 2e-16). The result of the PCA (**Fig. 3a, see also S6 Fig**) is therefore consistent with a South to North spatial pattern of genetic structure, with the most divergent samples found in California, and a second geographical gradient from East to West, mainly reflecting the pronounced differentiation of the Thompson R. populations. Along these axes, populations followed an IBD pattern. These results were also supported by an MDS analysis (**S7 Fig**). A model-based analysis of population structure implemented in LEA [50] failed to reveal a clear number of distinct populations (K value). Instead, K values ranging from 30 to 60 all fit the data well (**S8 Fig**), providing little insight on the number of ancestral clusters in the data due most likely to the confounding effect of IBD [51]. While all these summary statistics lend support to the hypothesis of an expansion from a single ancestral refugium, a formal testing of alternative scenarios is required as the signature of secondary contact can be transient and both models of secondary contact and stepping stone can converge to the same IBD signal and apparent demographic equilibrium [10].

**Figure 3).**
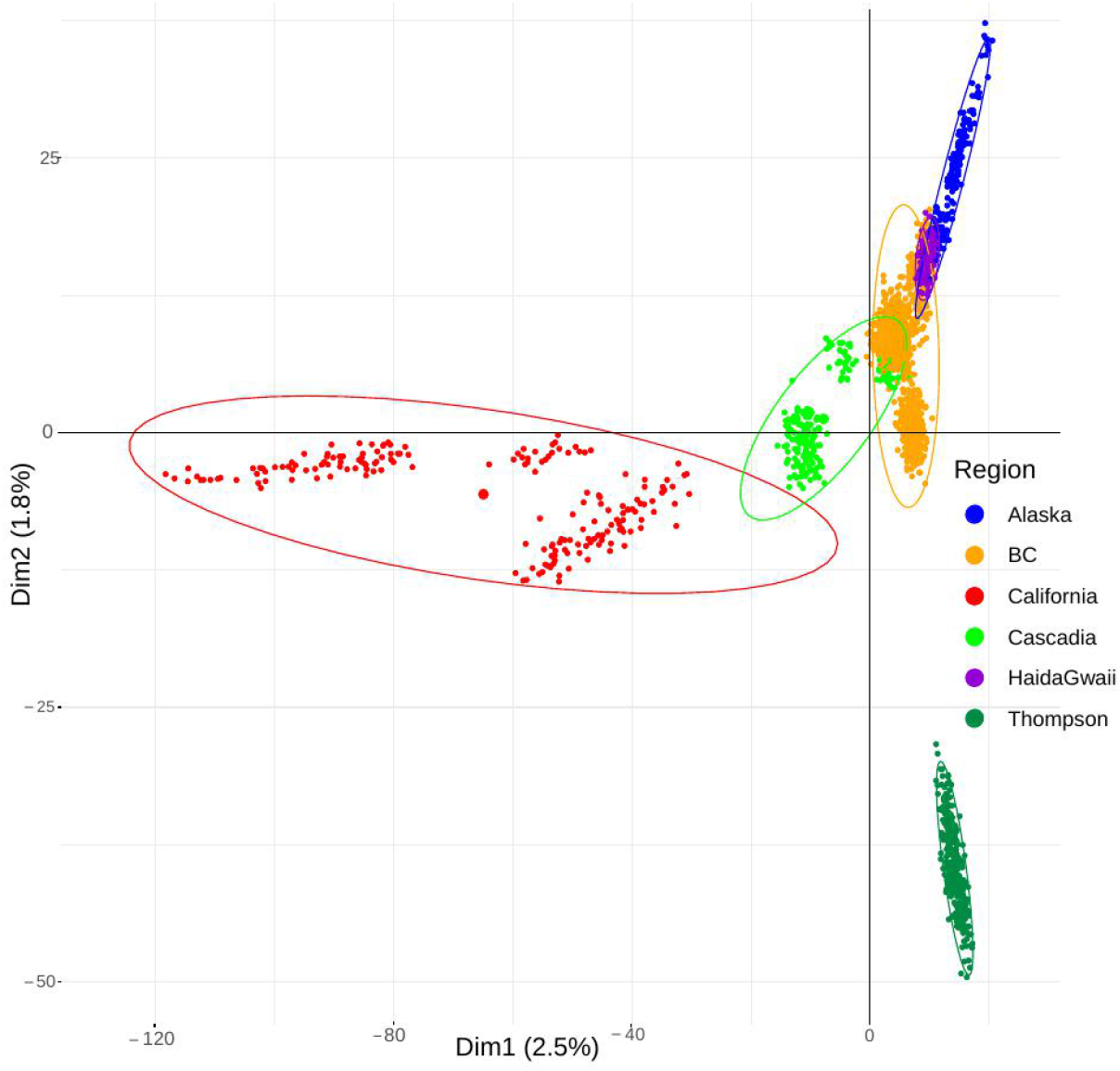
Genetic structure. Principal Component Analysis (PCA) summarising population genetic structure among 1,957 individuals based on the principal component axis 1 and axis 2. Each point represents an individual and the colours represent the major regional groups. The larger point represents the barycentre of each group.

Finally, using Treemix [52], we found that 99.1% of the total variance in allele frequency among populations can be explained by a single tree (S9 FigA-C) with four significant migration events (S9 FigD-E). California populations displayed pronounced genetic drift, corroborating the PCA results. Populations from Alaska (MSL River) and Thompson R. also displayed higher genetic drift, in line with our above results.We note that populations followed the south to north arrangement, with the samples from Cascadia displaying less drift than those located further north.

### Colonisation wave from a single major southern refugium

In order to formally assess the occurrence of one or more refugial origins for contemporary populations, we performed explicit model-based inferences of population divergence scenarios using ∂a∂i [53] and Fastsimcoal [54]. ∂a∂i was used to perform pair-wise population comparisons in order to 1) consider the confounding effects of linked selection and barriers to gene flow, and 2) calibrate a multi-population model to be used in Fastsimcoal. Under a model of a single refugium, either northern or southern, we expect to find support for a model of recent or ongoing divergence with gene flow (isolation with migration or IM) when comparing neighboring pairs. Following this hypothesis, IM models should be the best supported because the divergence time between the different populations should be recent, thus resulting in recent inferred divergence times. In addition, these populations are expected to still be connected by ongoing gene flow. Under a scenario with two ancestral refugia, we predict that comparing Alaskan populations (i.e. assuming that Coho survived in the Beringian refugium) to other genetic groups should reveal signals of secondary contact (SC). In addition to this evidence for postglacial gene flow, one can expect to infer a long divergence time between Alaskan populations and the rest of the populations. Finally we completed our inferences with a null model of strict isolation (SI) where populations would never have exchanged gene and a model of ancient migration (AM) where populations exchanged gene at the early stage of divergence only.

To reduce the number of pairwise comparisons and thus reduce the possibility of spurious results, we pooled individuals into higher order groups based on the PCA results (**Fig. 2a** and **S10 Fig**). The following six groups were constructed (named according to geographic labels): 1) California group 1, 2) California group 2, 3) Thompson 4) Cascadia 5) British Columbia and 6) Alaska. To avoid over-fitting of complex models, we performed model selection hierarchically. First, we compared all models (AM, IM and SC, SI) represented in **S11 Fig** assuming a constant size after divergence. Second, after choosing the best model based on ΔAIC, we incorporated changes in population size to estimate demographic parameter. All reported demographic parameter estimates are based on this last round of analysis. We compared Alaskan samples to all other groups (n = 4 pairwise comparisons). Then we compared the Thompson R. populations, which display an unusual pattern of reduced diversity and divergence from all other groups (n = 5 pairwise comparisons), resulting in a total of 9 pairwise comparisons.

Our inferences provided considerable support for an expansion from a single major refugium. Indeed, our analyses support models of divergence with ongoing gene flow between Alaska and all the other groups in three out of the four pairwise comparisons involving Alaska (**S12 Fig** for model fit and residuals and **S3 Table**; ΔAIC > 10; with the notable exception of Thompson, see below). These results are inconsistent with a major Beringian refugium, or at least a contribution of these northern populations to the present-day within-species variation. One out of the 4 comparisons (“California 1” versus “Alaska”) provided support for secondary contact. This result, was contrary to our expectations and highlights the well-known difficulty of distinguishing models of secondary divergence from primary divergence. Indeed, these two classes of models can quickly converge to the same state as observed here, namely large-scale isolation by distance and apparent equilibrium [10]. Interestingly, when any group is compared to Thompson R., all inferences find unambiguous support for postglacial secondary contact (**S3 Table**), suggesting that the Thompson R. might have act as a second refugium following the southern expansion and has been isolated from the main distribution.. The relative isolation of the Thompson R. would therefore be consistent with the lower genetic diversity observed in each local population within this area. It is noteworthy that in all models with support for secondary contact, the relative time of secondary contact represented on average 10% of the total divergence time. This proportion provided ideal statistical power to discriminate among models of divergence with continuous gene flow versus allopatric divergence followed by gene flow. Indeed, the signal of allopatric divergence is expected to be lost if gene flow was initiated too long in the past [17,55].

### Statistical support for ancestral expansion followed by recent demographic declines across all populations

We added a second layer of complexity by incorporating changes in population size in the models namely, expansions or bottlenecks. We hypothesized that these changes are likely to arise following postglacial expansion or through serial founder events during the recolonization process. We chose the best models identified through weighted AIC (**S3 Table**) and performed a round of parameter estimation. A salient result was that all populations exhibited a signature of expansion from the ancestral refugial population, with the effective population size at the divergence time being approximately 12 times higher than the ancestral population (**Fig. 4**). Then all populations declined to approximately 10-12% of their initial size (**Fig. 4**). More precisely, assuming a generation time of 3 years [56] and a mutation rate of 8e^−9^ bp/generation [57], our analysis inferred remarkably similar ancestral effective population sizes across all our pairwise comparisons (approximately 15,000 **Fig. 4, S4 Table**). Then we recovered a signal of expansion, with stronger expansion signals inferred in Cascadia and Alaska (**Fig. 4**). However, we observed some uncertainty in these parameter estimates (**S5-S6 Table**). This was expected because of the limited information available from the site frequency spectrum [58–59]. Moreover, we hypothesized that the life history trait of the species, particularly its strong natal homing behaviour [60] may result in different local evolutionary trajectories. Admittedly, such unmodelled process may further complicate the interpretation of demographic parameters.

**Figure 4).**
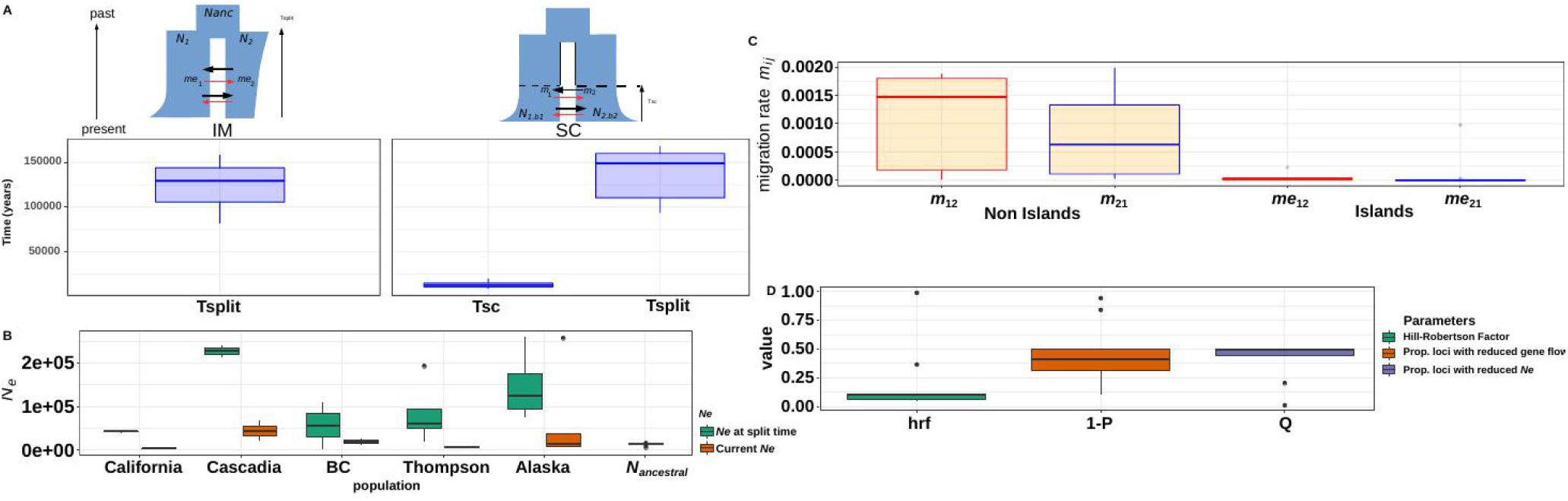
Inferences of demographic history using ∂a∂i. A) Estimates of divergence time (in years) between each major region as inferred by ∂a∂i under the best model (displayed in blue) based on SNP data. Plot displayed the parameter estimates for the best model for all pooled samples (IM = comparison between Alaska and all samples but Thompson SC = comparison between Thompson and all other samples as well as California versus Alaska) B) Estimates of effective population size Ne for each major region as inferred by ∂a∂i under each best demographic model. Effective size are based on the best model including population size change. Three extreme values (Ne > 2e6) were removed for readability). C) Estimates of migration rate among populations in neutral regions (m) of the genome and in areas of restricted recombination (m_e_) or islands of divergence. D) mean Hrf (Hill-Robertson factor) inferred across all comparisons. The Hrf estimates the extent of Ne reduction in areas affected by linked selection. Values are the averaged from multiple pairwise comparison across regions. Raw data are provided in table S4.

### Demographic inferences support glacial population isolation and recent postglacial recolonization

Our demographic inferences provided highly consistent divergence time estimates among all models and populations (**Fig. 4)**, with an estimated divergence time of 140 Kya (average = 107 KyA under IM and 137 Kya under SC; **Fig. 4, S5-S6 Table**) and suggesting a nearly simultaneous subdivision and colonization of the Northern range, likely before the Wisconsin glaciation. Accordingly, the median time of secondary contact (SC) among all models involving the Thompson R. was 13.5 Kya [min 9,200 – max = 20,100], corresponding to the onset of the last glacial retreat (**Fig. 4**), supporting the idea that the Thompson R. has evolved in isolation from the remaining populations for a long period of time.

Our models account for the confounding effects of selection at linked sites and that of the accumulation of local barriers to gene flow in the genome, two potential confounding factors that can lead to erroneous interpretations when they are not explicitly taken into account [12,17]. All models incorporating linked selection and restricted introgression along the genome always received the highest support. Here we modelled the effect of linked selection assuming that it can be approximated as a reduction of *Ne* in regions of low recombination [15]. Our inferences revealed that on average, 40% of the genome was undergoing reduced effective population size (Hill Robertson Factor in **S4 Table, Fig. 4**). Accordingly, the *Ne* values were reduced to approximately 20% of their initial values depending on the considered population (i.e. Hrf factor in **Fig. 4**). Obviously, the magnitude of the reduction would require further investigation as quantifying the effect of linked selection is likely more complicated than a simple rescaling of *Ne.* Still, these results provided increased support for a role of linked selection affecting the estimation of demographic parameters [17,61] and potentially shaping the species’ genomic landscape. Quantifying how background selection affects the species’ genome and shapes the genomic landscape of divergence is beyond the scope of this study and awaits further investigation. Finally, intrinsic barriers to gene flow reduced the estimated migration rate to nearly zero inside genomic islands and affected approximately 47% of the genome (**Fig. 4, S6 Table**).

To consolidate inferences made from ∂a∂i we constructed a global model based on ∂a∂i parameter estimates in Fastsimcoal v2.6 (Fig7). To limit the confounding of barriers to gene flow [11], linked selection [12] or gBGC [16], we restricted our jSFS to areas of high recombination, as inferred by LDHAT [62] from whole genome sequences [95]. Results largely validate our ∂a∂i inferences and provide rich information confirming strong population size reduction in California, Thompson, Alaska while revealing expansions in BC and Cascadia along with recent divergence of all these populations (details in **Supplementary Note S1, Supplementary Table S7 and Fig 7)**.

In summary, our results best supported a scenario whereby contemporary populations mainly originated from a single major ancestral refugium located south of the ice sheets in Cascadia/California, followed by a postglacial expansion and population divergence along the South to North recolonization axis. As a consequence, we expect that the South-North expansion should favour the accumulation of deleterious mutations at the expansion front toward the north [63,64], but also with the possibility for a pulse of deleterious variants from the isolated populations from the Thompson River watershed when they came into secondary contact with other expanding populations.

### The recent evolution of Ne shaped the current mutation loads

Our above demographic reconstruction revealed expansion from a major southern refugium. Consequently, we hypothesized that such a history has generated significant differences in the deleterious mutation load among populations, consistent with patterns documented in human populations [63–65]. Therefore, we hypothesized that populations having undergone a strong expansion following a bottleneck should display a higher load than populations of more constant size. Under this hypothesis we predicted a negative correlation between Tajima’s D, a classic population genetics statistic capturing population demography changes, and the ratio of non-synonymous to synonymous diversity (π_N_/π_S_), a commonly used proxy for the deleterious mutation load [66]. Accordingly, π_N_/π_S_ was significantly and negatively correlated with Tajima’s D (**Fig 5,** R^2^ =0.65, *p = 1e-15,* slope = 3.5). Considering our demographic inferences, this result illustrates how the postglacial recolonization changed local effective populations and shaped the present-day deleterious load variation among these populations. Interestingly, even after excluding the two most highly loaded populations (Clakamas and Upper Pitt River in Cascadia and BC respectively), the relationship between the π_N_/π_S_ and the Tajima’s D still remain strong (R^2^ = 0.42; *p = 4e-8,* slope= −2.7). Our results suggest that the Clackamas River populations exhibit high levels of admixture with the Thompson R populations (details in **S13 Fig**). As a consequence, we hypothesize that the post-glacial secondary contact with this population contributed to the introgression pulses of maladaptive alleles that are still detectable nowadays. This raises exciting future research questions regarding the role of past introgression and the maintenance of deleterious alleles [67]. It must however be noted that the Tajima’s D values were more widely distributed than expected under a strict model of population expansion from the South (assuming a linear decrease with distance, **S14 Fig**), suggesting additional sources of variation that remain to be identified and their effect evaluated.

**Figure 5).**
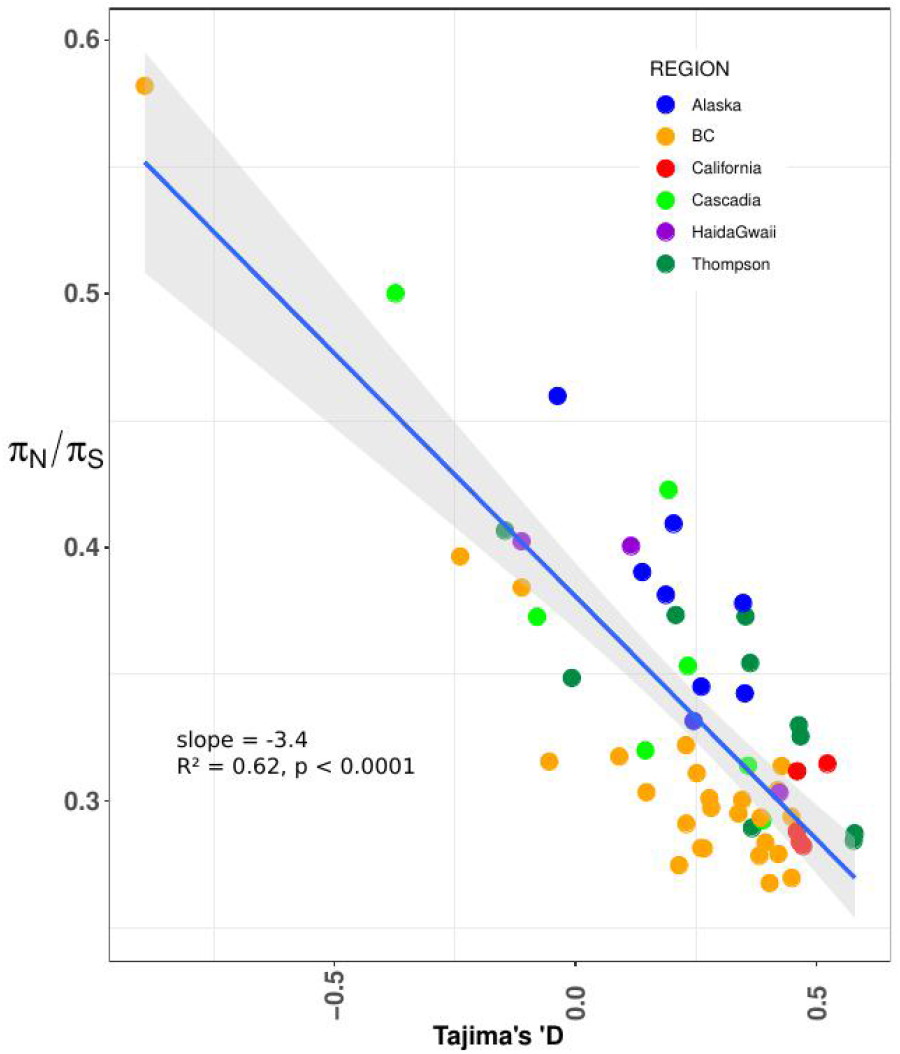
Recent demographic history shaped the deleterious mutation load: Correlation between Tajima’s D and the deleterious load (π**_>N_/ π_S_**). R^2^ = coefficient of determination of the linear model.

### Genetic surfing of deleterious mutations during the postglacial recolonization

One possible consequence of postglacial recolonization is that neutral and deleterious mutations may increase in frequency (I.e.surf) at the expansion front [63]. This situation occurs because populations at the expansion front exhibit smaller sizes that prevent the efficient purging of deleterious variants. Theory predicts (i) a higher recessive load (measured as the proportion of homozygous derived deleterious mutations) at the expansion front and (ii) an approximately constant total load (which can be approximated by the total number of derived deleterious mutations under an additive model [28,63,65]). We found support for these predictions.

First, we found a nearly linear relationship between the distance to the candidate source (defined as the one with the lowest β_ST_ and highest diversity) and the recessive load (linear models, p<0.003; R2= 0.13, **Fig. 6a**), validating theoretical expectations from range expansion [65]. This latter analysis was performed with PROVEAN [68] based on a total of 1,297 deleterious mutations for which we were able to identify the derived allele using the genome sequences of the 1) Chinook salmon (*Oncorhynchus tshawytscha*) [69], 2) Rainbow trout (*O. mykiss*) [70] and 3) Atlantic salmon (*Salmo salar*) [70] (see methods). Second, and as expected, the total load was not significantly correlated with the distance to the source (R^2^ =0.02, p = 0.96, **Fig. 6b**). Since PROVEAN is primarily designed for Human and Mouse (although it has been used successfully in non-mammalian studies [71,72]), we also plotted the distribution of π_N_/π_S_ as a function of the distance to the southernmost site. The π_N_/π_S_ ratio was significantly correlated with the distance from the southernmost site **(Fig. 6c**, *p = 0.05;* R^2^ = 0.05, for all populations). After excluding Clackamas and Upper Pitt populations (hypothesized to display increased load due to admixture), the correlation significantly increased (R^2^ = 0.2, *p = 0.0003*, slope =2.395e-05). Third, we observed significant differences in the derived allele frequency (DAF) at predicted deleterious non-synonymous mutations among regions (Kruskal-Wallis chi-squared = 100.57, df = 5, *p-value < 2.2e-16*) as well as among populations (Kruskal-Wallis chi-squared = 638.59, df = 57, *p-value < 2.2e-16,* S11 Table) with higher values indicative of less efficient purging (See supplementary **Fig S15-S18 and S9-S10 Table for details**). To the best of our knowledge it is the first time that such correlations between different proxies of the load and distance to the putative refugial origin are reported other than in studies on humans [28,64] and a few in plants [73,74] or bacteria [75]. As such, our two major results regarding Tajima’s D and the evolution of expansion load as a function of the distance aligned perfectly with the inferred expansion history and recent change in effective population size in each local population.

**Figure 6).**
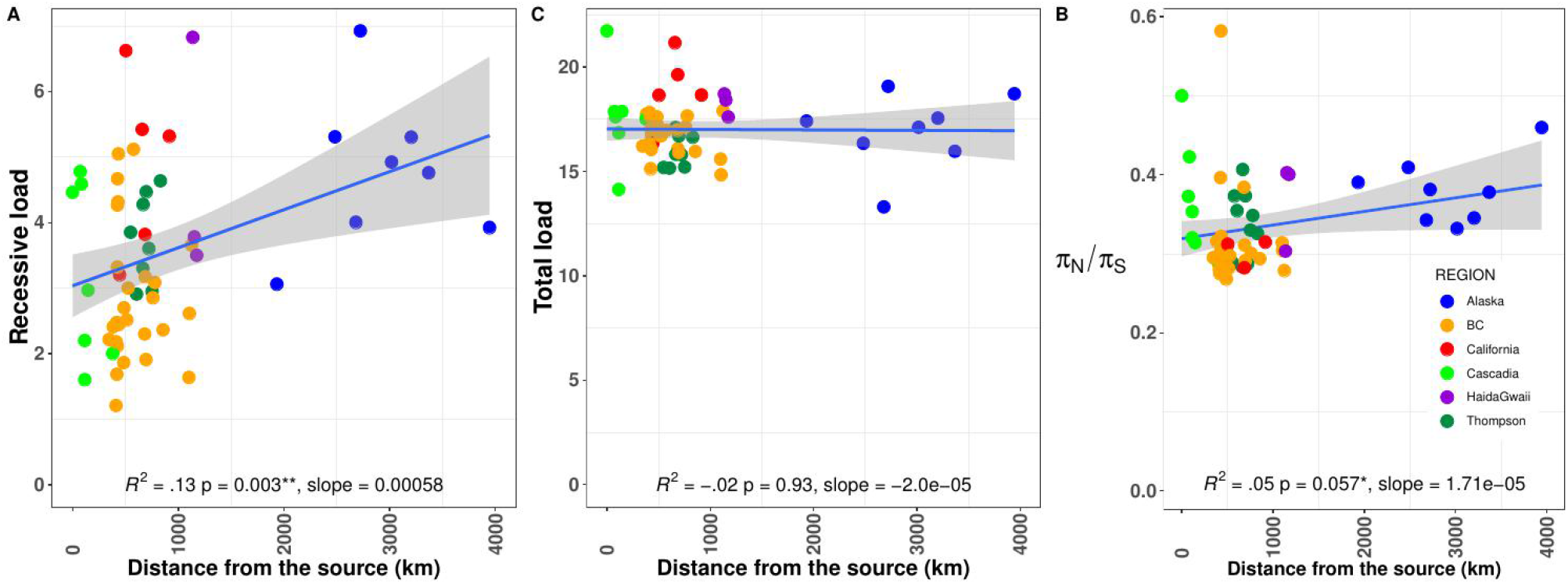
Expansion history and geographic pattern of deleterious mutation load: Correlation between the distance to the southernmost site and the deleterious load measured as (A) the distribution of homozygous derived putatively deleterious mutations (i.e. recessive load). B) the total load expected to be approximately constant among all populations) and C) the π_N_/π_S_ ratio. R^2^ = coefficient of determination of the linear model.

Finally, a general prediction from population genetic theory is that deleterious mutations in a heterozygous state should be more frequent than in a homozygous state, especially in populations with higher effective sizes, where selection should be more effective at purging mutations with increased homozygosity [19]. Accordingly, we found that 77% of deleterious mutations were maintained in heterozygous states across all samples, as reported in other species [71]. Also, salmon from Alaska, Haida Gwaii, and California populations harboured a significantly higher number of deleterious mutations in a homozygous state when compared to Cascadia or British Columbia (Wilcoxon-test, *p < 0.01*) **(S17a Fig, S11 Table**), a result that is in line with our finding of increased expansion load in **Fig 6a**. When considering the total load of derived deleterious mutations, we found that, on average, there were indeed significantly more putatively deleterious variants per individual in California, Cascadia, and Haida Gwaii populations than in fish from Alaska, British Columbia, or the Thompson R. watershed (**S10 and S11 Table, S17b Fig**, Wilcoxon-test, p < 0.01), although these differences were modest. Accordingly, we observed that π_N_/π_S_ varies as a function of the levels of nucleotide diversity (π_S_). We indeed found a positive relationship between π_N_/π_S_ and π_S_ (slope = 175.5; R^2^ = 0.45, *p < 0.0001,* **S19 Fig**). We note again the same two populations (Clackamas River & Upper Pitt) were strongly contributing to the linear regression. However, the relationship remained significant after their exclusion (R^2^ = 0.10, *p < 0.0078*). This result provides increased support for our above findings that populations with higher effective population size also displayed a higher deleterious load when considering mildly deleterious variants in heterozygous states (those that are more likely to contribute to π_N_). This suggests strongly that deleterious mutations are efficiently purged in populations of large *Ne* while mildly deleterious mutations would still segregate at appreciable frequency [23], as observed elsewhere [21,22,26]

The role of historical demography in generating differences in the mutation load at the population level remains debated among evolutionary biologists, particularly among human geneticists [7,9,28,29,76]. Our results are however consistent with the nearly neutral theory and the accumulation of deleterious mutations in populations on the expansion front, thus producing a so-called expansion load [63,65]. It is noteworthy how well our findings mirror the ‘out-of-Africa’ expansion model of human populations [77], despite the fact that the investigated geographic distance for the Coho salmon is an order of magnitude smaller than that involved in the human studies, and that the presence of additional smaller refugia such as Thompson R. (as opposed to the sole African ancestral origin for humans) led to more challenging inferences.

## Conclusion

In this study, we provided rare evidence of how demographic modelling can provide valuable information for conservation genomics. We found considerable statistical support for a recent postglacial demographic history in the Coho salmon, consistent with a main southern refugium, which generated relatively simple latitudinal gradients in nucleotide diversity and mutation load. Complex demographic processes including past population expansions or periods of isolation and secondary contact, have also been reconstructed with state-of-the-art demographic modelling methods. This was made possible by jointly considering the confounding impact of linked selection for such demographic reconstructions. Altogether, this postglacial history, as well as some additional demographic changes, have influenced the efficacy of selection and favoured the accumulation of deleterious mutations in the most distant populations from the main refugium (**Fig 7**). Using the Coho salmon as a case study, we demonstrated how these approaches are meaningful for conservation genomics and provide opportunities to disentangle the impacts of historical *vs.* contemporary drivers of population declines. Such investigations also allow assessing the distribution of all selected variants, including not only the beneficial, but also deleterious ones.

**Figure 7:**
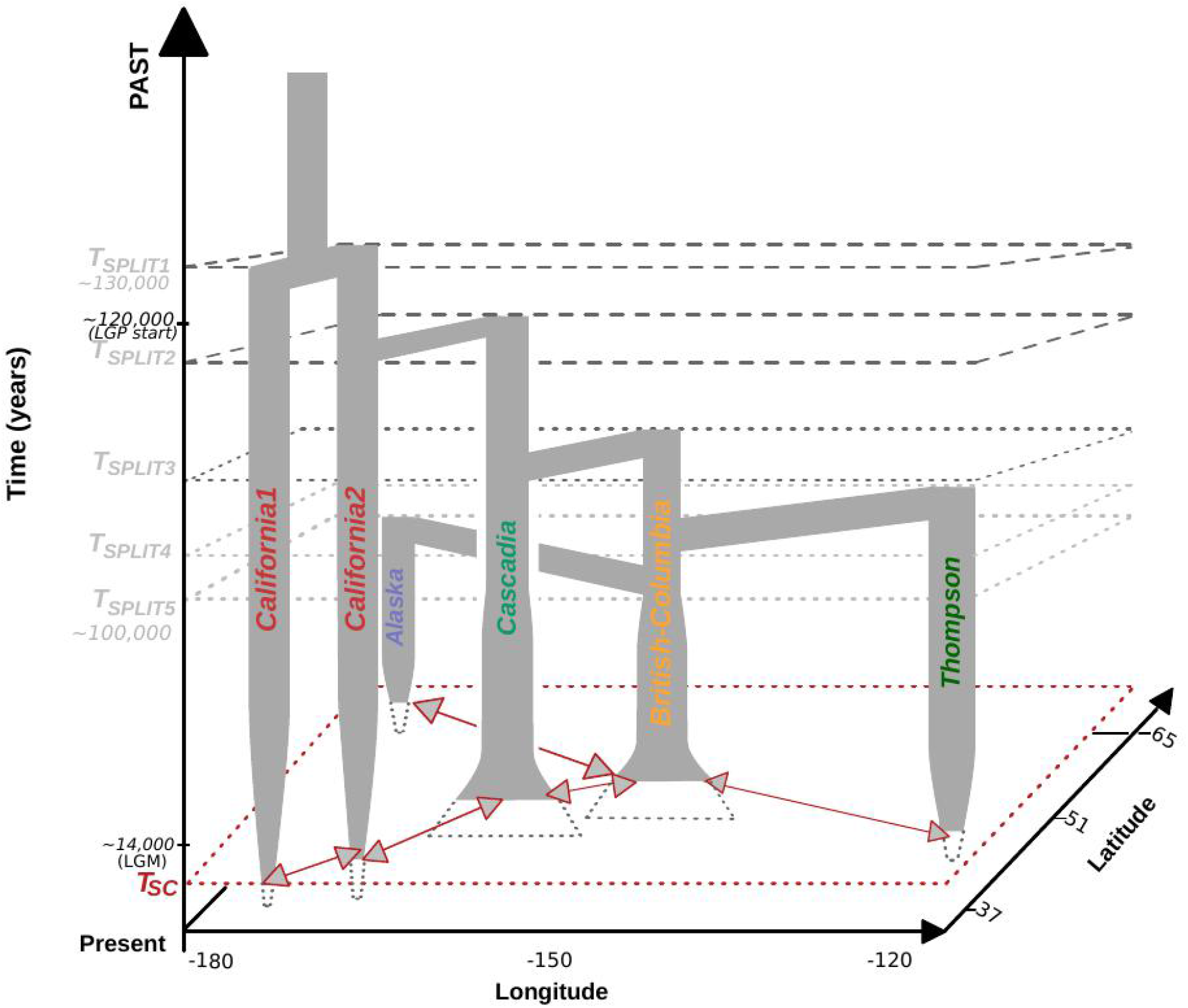
Simplified spatio temporal representation of the coho salmon demographic history based on ∂a∂i and Fastsimcoal results. Each bar represents a population branch. the width of the branch is not proportional to the population effective size and do not account for historical population size changes. The split time are not to scale and geographic positions are approximate. Tsc = Time of secondary contact. Red arrows represents major gene flow events between neighboring populations. Details about gene flow event can be found in table S5 and table S7. LGP = approximate start time of the Last Glacial Period [96]. LGM = Last Glacial Maxima Beringia was unglaciated during most of the pleistocene [97]. See results for details of the grouping.

## Methods

### Genotyping by Sequencing

A total of 2,088 individuals were collected from 58 sample sites located along the Pacific coast from California to Alaska (S1 Table and Figure 1). DNA was extracted from all individuals and sequenced using a GBS method (protocol detailed in [78]). Reads were aligned to the Coho salmon reference genome v1 (GCF_002021745.1) using bwa-mem 0.7.13 [79]. Samtools v1.7 [80] was used to keep reads with a mapping quality above 20, remove supplementary alignment and unmapped read. Variants were then called with Stacks v1.46 [81]. To do so, the module “pstacks” was used with a minimum depth of 5, and up to three mismatches were allowed in catalog assembly. The module “populations” was run to produce a vcf file that was filtered with a custom python script. We performed stringent filtering to remove SNPs that were 1) genotyped in less than 60% of the individuals; 2) at a mean depth of sequencing (averaged across all individuals) below 7, and 3) with observed heterozygosity above 0.60, thus resulting in 93,000 SNPs. The pipeline for SNP calling is available on github at https://github.com/enormandeau/stacks_workflow/releases/tag/coho_demography_paper. Next, we removed any individuals with more than 5% missing data and finally only kept SNPs present in at least 95% of the individuals yielding a total of 82,772 filtered SNPs for 1,957 individuals. Remaining filtration was done according to the requirement of each analysis performed below. To perform explicit demographic analyses based on the site frequency spectrum, the likelihood approach implemented in ANGSD v0.930 [82] was used instead of the commonly used Stacks pipeline because it is less biased than the genotype calling approach [83]. We first used the SAMTools model [80] in ANGSD to estimate genotype likelihoods from BAM files using only reads with a minimal base quality score of 20 and a minimal mapping quality score of 30 across all individuals. We further filtered out reads with a depth below 5 and above 100. Given the high residual tetrasomy in the species, the upper bounds enabled removing the high rates of paralogs that would otherwise generate an excess of high frequency shared variants in the jSFS. We verified that such variants were effectively removed in all jSFS before ∂a∂i fitting. We further constructed 1D SFS for all localities to then compute Tajima’s D value as a summary of recent effective population size change.

### Genetic diversity and ancestral populations

For each sampling location we estimated the observed heterozygosity and π using vcftools 0.1.16 [84] and Hierfstat [85]. The most likely ancestral populations were identified using β_ST_ [42]. A total of 1,000 bootstraps was performed to obtain the 95% confidence intervals around the β_ST_. Weir and Cockerham’s *F*_ST_ estimator θ [46] was computed in vcftools. We measured the relationship between observed heterozygosity, β_ST_, *F*_ST_ and the distance to the southernmost site using linear models with the lm() function implemented in R. We also verified the relationship between *F*_ST_ and the distance to the southernmost site using Mantel tests with 10,000 permutations. Vcftools was also used to identify singletons (i.e. variants present in one single individual across the whole dataset). The power to discover singletons is however dependent upon the sample size. Therefore, we reduced the size of each population to the smallest size of 13 by randomly sampling individuals in each population. We repeated the procedure 200 times to obtain standard deviations of the distribution of singletons (S3 Fig). We then computed the averaged number of singletons at the regional level as well as levels of private and shared polymorphisms (S2 Fig and S3Fig.). We tested the significance of differences in singletons distribution acrross the 200 replicated using a pairwise Wilcoxon test corrected with the Benjamini Hochberg method for multiple tests [86].The scripts are available on GitHub at https://github.com/QuentinRougemont/utility_scripts

### Population structure, admixture and gene flow

Levels of ancestry and admixture proportions were inferred with the snmf function implemented in the R package LEA [50], allowing only less than 5% of missing data. We then kept a set of SNPs in approximate linkage equilibrium by removing pairs of variants with r^2^ greater than 0.2 (option --indep-pairwise 50 10 0.2) resulting in 40,632 SNPs. K-values ranging between 1 and 60 were tested and cross-validations were obtained using the cross-entropy criterion with 5% of masked genotypes. The default value for the regularization parameter was used to avoid forcing individuals into groups and hence underestimating admixture. Similar results were obtained from Admixture [87] and are not presented here. The genetic relationship among all salmon was assessed using a PCA with the R package ade4 [88] based on the LD-pruned dataset (40,632 SNPs). We used a 1% minor allele frequency (MAF) threshold and allowed less than 4% missing data. Formal tests of admixture were performed using Treemix [52] using the LD-pruned dataset of 40,632 SNPs and without any MAF threshold. An MDS was also constructed using plink and plotted with the ggplot2 [89] R package. We ran Treemix allowing up to 20 migration events and performed 500 bootstrap replicates of the best model to obtain robustness of the nodes. The “best” model was inferred after inspecting the relevant migration edges by measuring the percentage of variance explained as migration edges were added to the tree, as well as by assessing the p-value associated to each migration edge. A total 500 bootstrap replicate runs were performed under the “best” model and under a model without migration to infer the robustness of the nodes. The scripts are available on GitHub at https://github.com/QuentinRougemont/treemix_workflow

### Explicit demographic inferences

We tested alternative hypotheses of divergence from a single southern or northern refugium versus divergence from two or more refugia (e.g. Alaska & California). In the first case, by comparing different regional groups, we expected a model of divergence with ongoing migration (IM) to be supported, or alternatively a model of divergence with ancient migration if gene-flow has stopped recently (AM) following the divergence and recent colonization of rivers. In the case of divergence into multiple refugia we expected models of secondary contact between putative groups to be best supported. We further expected postglacial gene flow between the historical refugia to leave detectable traces that should correspond to the signals of secondary contact. In the unlikely case of no gene flow between regional groups, we expected a model of strict isolation to be most likely. All divergence scenarios are represented in Fig S12 and initially described in [17,90]. The four major models tested included a model of Secondary Contact (SC), a model of Strict Isolation (SI), a model of Ancient Migration (AM) and a model of Isolation with Migration (IM).

The models shared the following parameters: the ancestral populations of size N*anc*, splits at time *T_split_* into two daughter populations of size *N*1 and *N*2. Under the SI model, no gene flow occurs between the two populations. Under AM, gene flow occurred between *T_split_* and *T_am_* and is followed by a period of strict isolation. Under IM, gene flow occurs at a constant rate at each generation between the two populations. Gene flow can be asymmetric, so that two independent migration rates *m_12_* (from population 2 to 1) and *m_21_* (from population 1 to 2) were modeled. Under the SC model, the population evolved in strict isolation between *T_split_* and until *T_sc_* where a secondary contact occurs continuously up to present time. Gene flow is modeled as *M = 2Nref.m*. In ∂a∂i, heterogeneous introgression was modeled using two categories of loci occurring in proportions *P* (i.e., loci with a migration rates *M_12_* and *M_21_*) and *1-P* (i.e., loci with a reduced effective migration rates *Me_12_* and *Me_21_*) across the genome. The same procedure was used to account for linked selection by defining two categories of loci with varying effective population sizes (proportion 1-Q of loci with a “neutral *N_e_*” and a proportion Q of loci with a reduced effective population size due to either selection at linked site). To quantify how linked selection affects reduced *Ne,* we used a Hill-Robertson scaling factor (Hrf) to relate the effective population size of loci influenced by selection (*Nr = Hrf* * *Ne*) to that of neutral loci (*N_e_*). A hierarchical approach was used to avoid over-fitting: first we compared models assuming constant effective population size. Second, the best identified models were modified to incorporate population expansion or decline, as expected given the observed distribution of genetic diversity. Population expansion was implemented using two additional parameters for population 1 and population 2, allowing each population to either grow or decline exponentially.

Models were fitted using the diffusion theory implemented in ∂a∂i [53] and includes the effect of linked selection and barriers to gene flow as detailed in [17,90]. ∂a∂i uses the SFS as a summary of the data. For a given demographic model, the SFS is computed using diffusion approximation and compared to the empirical SFS using AIC. Here, we started from the whole file containing 200,000 SNPs and used one single SNP per GBS locus and filtered the data to minimize missing data. No MAF filter was used and singletons were kept to avoid ascertainment bias in estimates of demographic parameters. For each model, 20 independent replicate runs were performed and only models with the lowest AIC and ΔAIC were kept. A model was classified as “ambiguous” and not used for parameter estimation if ΔAIC between the best model and second-best model was below 10. The Godambe information criteria was initially used to evaluate parameter uncertainties based on 100 bootstrapped datasets, which resulted in large confidence intervals in a few pairwise comparisons. Therefore, we then performed parametric bootstrap. More specifically, we converted ∂a∂i estimates into ms [91] estimates to construct sets of 100 bootstrapped datasets over the different categories of the SFS (i.e. neutral SFS, SFS with reduced rate of migration, SFS in low recombination areas undergoing linked selection, SFS in areas of normal recombination) that were summed and then fitted with ∂a∂i. The new estimates were then converted into demographic units to obtain the 95% confidence intervals. The whole pipeline is available at https://github.com/QuentinRougemont/DemographicInference.

#### Inferring a global divergence history

While the ∂a∂i analyses provided valuable results regarding the demographic history and how it is confounded by linked selection and barriers to gene flow, this analytical framework is currently limited to pairwise comparisons, which does not allow painting a global scenario of the species divergence history. Therefore, we build upon the ∂a∂i models and other observed statistics to construct a global scenario of population expansions from the South to the North with and apostglacial secondary contact between Thompson and the main distribution. This scenario was then tested using Fastsimcoal v2.6 [54] and is described in supplementary note S1.

### Genetic load estimation from the GBS data set

#### Estimating ancestral and derived alleles

We identified derived and ancestral allele using the genomes of three outgroup species, the chinook salmon, the rainbow trout, and the Atlantic salmon, to classify SNPs as ancestral or derived. These informations were subsequently used to analyze the load of deleterious mutation. Whole genome data for the chinook salmon (n = 3 individuals) were provided by one of us (B. Koop, unpublished), whereas rainbow trout (n = 5) and Atlantic salmon (n = 5) data were downloaded from NCBI Sequence Read Archive (rainbow trout, SRA, Bioproject: SRP117091; *Salmo salar* SRA Bioproject: SRP059652). Every individual was aligned against the Coho salmon V1 reference genome (GCF_002021745.1) using GATK UnifiedGenotyper and calling every SNP using the EMIT_ALL_SITES modes. We then used a python script to determined the ancestral state of the GBS SNPs if 1) the SNP was homozygous in at least two of the three outgroups, and 2) match one of the two alleles identified in Coho salmon. Otherwise, the site was inferred as missing and not used in subsequent analyses.

#### Measuring the deleterious load through piN/piS ratio

The ratio of non-synonymous diversity to synonymous diversity is a commonly used metric to quantify the deleterious mutation load [66]. To do so, we first generated aligned fasta sequences and then computed summary statistics. More specifically, for every individual, we ran GATK v4.1.2.0 on the bam file generated above following GATK best practices. We generated individual gVCF files then combined individuals into populations using the CombinedGVCFs module from GATK and finally performed a joint genotyping using the all-site options. We then reconstructed fasta sequences from VCF files for every individual by reconstructing two genomic sequences with a pipeline modified from [92]. We then used cutSeqGff.py to subsample and clean sequences considering several genomic features (CDS, introns, intergenic regions) from gff files. The resulting alignment was then used to compute the π_N_/π_S_ ratio in sequences containing 4-fold degenerate codons only. We then tested a number of correlations between the load (π_N_/π_S_) and 1) π_S_ used as a proxy of the long-term effective population size; 2) Tajima’s D used as a proxy for historical change in population size and 3) distance to the Southernmost site. Significance of the correlation were tested using the lm() function in R. Plots were drawn using ggplot2 [87]. All pipelines to run GATK and compute deleterious load are available at *:https://github.com/QuentinRougemont/gatk_haplotype and https://github.com/QuentinRougemont/piNpiS.*

#### Measuring damaging impact of non-synonymous alleles

While the computation above provides insights into the deleterious mutation load, it does not allow distinguishing among recessive and additive load nor testing for differences in allelic frequencies. We therefore further tested differences in mutation load among populations as follows. The software Provean [68] was used to predict whether a given non-synonymous mutation was deleterious with a threshold score of −2.5 or less using the pipeline available at https://github.com/QuentinRougemont/gbs_synonymy_with_genome. We analysed the data in two ways: first we counted the total number of putative homozygous deleterious alleles per individual as well as the total number of deleterious alleles (both in homozygous and heterozygous states) using: N_total_ = 2 Χ N_homo_ + N_hetero_ [34]. These two metrics are expected to be proportional to the recessive and deleterious load respectively [28,65]. These individual values were then averaged per population and major regional group (i.e., California, Cascadia, British Columbia, Haida Gwaii, Thompson, and Alaska). We then computed derived allele frequencies (DAF) in all sampling locations and across mutation categories (synonymous, non-synonymous, and non-synonymous deleterious) and tested for significant allele frequency differences among populations in non-synonymous and non-synonymous deleterious mutations using Wilcoxon rank sum tests. DAF spectra were constructed for all populations separately. For the ease of visualisation, we also constructed DAF spectra by region for a sample of size n = 100 individuals (Fig 7A). This size was chosen according to the smallest sample size of the three combined Haida Gwaii populations. Finally, we used SnpEff v3.4 [93] to obtain the functional annotations of the putatively deleterious variants. The annotations were comprised of missense variants, non-coding transcripts, 3’ and 5’ untranslated regions, 5kb up- and down-stream variants, intergenic and intronic, splice acceptor and splice region, stop gained and start loss. We found 35% of deleterious mutations to be missense variants, 38% to be non-coding transcripts and 22% to be either upstream or downstream gene variants, with the remaining being spread over the other categories.

### Ethic Statement

A permit number SIRUL 111722 was obtained to work on DNA sequences. (i) No specific guidelines were followed as we did not work with animals per se and obtained all of our samples from a tissue/DNA repository of the department of Fisheries and oceans Canada; (ii) the study was approved by the following committee “Comité de protection des animaux de l’Université Laval (CPAUL)= (iii) the approval number “SIRUL 111722= was the project label at University Laval. Following answering the CPAUL questionnaire, we did not get a permit because we did not need one since there were no animals manipulations involved. Raw data will be deposit on NCBI together with Short Read Archive (SRA) accession number. All relevant vcf and jSFS files will be deposited on dryad.

## Supporting information

SuppFig

SupTab

## Acknowledgements

We thank B. Bougas, A. Perrault-Payette, C. Hernandez for laboratory support and K. Wellband, H. Cayuela, F. Hartmann for their constructive comments. TL and QR thank Benoit Nabholz for discussions regarding the deleterious mutation loads and Hill-Robertson effects. We want to thank three reviewers (Christelle Fraïsse & two anonymous reviewer) and Associate Editors for their thoughtful comments and efforts towards improving a previous version of our manuscript. Computations were mainly performed on Colosse (Calcul Quebec), Graham and Cedar (Compute Canada) servers. TL is also grateful to the Genotoul Bioinformatics Platform Toulouse Midi-Pyrenees (Bioinfo Genotoul) and the Biogenouest BiRD core facility (Universit é de Nantes) for providing computing and storage resources. This research was carried out in conjunction with EPIC4 (Enhanced Production in Coho: Culture, Community, Catch), a project supported by the government of Canada through Genome Canada, Genome British Columbia, and Genome Quebec. The authors declare no conflicts of interest.

## Supporting Information Legend

**S1 Fig. Linear decrease in genetic diversity when considering πSNP as a function of the distance to the southernmost sample site.** Each points represents a sample site and is colored by region.

**S2 Fig. Network of shared and private polymorphisms.** The branch (grey) represent shared polymorphism between sample site and are proportional to levels of sharing. Each point represents the number of private polymorphisms and is colored by region. Computation were based on a sample of size 100 in each region to enable comparison. Regional groups were chosen based on the literature regarding expected ancestral refugia.

**S3 Fig:** Violin plot of singleton distribution averaged over each region after correcting for differences in sample size. Shown is the distribution observed across 200 dataset obtained by randomly sampling individuals across populations. Black dots with errors bars represent the mean ±1 standard deviation.

**S4 Fig. Patterns of Isolation By Distance.** Increasing *F*_ST_ as a function of the distance to the southernmost site. Each point represents a sample site and is coloured by region. The *F*_ST_ was computed between the southernmost site and all other remaining sites.

S5 Fig. Summaries of *F*_ST_ values.

A. *F*_ST_-based Hierarchical tree depicting relationship among samples. Colors represent the major region.

B. Heatmap of *F*_ST_ values among samples ordered from North to South on the X and Y-axis.

**S6 Fig. Principal Component Analysis recapitulating the relationship among individuals. The Axis 3 and axis 4 are displayed.**

**S7 Fig. Multidimensional Scaling (MDS) plot depicting relationship among individuals**. Each point represents an individual site and is colored by region.

**S8 Fig. Structure and Admixture inferences.** A. Admixture Barplot obtained from LEA for various K-values. B. Progressive decrease of LEA cross-entropy criterion. Lower cross-entropy values indicates the number of cluster compatible with the data (here from 30 to 60).

**S9 Fig. Treemix results.** A. Proportion of variance explained (y-axis) as a function of the number of migration edge (x-axis) B. Treemix tree inferred without gene flow and residuals. C. Residuals for Treemix tree with four migration edges

**S10 Fig. PCA representation of the major groups used in the demographic inferences.** The site indicated as “blacklisted” are site that were not included in the ∂a∂i analyses. These corresponds to potentially admixed site between BC and Alaska and display a reduced number of individuals to constitute a coherent unit for demographic comparison.

**S11 Fig. Compared Demographic Models**

Strict Isolation (SI), Isolation with constant Migration (IM), Ancient Migration (AM) and Secondary Contact (SC). The models shared the following parameters: Tsplit: number of generation of divergence (backwards in time). *Nanc, N_1_, N_2_*: effective population size of the ancestral population, of the first and second daughter population. *M_1_* and *M_2_* represent the effective migration rates per generation that is (M = 2.Nref.m) with *m* the proportion of population made of migrants from the other population and Nref the size of the reference population. T*sc* is the number of generations since gene flow started (secondary contact) after a period of isolation. T*am* is the number of generations since the two populations have diverged without gene flow until present. Each model is declined in alternative version allowing homogeneous or heterogeneous effective size and homogeneous or heterogeneous gene flow to account for the effect of linked selection (affecting *Ne)* and barrier to gene flow (affecting *m*) respective.

**S12 Fig. Models and residuals for each best model**

Each plot displays the observed jSFS (data), the modeled jSFS (model) and the residuals. Left part: Best model without population size change. Each best model was inferred using ΔAIC and AIC weights. Right panel: The same model as the left but including the possibility for population size change of the diverging daughter populations.

**S13 Fig: Global model tested with Fastsimcoal.**(see details in Supplementary Note S1)

**S15 Fig: Summary of admixture coefficient obtained from Fig6. F** or easier interpretations of admixture coefficient among the 58 samples site a PCA was performed to summarized the distribution of admixture. Populations with lower cos2 contributed weakly to the plot and hence have higher admixture.

**S15 Fig: Relationship between Tajima’s D and distance to the southernmost site.** No significant relationship was observed (p=0.5, r=−0.07), suggesting that each local population has undergone different evolutionary trajectory in post-glacial time.

**S16 Fig**: **DAF spectrum of synonymous, non-synonymous and putatively deleterious mutation aggregated at the regional level for all samples.** Data are normalized for a sample of size n = 102 corresponding to the smallest size for the combined samples in Haida Gwaii.

**S17 Fig. DAF spectrum of synonymous, non-synonymous and putatively deleterious mutation in each locality for all samples.**

**S18 Fig: A) Distribution of the count of homozygous derived deleterious alleles in each major group. B) Distribution of the count of total derived deleterious alleles in each major group.**

**S19 Fig. Mean derived allele frequencies distribution of deleterious mutation.** Displayed are the mean derived allele frequencies of polymorphic deleterious sites in each region +/− 2 standard deviation.

**S20 Fig. Correlation between π_N_/ π_S_ and π_S_ used as a proxy for long term effective population size of each local population.**

## Table Legend

**S1 Table:** Abbreviation, Region and coordinates (Longitude and Latitude) of each river used in the GBS data with the number of individuals provided (nb. Inds).

**S2 Table:**β_ST_ values along with 95% confidence intervals for each river from the GBS data (82 K SNPs). 95% confidence intervals obtained after 1000 bootstraps.

**S3 Table:** model choice results for dadi. AIC, ΔAIC and AIC weights are provided for each pairwise comparison and model. AM = Ancient Migration, IM = Isolation with Migration, SI = Strict Isolation, SC = Secondary Contact, the simplest models assume homogeneous migration and homogeneous effective population size. 2N suffix = heterogeneous effective population size, 2M suffix = heterogeneous migration. Model with both suffix assumes that both effective population size and migration are heterogeneous. Model with a single suffix assumes that either migration or effective population size are heterogeneous. In one cases (Calif2 vs Thompson) the SC2N, SC2m and SC2N2m provided similar fit. For consistency the SC2N2m was considered for parameter estimates. Its parameter estimates were more consistent than those from either SC2N or SC2m.

**S4 Table:** Parameter estimates obtained under the best demographic model with dadi. Ne1 and Ne2, effective population size of the compared pair. m1 ← 2 and m2 ← 1, migration from population 2 to population 1 and migration from population 1 into population 2. me12 and me21, effective migration rate estimated in the most differentiated regions of the genome Ts: Split Time of the ancestral population in two population; Tsc: duration of the secondary contact P: proportion of the genome freely exchanged (1-P provides the proportion of the genome non-neutrally exchanged); Q: proportion of the genome with a reduced effective population size due to selection at linked sites; hrf = Hill-Robertson factor representing the reduction of Ne in the region Q with reduced Ne.

**S5 Table**: Confidence Intervals obtained from the Godambe Information Matrix (GIM).

**S6 Table:** Confidence Intervals obtained from 100 dataset constructed with the coalescent simulator ms.

**S7 Table**: Parameter estimates and confidence intervals obtained from Fastsimcoal.

**S8 Table:** Tajjima’s D value and π_N_/ π_S_ value observed for each locality.

**S9 Table:** Summary of deleterious variation by region. 1)Derived Allele Frequency (DAF) of deleterious mutation, after averaging by rivers and then by major regional group. 2) Count of deleterious mutations in each river and then averaged by major regional group. 3) Number of homozygous derived deleterious mutations by individual, after averaging by rivers and then by major regional group. 4) Number of heterozygous mutations by individuals, after averaging by rivers and then by major regional group 5) Total load of derived deleterious mutations by individuals, after averaging by rivers and then by major regional group.

**S10 Table:** Results of Wilcoxon test (Mann-Whitney tests) for differences in derived allele frequencies among major groups for a sample of size 100.

**S11 Table:** Results of Wilcoxon test (Mann-Whitney tests) for differences in count of derived homozygous variants and total load among individuals in each region.

**Supplementary Note S1:** Details of Fastsimcoal analyses to infer a global scenario of divergence. The tested model is displayed in Fig7 and the S7 Table provide details of parameter estimates.

