## Supplementary material for "Demographic history shaped geographical patterns of deleterious mutation load in a broadly distributed Pacific Salmon": SuppFig: FigS2.pdf

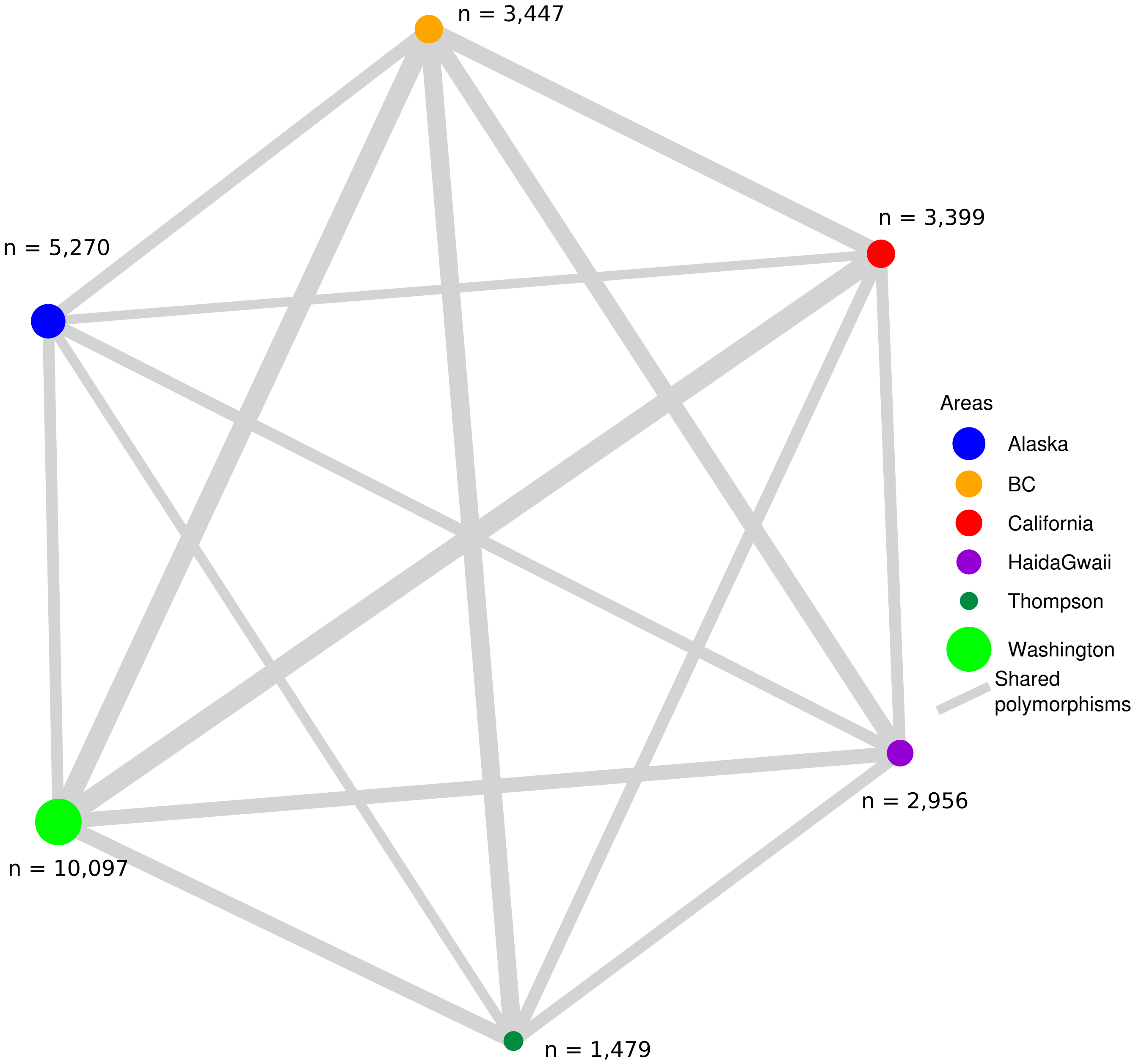


Levels of shared polymorphisms based on a sample of size = 100

| | Alaska | BC | California | HaiaGwai | Thompson | Cascadia |
|---|---|---|---|---|---|---|
| Alaska | –– | 6781 | 4874 | 6467 | 5215 | 6036 |
| BC | 6781 | –– | 8621 | 8699 | 9484 | 10346 |
| California | 4874 | 8621 | –– | 6160 | 6613 | 10629 |
| HaidaGwai | 6467 | 8699 | 6160 | –– | 6447 | 7657 |
| Thompson | 5215 | 9484 | 6613 | 6447 | –– | 7830 |
| Cascadia | 6036 | 10346 | 10629 | 7657 | 7830 | –– |
