## Supplementary material for "Demographic history shaped geographical patterns of deleterious mutation load in a broadly distributed Pacific Salmon": SuppFig: FigS12.pdf

***CALIFORNIA1 vs ALASKA***

***best: SC2N2m*** ***AIC = 25,572***

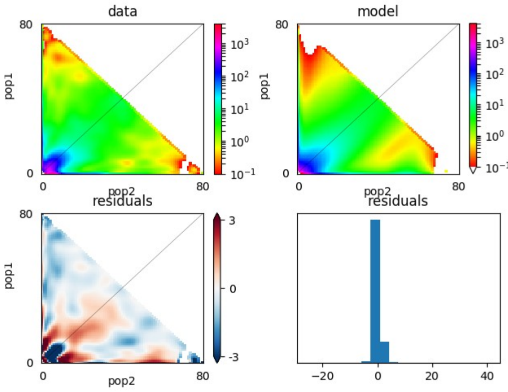


***SC2N2mG*** ***AIC = 21,943***

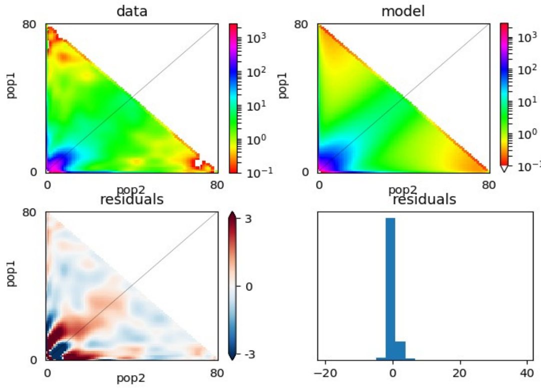


***CALIFORNIA2 vs ALASKA***

***best: IM2N2m*** ***AIC = 6,433***

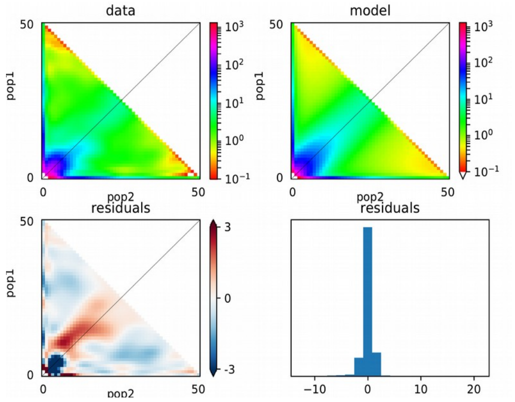


***IM2N2mG*** ***AIC = 6,317***

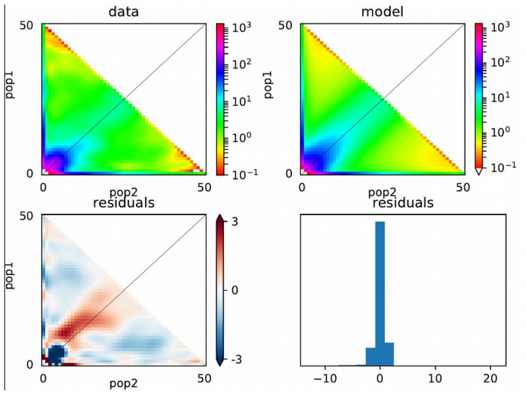


***CASCADIA vs ALASKA***

***best: IM2N2m*** ***AIC =18,061***

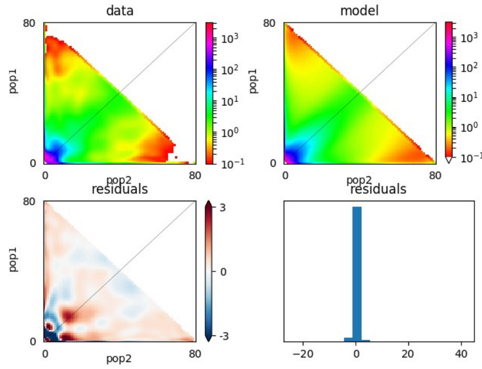


***IM2N2mG*** ***AIC = 17,346***

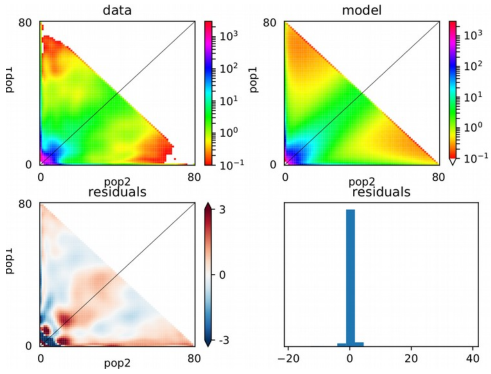

***BC vs ALAKSA***

***best: IM2N2m*** ***AIC = 19,844*** ***IM2N2mG*** ***AIC = 18,703***

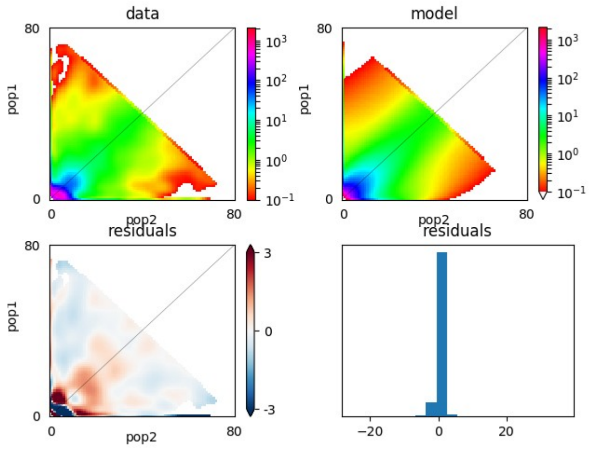


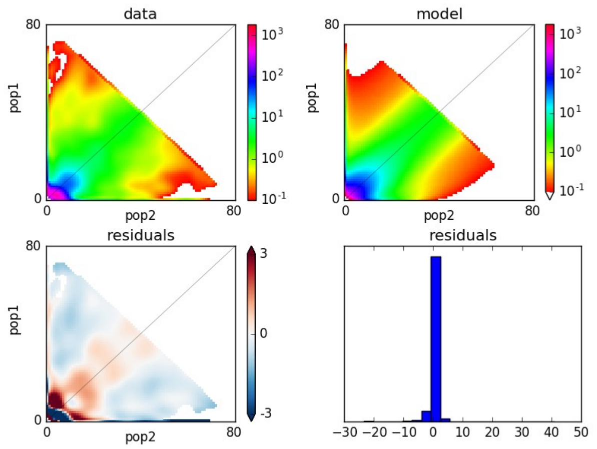


***THOMPSON vs ALASKA***

***best: SC2N2m*** ***AIC = 33,229*** ***SC2N2mG*** ***AIC = 25,825***

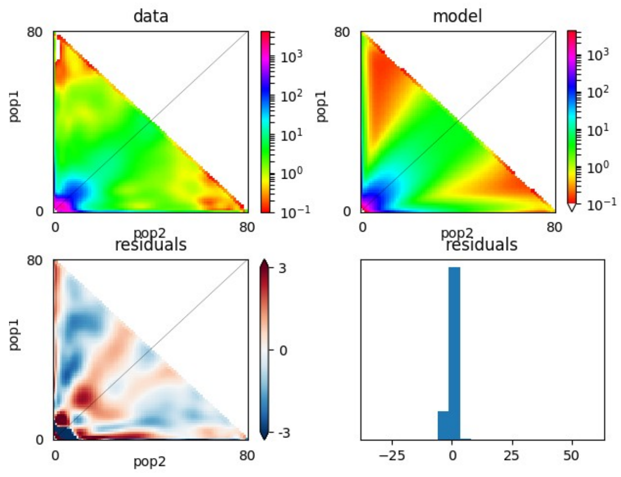


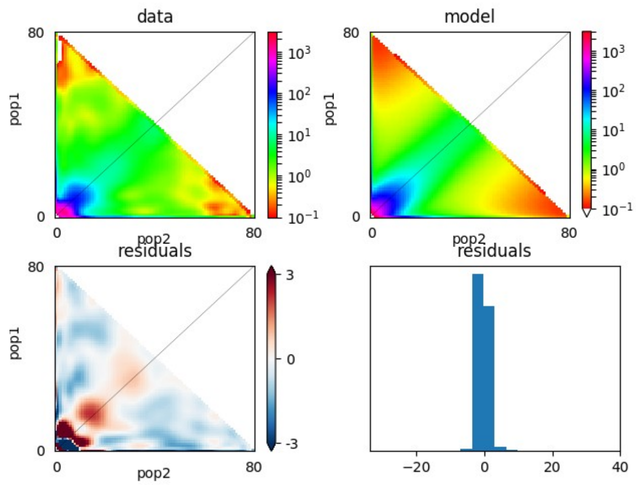

***CALIFORNIA1 vs THOMPSON :***

***best: SC2N2m*** ***AIC = 26,987***

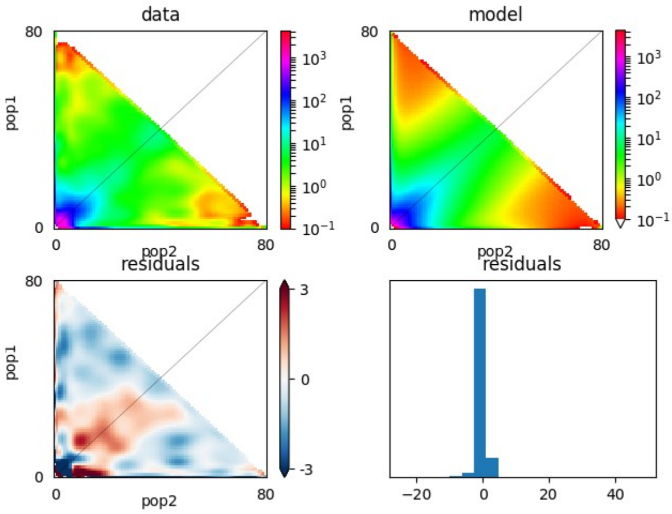


***SC2N2mG*** ***AIC = 22,354***

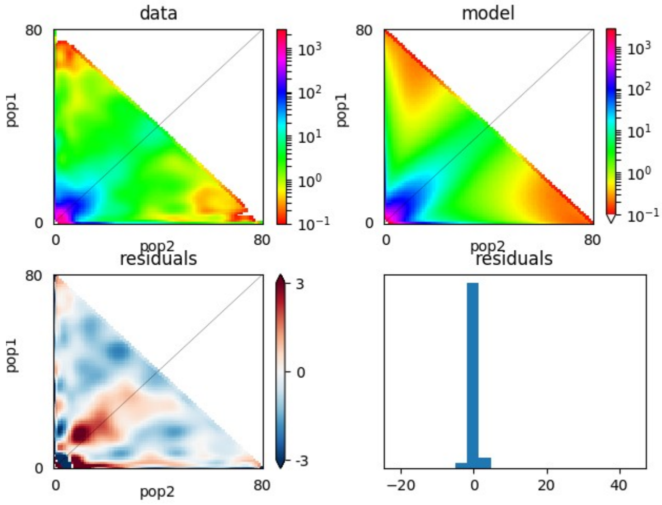


***CALIFORNIA2 vs THOMPSON :***

**SC2N2mG** **AIC = 8298**

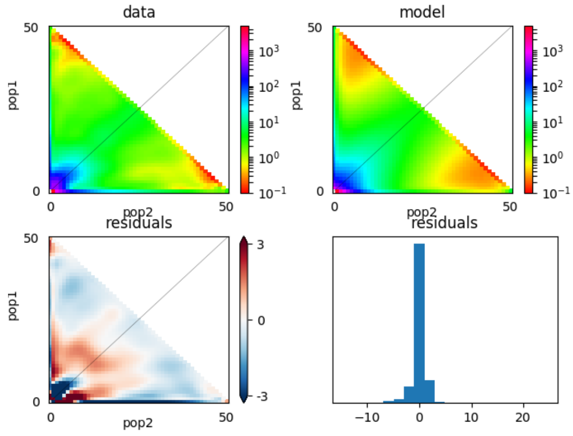


**SC2N2mG** **AIC = 7,367**

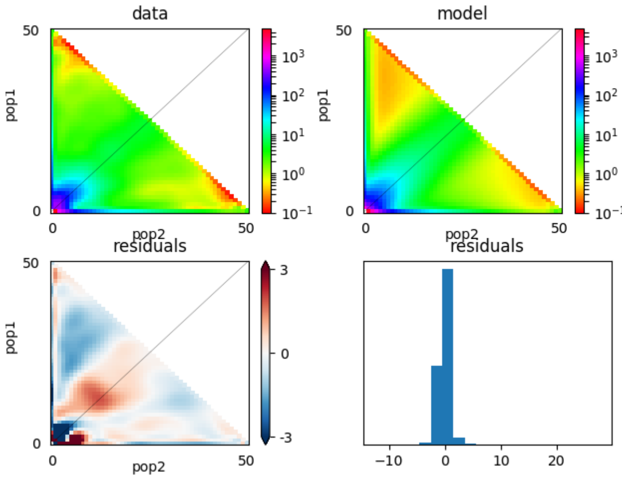

***CASCADIA vs THOMPSON***

***SC2N2m*** ***AIC = 45,178***

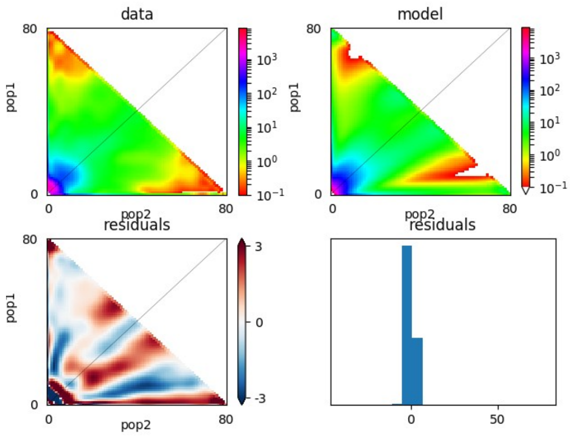


***SC2N2mG*** ***AIC = 39,394***

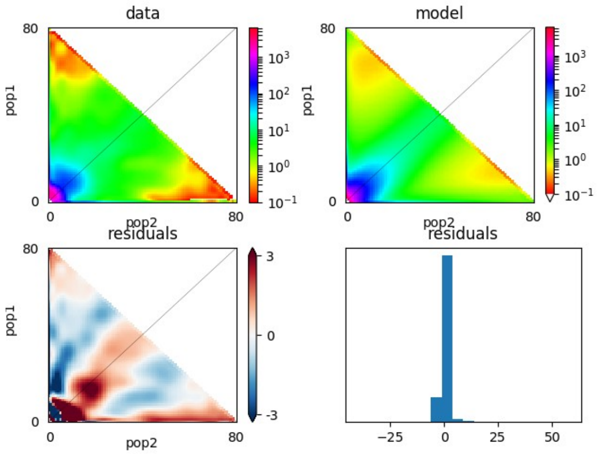


***BC vs Thompson***

***SC2N2m*** ***AIC = 48,069***

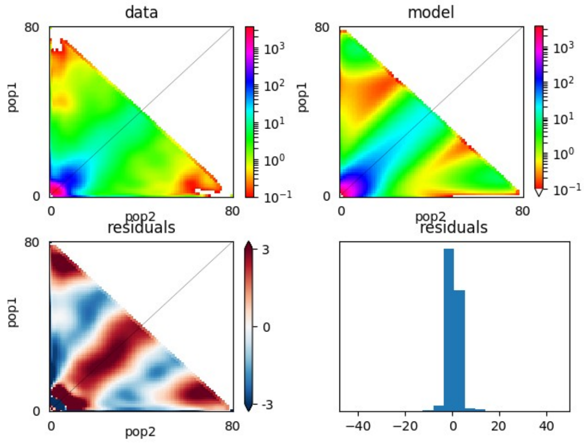


***SC2N2mG*** ***AIC = 39,270***

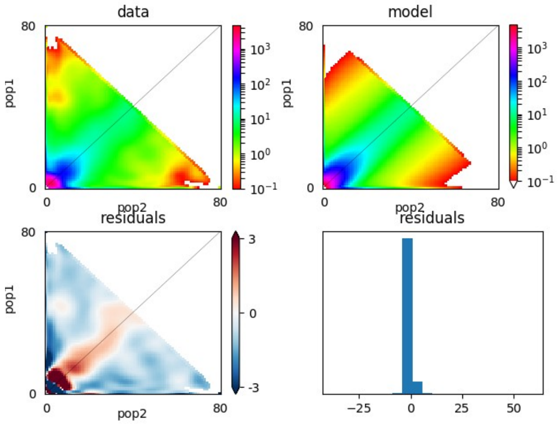
