## Supplementary figures and images for "Demographic history shaped geographical patterns of deleterious mutation load in a broadly distributed Pacific Salmon"

### FigS1.pdf

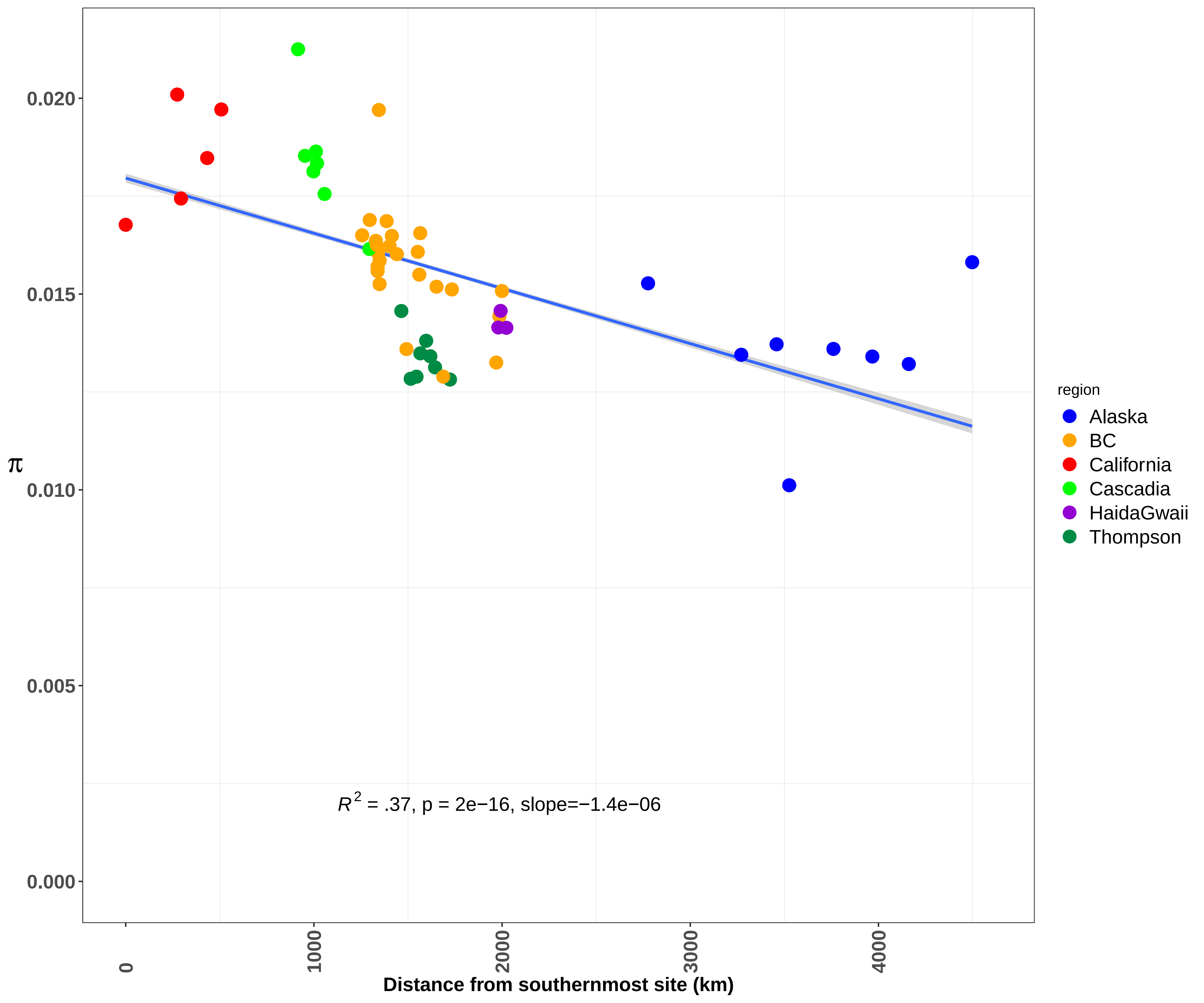

### FigS3.pdf

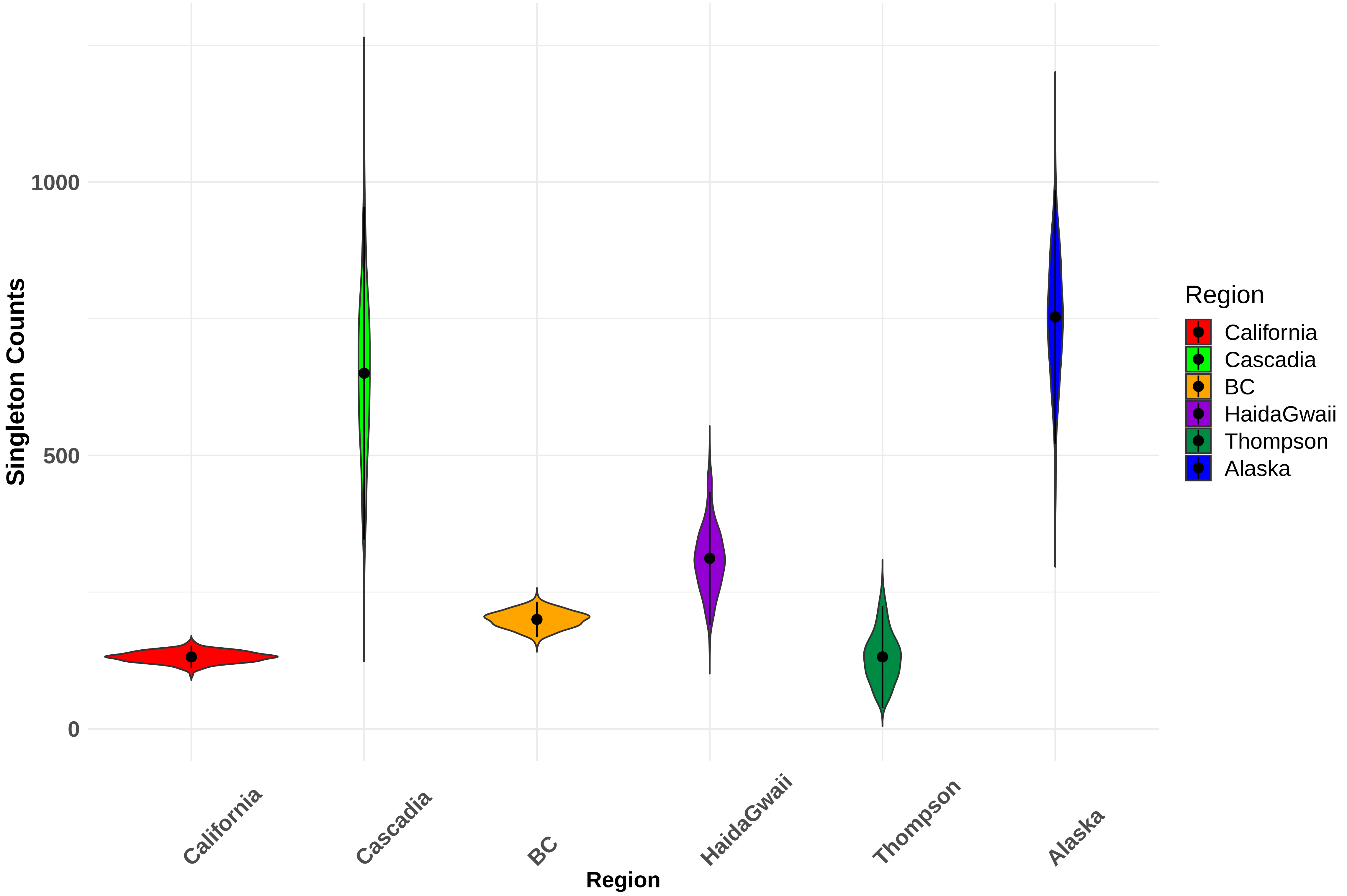

### FigS4.pdf

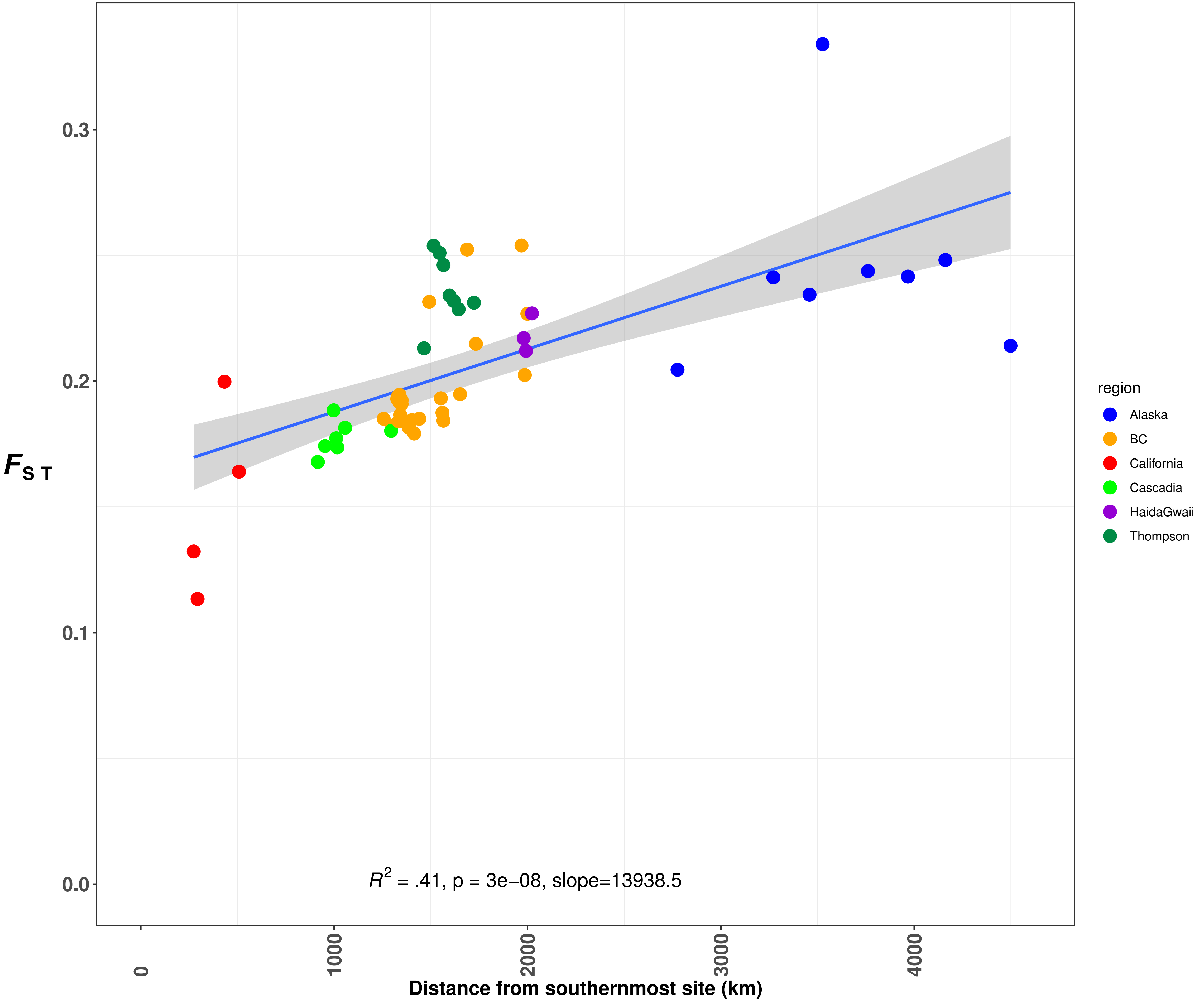

### FigS5.pdf

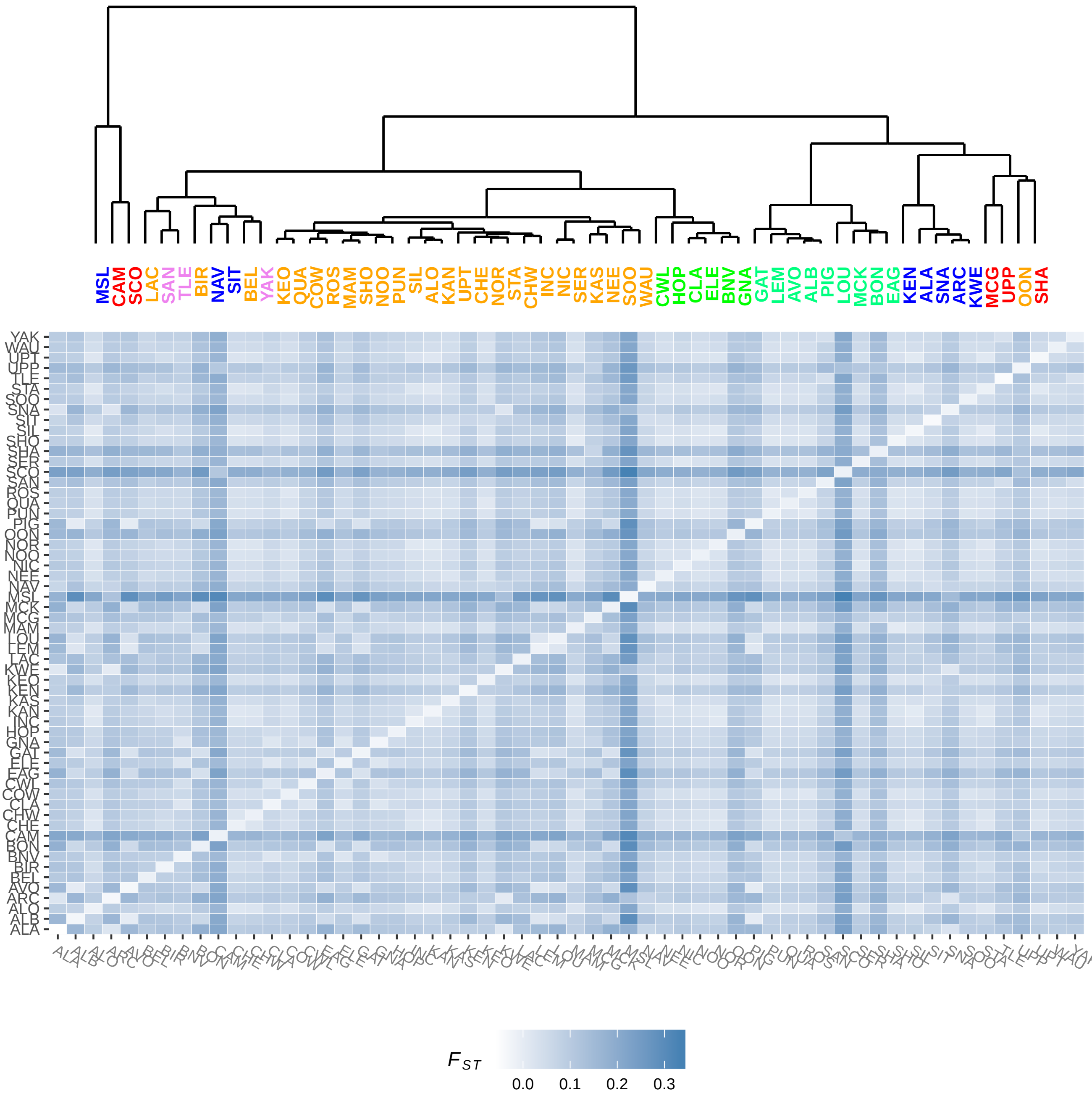

### FigS6.pdf

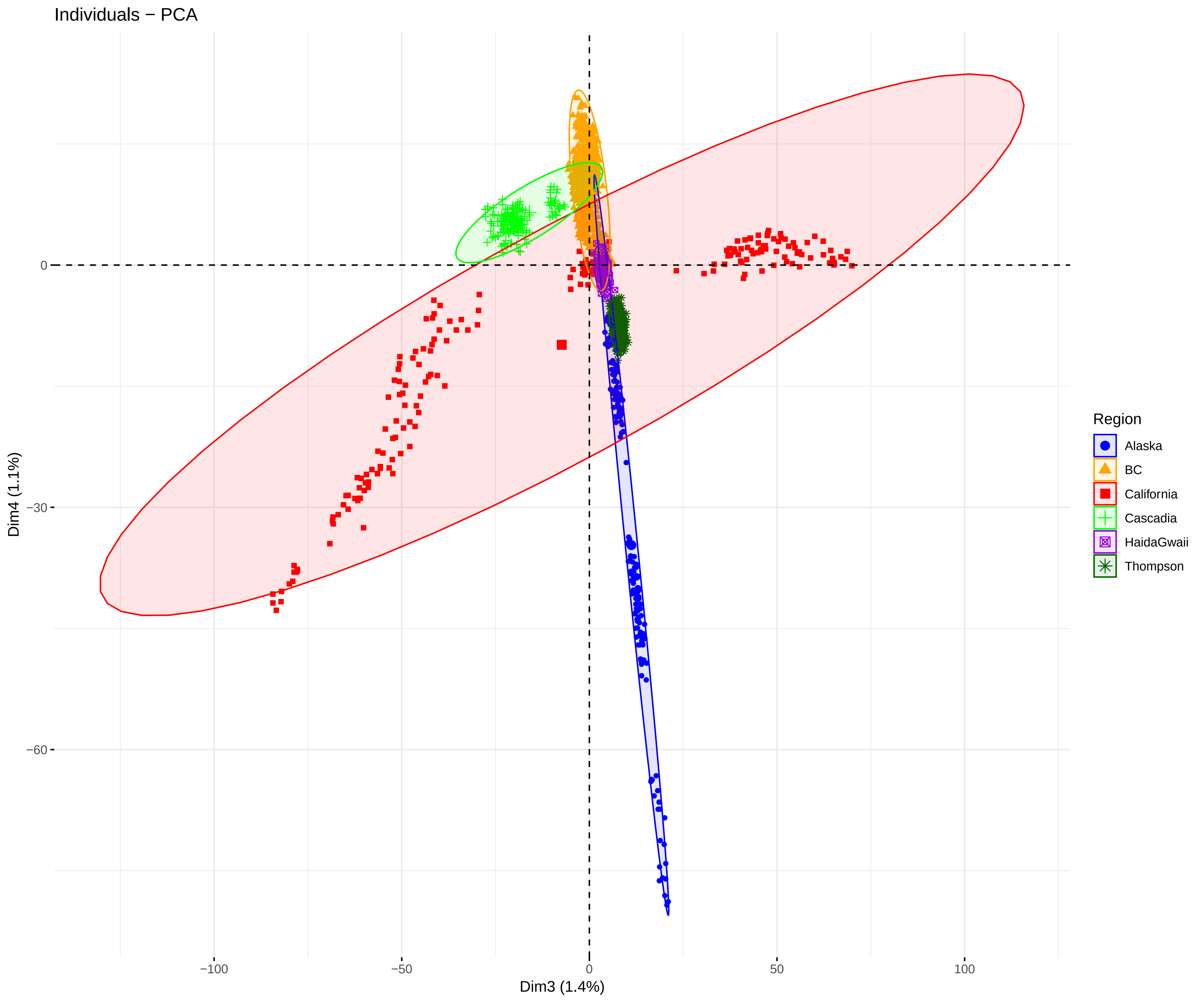

### FigS7.pdf

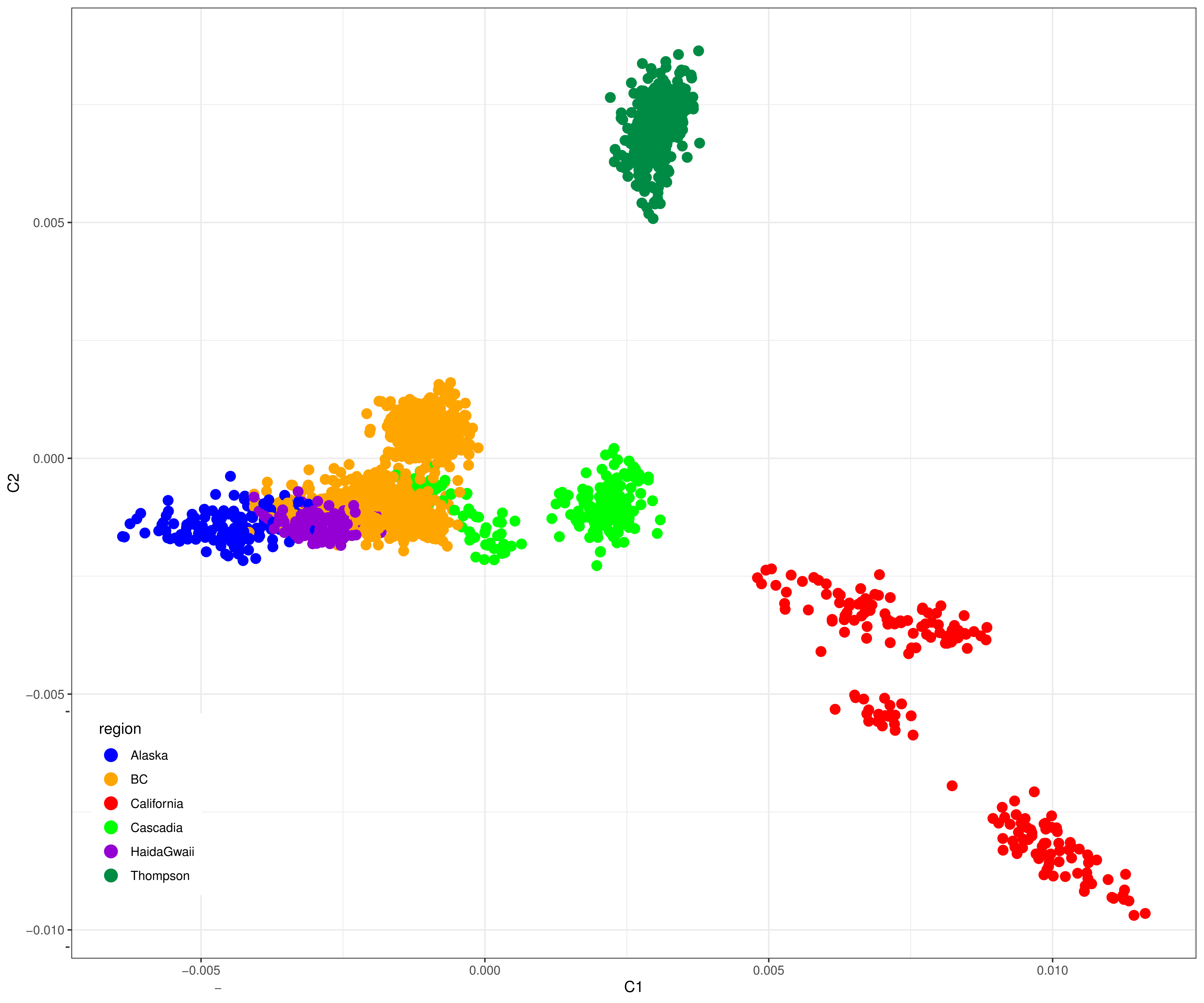

### FigS8.pdf

A. Admixture proportions

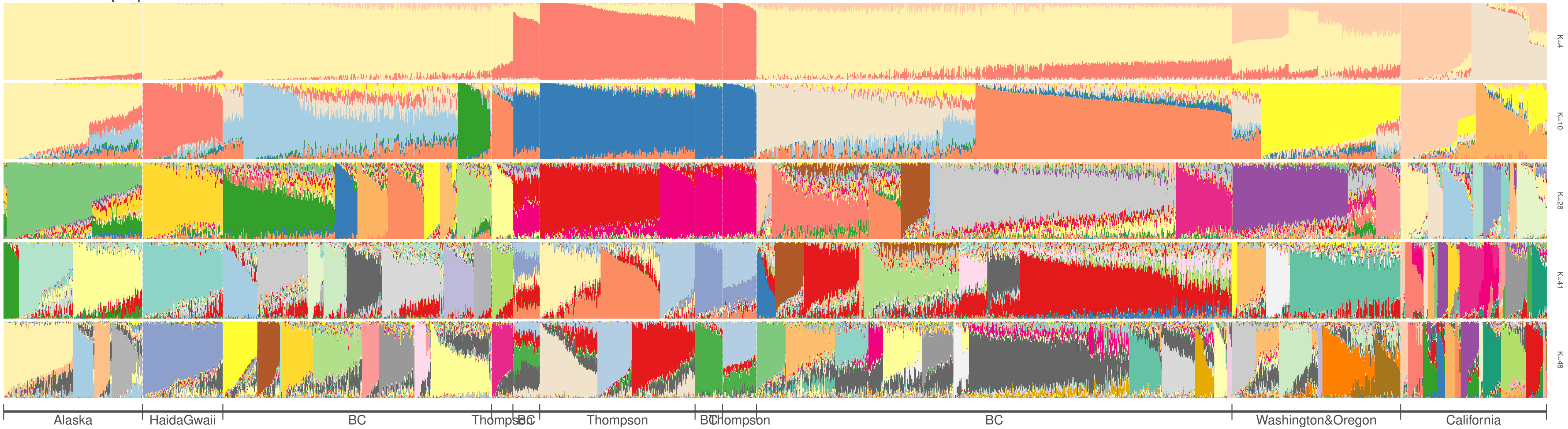


B. Cross entropy as a function of the number of cluster

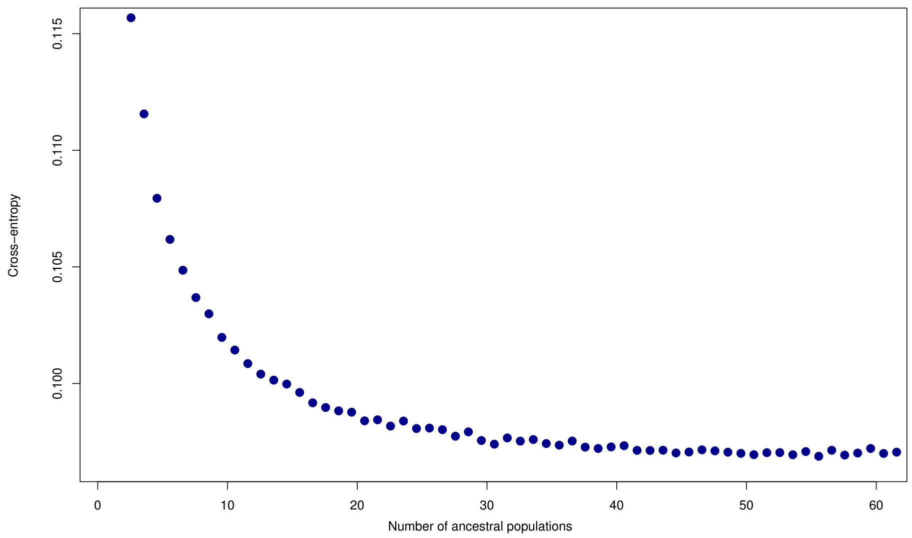

### FigS9.pdf

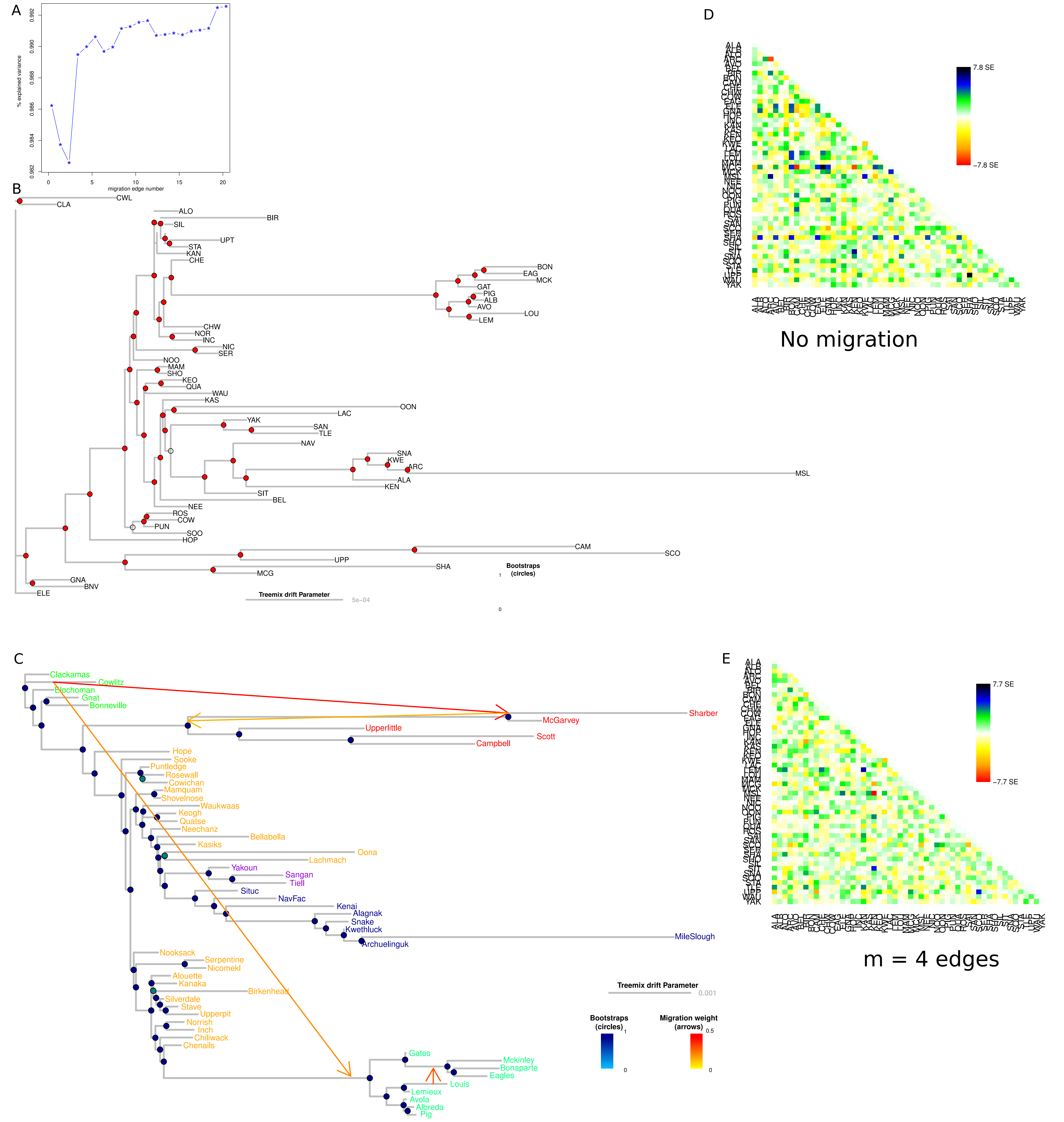

### FigS10.pdf

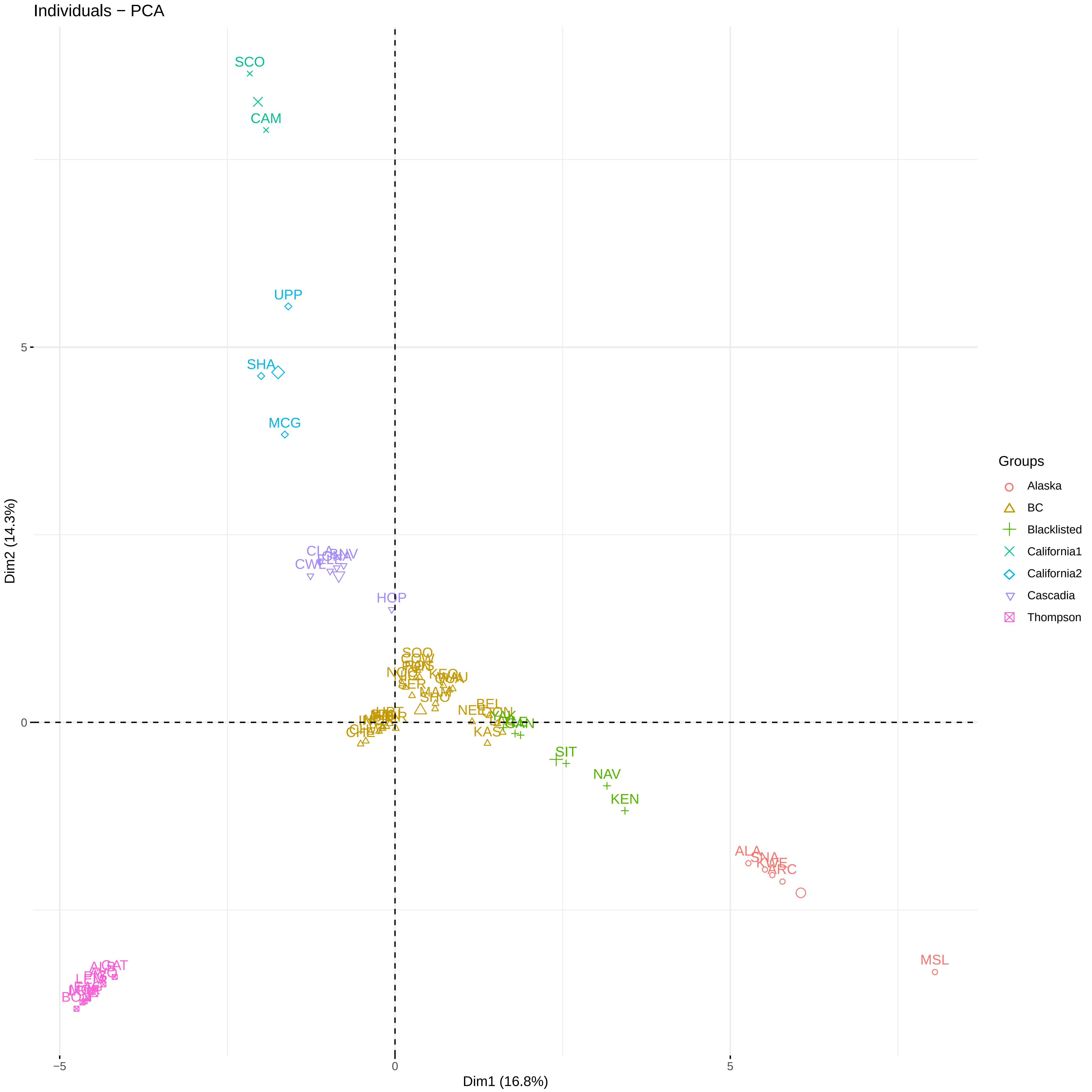

### FigS11.pdf

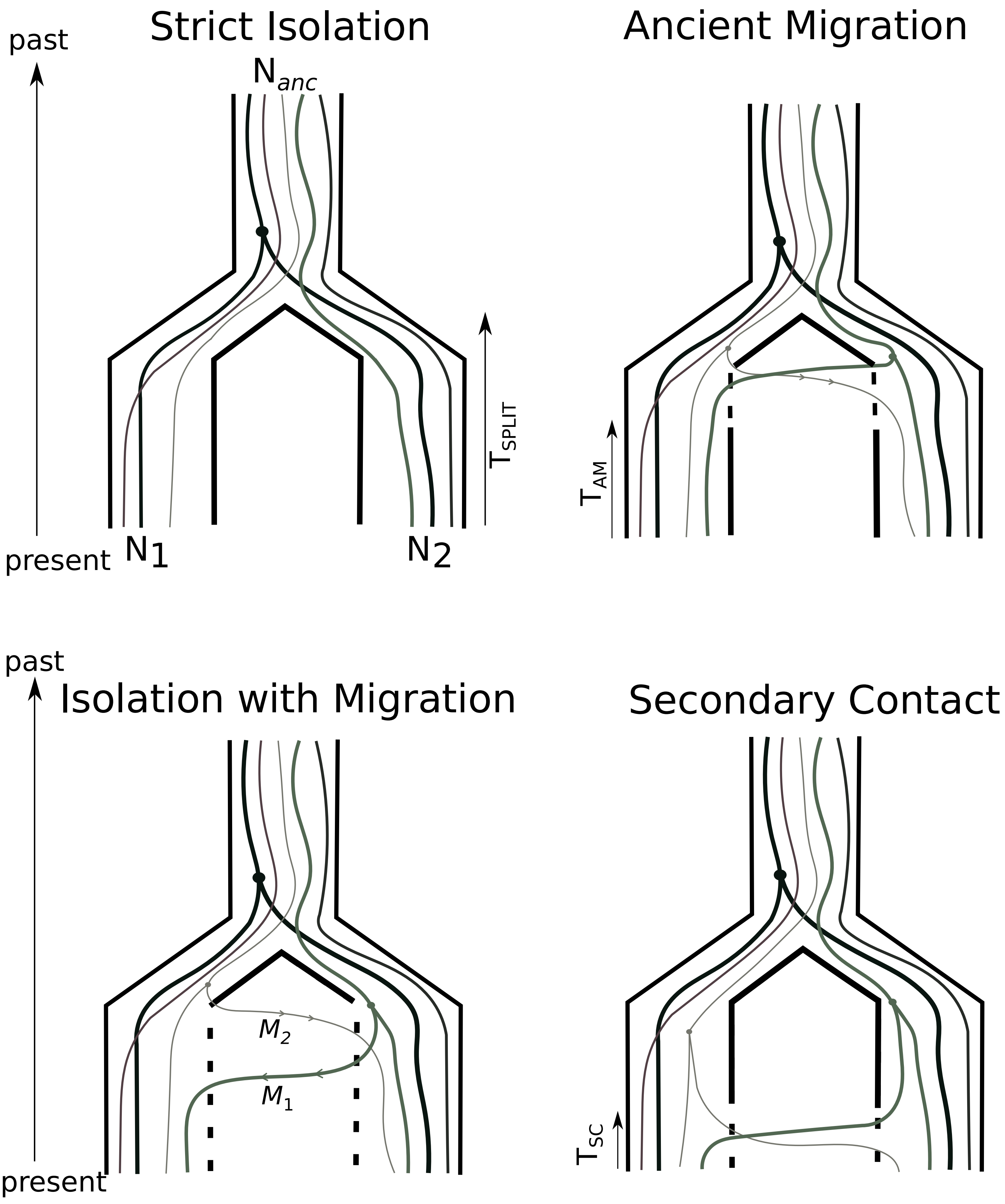

### FigS13.pdf

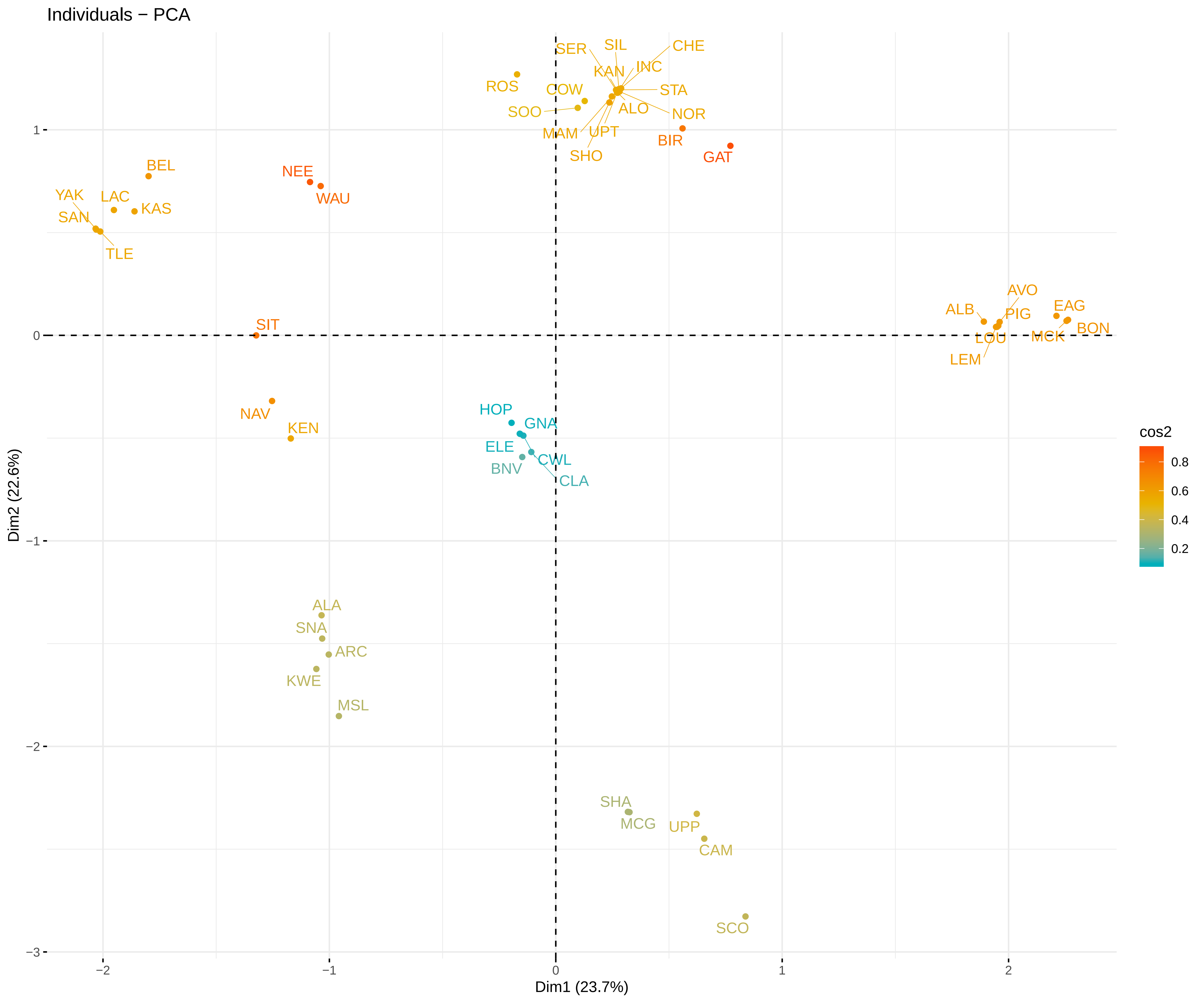

### FigS14.pdf

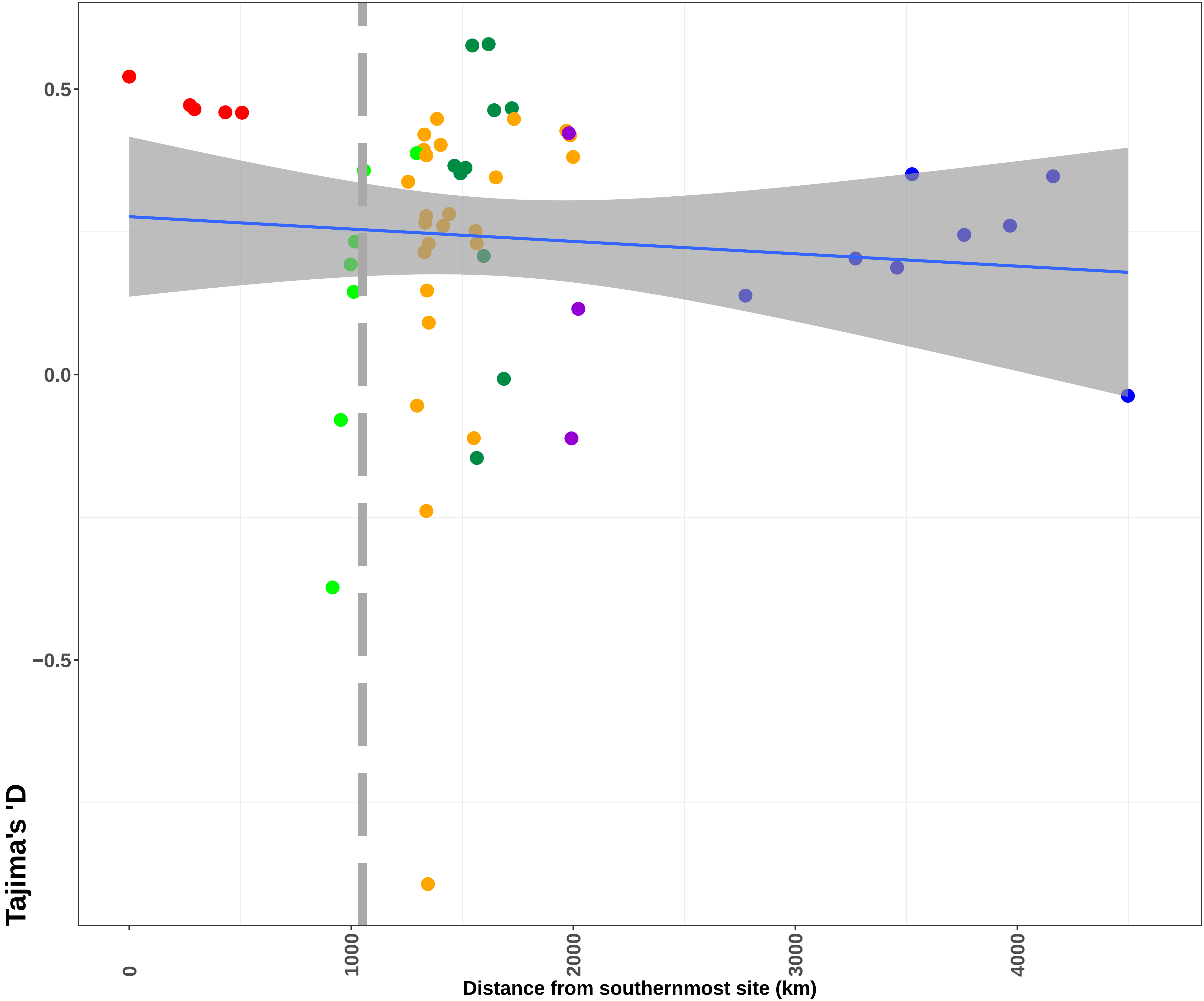

### FigS15.pdf

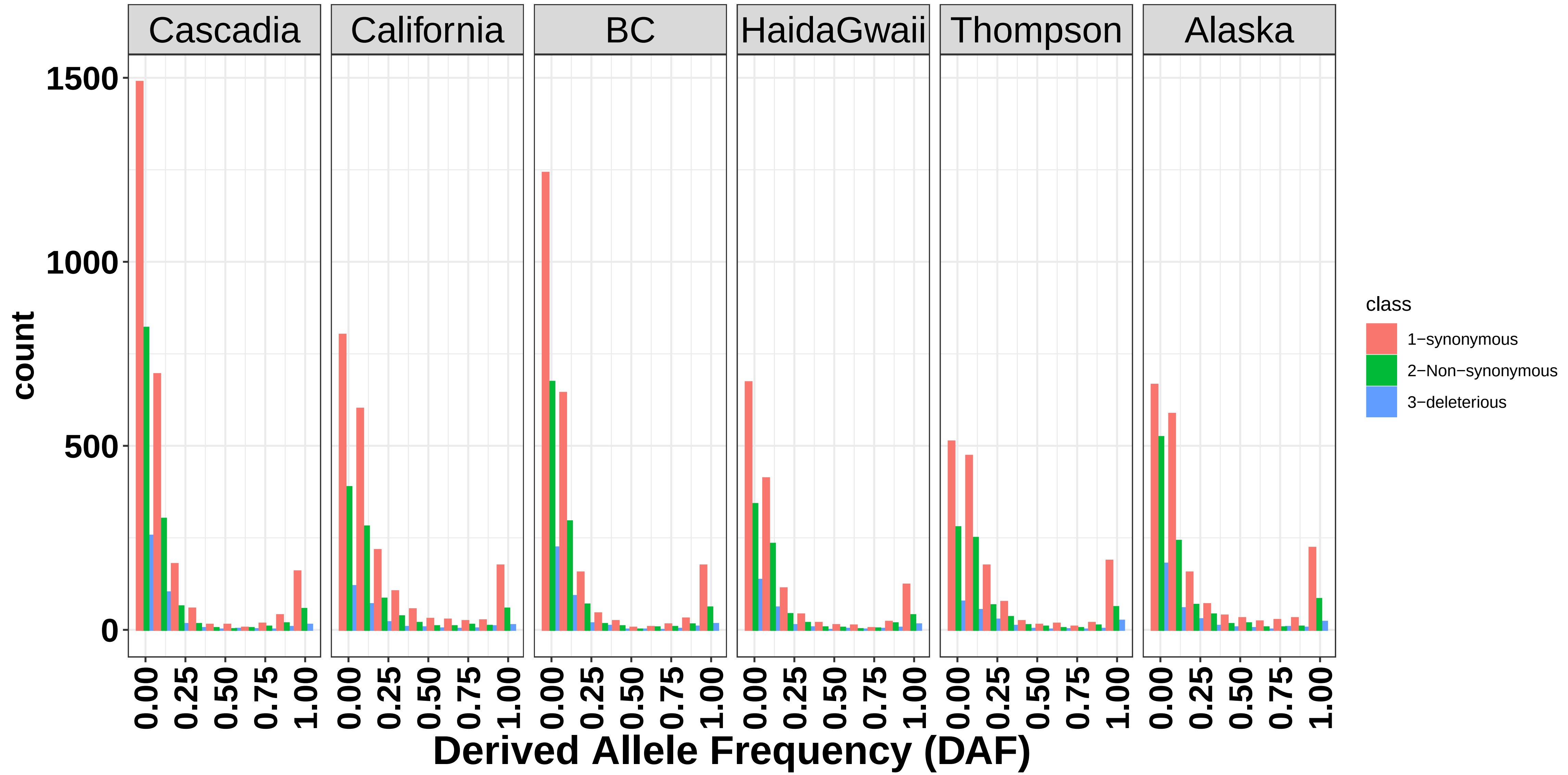

### FigS16.pdf

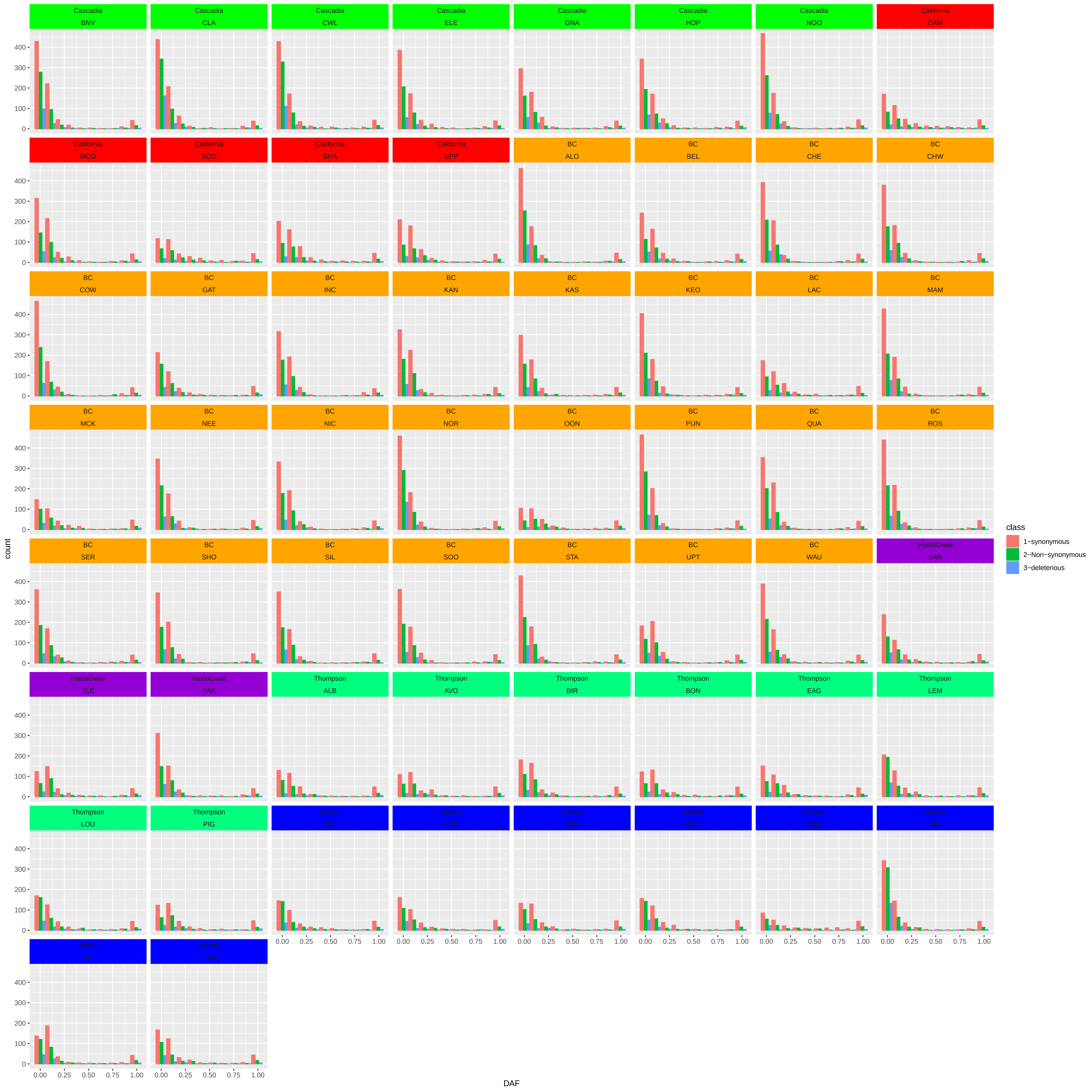

### FigS17.pdf

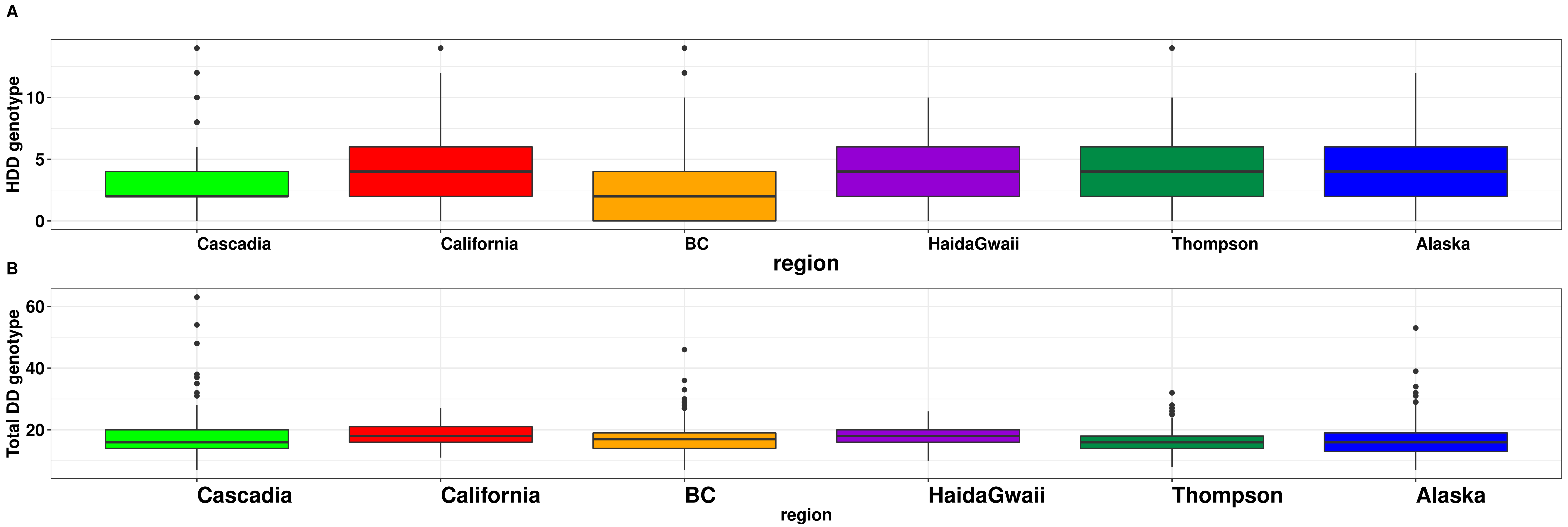

### FigS18.pdf

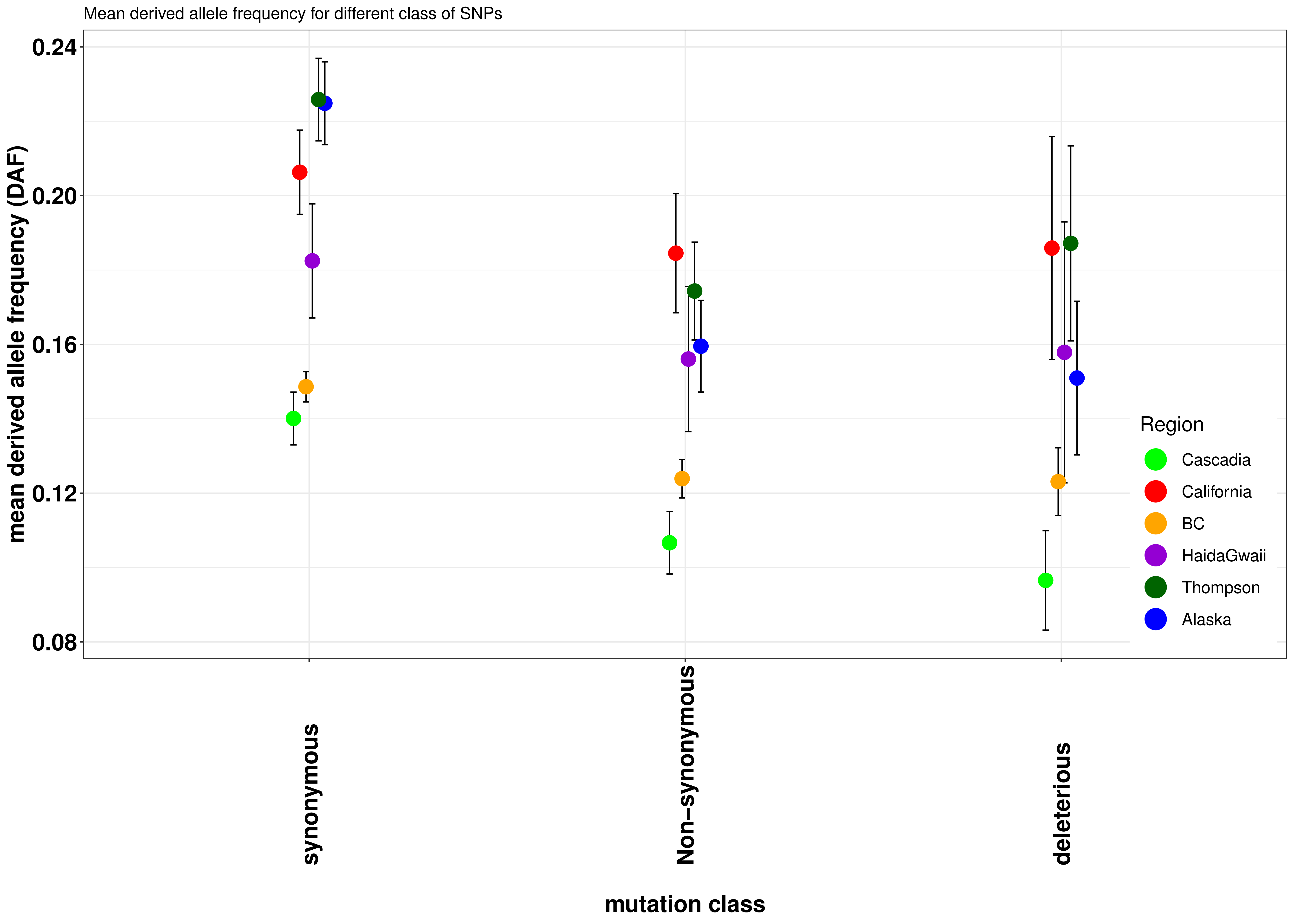

### FigS19.pdf

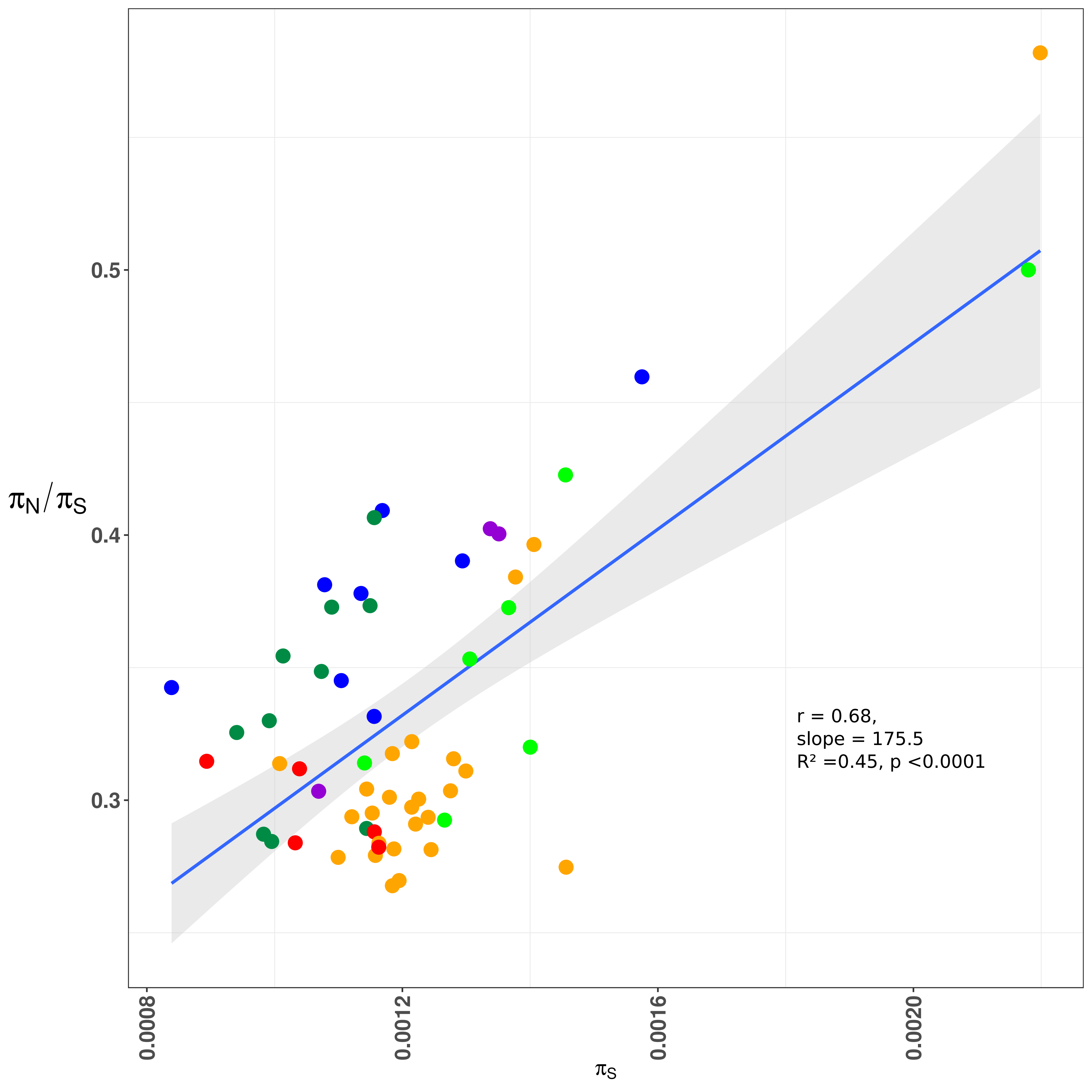
