## Supplemental Note for "Demographic history shaped geographical patterns of deleterious mutation load in a broadly distributed Pacific Salmon"

**Supplementary Note S1:** ***Inferring a global divergence history***

***Context:***

Ideally, a direct comparison of a single versus two refugia model would be needed to disentangle among a single versus two refugia scenarios. However, a direct comparison based on the site frequency spectrum of one versus two populations is not possible. Therefore, the comparison of two-dimension site frequency spectrum, while imperfect, still allows circumventing these limitations and provided a first overview of the global demographic history. To refine our understanding of the demographic history we also tested a global model of divergence with Fastsimcoal v2.6 [1] and our observed sites frequency spectrum by constructing a model including all six populations simultaneously. Unlike our ∂a∂i procedure, Fastimcoal does not accounts for the confounding effects of the linked selection or barriers to gene flow. To limit the confounding by barriers to gene-flow [2], linked selection [3,4] or gBGC [5], we restricted our jSFS to areas of high recombination, as inferred by LDHAT [6] from whole genome sequences (unpublished data). Our goals here were to 1) test the topological order of population divergence (i.e. verify if Cascadia or California diverged first and were most ancestral) and 2) refine parameter estimates of this model, in particular refining the splitting time of the population and testing for population expansion or reduction during their divergence.

***Methods**:*

Ideally, model choice based on AIC should be performed with strictly independent SNPs and directly from the multi-SFS. Moreover, very large number of SNPs are required for using the multi-dimensional SFS [1]. Here, we only constructed multiple joint site-frequency spectrum. Therefore, a simple approach was undertaken: first we seek to verify the topological order of population divergence. Namely we wanted to address the question of whether California or Cascadia diverged first. To do so, we compared two alternative topologies. In the first topology, California diverged first and populations followed a strict south to north expansion (California1 – California2 – Cascadia – BC – Thompson – Alaska). In the second topology, Cascadia diverged first, followed by California (Cascadia – California2 – California1 – BC – Thompson – Alaska). Due to an exponential increase in computational time of this 6 population model, these comparisons were performed with gene flow. Second, we used the best topology and included gene flow in this model to infer the the parameter of a global model of divergence, based on the observed scenario in ∂a∂i. We constructed a model of population expansion from the South to the North, we modeled multiple founder event and allowing for population size change (growth or decline). Population diverged hierarchically with a decreasing age of divergence as one move Northward (Fig 7).

All populations evolved in strict isolation and were allowed to grow or decay independently following their divergence. Prior on all demographic parameter were uniform and were set according to the range of divergence time, migration rate and effective size observed in ∂a∂i. multiple-joint frequency spectrum were obtained from ANGSD by masking regions of low recombination. Unfolded jSFS were constructed using the three outgroups (Chinook salmon, Rainbow trout and Atlantic salmon) to reconstruct the ancestral sequence. Details of the models and script can be found on Git hub at https://github.com/QuentinRougemont/fsc_modeling

To perform a model comparison of the best topological order, we ran 100 independents replicates, with 50,000 coalescent simulations per likelihood estimates and 20 cycles of the likelihood maximization algorithm. All models assumed a finite site mutation model. Model choice was performed by computing AIC of each model.

After choosing the best topology, we constructed the final model for parameter estimates. Here,all populations were connected by ongoing and asymmetric migration and were allowed to grow or decay independently following their divergence. After the split of Thompson from the rest of the main distribution the population evolved without gene-flow until post-glacial time were a secondary contact occurs.

To obtained the maximum likelihood estimates of the demographic parameters under this model, we performed 50 independent runs, with 50,000 coalescent simulations per likelihood estimates and 20 cycles of the likelihood maximization algorithm. All models assumed a finite site mutation model. Finally, we derived confidence intervals using 100 bootstrap replicates, and 5 independent runs in each bootstrap. The inclusion of migration within our models considerably increases the computational time, preventing the use of more replicates for each bootstrapped replicates. The code to reproduce the results is available on git hub at https://github.com/QuentinRougemont/fsc_modeling.

***Results*** :

**Model comparison:**

The model choice procedure clearly indicated that the Californian samples diverged first, followed by Cascadia and then northern population as opposed to a split of Cascadian samples first (AIC = 5,562,351.8 vs 5,981,900.4). This model was not used for parameter estimates given that the exclusion of gene-flow within the model is expected to bias our estimates of parameter.

**Parameter estimates:**

We ran 50 replicates that followed the topology inferred above but included migration between all populations. As expected, including gene-flow strongly outperform models of isolation (AIC = 4,350,420). Our parameter estimates based on the best run out of 50 independent replicates are provided in the **S7 table** below. The model provided highly informative and complementary results to our ∂a∂i results. In particular, according to **S7. Table,** we observed that the population from California (1 & 2) and Cascadia were those exhibiting highest ancestral size. This was also true for the Alaskan samples. The Californian population was inferred to have undergone strong population contraction, whereas the Thompson and Alaskan samples exhibited modest size reductions. The size of the BC samples was highly reduced at the onset of divergence and then increased to a larger size. All the populations were inferred to have diverged in a very narrow time window ~ 120 – 95 KyA, a result slightly more recent than the inferred divergence time with ∂a∂i. Although the confidence intervals were narrower with Fastsimcoal they still overlap due to larger uncertainty of the ∂a∂i estimates. The time of secondary contact between Thompson and the main distribution was also inferred to have occurred very recently (~10KyA), in line with those observed in ∂a∂i. Finally, in all our comparisons we note that migration rate were inferred to be low and asymmetric.

***Discussion:***

Our results provide an overview of the global divergence history of the populations that are complementary to ∂a∂i (see also **Fig 7**). Importantly, we were able to formally test the topology and provided evidence that southernmost populations are most ancestral. Then, populations have quickly expanded North and the series of funding events resulted in variable change in effective population size. Estimates of population split time were very narrow. This suggests that populations have expanded Northward in a very narrow window of time and quickly colonized new areas where each local population start to diverge in different rivers. Admittedly, while the model comparison indicated that Californian samples were likely most ancestral, it is very likely that both California and Cascadia individuals were segregating in a single refugia, given that the entire area was not glaciated in the past. In recent times, Californian individuals undergone strong effective population size reduction which is in line with the analyses of genetic diversity (see main text). Therefore, considering the observed genetic diversity levels (Fig 2A, S4 Fig) distribution of singletons (S3 Fig), genetic differentiation (Fig 2B, S4 Fig), PCA results, and other summary statistics, we considered the Cascadian samples with the lowest observed $\beta_{ST}$ coefficient as the most ancestral for the analyses of mutation load (**Fig 6** in main text).

In all cases, our demographic reconstruction provide increased support for our scenario of a major refugia in the South contributing most of present day genetic variation (see main text) and constitute added justification for the study of the fate of deleterious variants, under models of expansion load [6–8].

**Supplementary Table S7 parameter estimates.** (a) distribution of ancestral (Nanc) and current effective population size (Ncurrent). b) estimates of split time. Tsc = Time of secondary contact between Thompson and the main distribution group c) estimates of population growth rate (negative value indicates population expansion). d) migration rate. Value : Parameter estimates obtained for the best model. CI2.5 and CI97.5: Lower and upper bound of confidence intervals around the parameter estimates, respectively.

**(a) ancestral and current effective population size.**

| | ancestral effective population size | | | | current effective population size | | | |
|---|---|---|---|---|---|---|---|---|
| population | Nanc | value | CI2.5 | CI97.5 | Ncurrent | value | CI2.5 | CI97.5 |
| California1 | Nanc0 | 420024 | 335838 | 504210 | N0 | 33041 | 0 | 155267 |
| California2 | Nanc1 | 580442 | 518317 | 642567 | N1 | 96803 | 26264 | 167342 |
| Cascadia | Nanc2 | 785420 | 673755 | 897085 | N2 | 858447 | 766591 | 950303 |
| BC | Nanc3 | 1594 | 0 | 84254 | N3 | 281886 | 201585 | 362187 |
| Thompson | Nanc4 | 350346 | 269007 | 431685 | N4 | 285316 | 199666 | 370966 |
| Alaska | Nanc5 | 815496 | 663459 | 967533 | N5 | 750494 | 591620 | 909368 |

**b) population split time**

| | Split time | | |
|---|---|---|---|
| Time | value | CI2.5 | CI97.5 |
| T0 | 116500 | 108000 | 125000 |
| T1 | 112900 | 104300 | 121400 |
| T2 | 101200 | 92500 | 109900 |
| T3 | 98000 | 89000 | 106600 |
| T4 | 92200 | 84000 | 100500 |
| Tsc | 10986 | 10986 | 11709 |

### c) population growth rate

| | Growth Rate | | | |
|---|---|---|---|---|
| population | growth | value | CI2.5 | CI97.5 |
| California1 | G0 | 0.0020671 | 0E+00 | 2E-03 |
| California2 | G1 | 4.60E-04 | 0E+00 | 6E-04 |
| Cascadia | G2 | -2.28E-05 | 0E+00 | 1E-04 |
| BC | G3 | -4.67E-03 | 0E+00 | -4E-03 |
| Thompson | G4 | 1.10E-04 | 0E+00 | 5E-04 |
| Alaska | G5 | 2.70E-06 | 0E+00 | 1E-05 |

### d) population migration rate

| | Migration rates | | | | | | | | | |
|---|---|---|---|---|---|---|---|---|---|---|
| | *M01* | *M02* | *M03* | *M04* | *M05* | *M10* | *M20* | *M30* | M40 | *M50* |
| **value** | 0.001547 | 0.011045 | 0.002851 | 0.005435 | 0.008198 | 0.002655 | 0.000243 | 0.000089 | 0.004508 | 0.002085 |
| **CI2.5** | 0 | 0 | 0 | 0 | 0 | 0 | 0 | 0 | 0 | 0 |
| **CI97.5** | 0.0024 | 0.0119 | 0.0036 | 0.0065 | 0.0091 | 0.0033 | 0.0011 | 0.0011 | 0.0055 | 0.0028 |

| | Migration rates | | | | | | | |
|---|---|---|---|---|---|---|---|---|
| | *M12* | *M13* | *M14* | *M15* | *M21* | *M31* | *M41* | *M51* |
| **value** | 0.00853 | 0.00250 | 0.00244 | 0.00534 | 0.00009 | 0.00006 | 0.00086 | 0.00136 |
| **CI2.5** | 0 | 0 | 0 | 0 | 0 | 0 | 0 | 0 |
| **CI97.5** | 0.0093 | 0.0032 | 0.0034 | 0.0061 | 0.0007 | 0.0009 | 0.0020 | 0.0020 |

| | Migration rates | | | | | |
|---|---|---|---|---|---|---|
| | *M23* | *M24* | *M25* | *M32* | *M42* | *M52* |
| **value** | 9.65E-05 | 2.27E-04 | 1.12E-03 | 4.18E-04 | 9.35E-03 | 5.78E-03 |
| **CI2.5** | 0 | 0 | 0 | 0 | 0 | 0 |
| **CI97.5** | 0.00086 | 0.00108 | 0.00254 | 0.00121 | 0.01035 | 0.00680 |

| | Migration rates | | | | | |
|---|---|---|---|---|---|---|
| | *M34* | *M43* | *M35* | *M53* | *M45* | *M54* |
| **value** | 0.0001 | 0.0073 | 0.0000 | 0.0040 | 0.0030 | 0.0008 |
| **CI2.5** | 0 | 0 | 0 | 0 | 0 | 0 |
| **CI97.5** | 0.00119 | 0.00838 | 0.00101 | 0.00485 | 0.00382 | 0.00176 |
