## Supplementary material for "Demographic history shaped geographical patterns of deleterious mutation load in a broadly distributed Pacific Salmon": SupTab: TableS01.pdf

| | Pop. abbreviation | River | Region | nb. inds. | Longitude | Latitude |
|---|---|---|---|---|---|---|
| 1 | ALA | Alagnad | Alaska | 13 | –156.4657 | 59.09397 |
| 2 | ALB | Albreda | Thompson | 22 | –119.1308 | 52.47861 |
| 3 | ALO | Alouette | BC | 44 | –122.7086 | 49.26444 |
| 4 | ARC | Achuelinguk | Alaska | 20 | –163.7190 | 62.17600 |
| 5 | AVO | Avola | Thompson | 19 | –119.3178 | 51.77750 |
| 6 | BEL | Bellabella | BC | 44 | –128.1203 | 52.15472 |
| 7 | BIR | Birkenhead | Thompson | 27 | –122.6056 | 50.30611 |
| 8 | BNV | Bonneville | Cascadia | 36 | –121.8108 | 45.70641 |
| 9 | BON | Bonaparte | Thompson | 44 | –121.2586 | 50.73917 |
| 10 | CAM | Campbell | California | 38 | –123.7155 | 39.51222 |
| 11 | CHE | Chenails | BC | 45 | –121.9358 | 49.26972 |
| 12 | CHW | Chiliwak | BC | 42 | –121.9639 | 49.09722 |
| 13 | CLA | Clackamas | Cascadia | 26 | –122.6069 | 45.37253 |
| 14 | COW | Cowichan | BC | 39 | –123.6367 | 48.75333 |
| 15 | CWL | Cowlitz | Cascadia | 34 | –122.9175 | 46.09564 |
| 16 | EAG | Eagles | Thompson | 43 | –119.0108 | 50.84306 |
| 17 | ELE | Elochomon | Cascadia | 30 | –123.4197 | 46.23889 |
| 18 | GAT | Gates | Thompson | 34 | –122.4722 | 50.55194 |
| 19 | GNA | Gnat | Cascadia | 20 | –123.5291 | 46.17969 |
| 20 | HOP | Hope | Cascadia | 31 | –123.2050 | 46.61600 |
| 21 | INC | Inch | BC | 35 | –122.1561 | 49.17028 |
| 22 | KAN | Kanaka | BC | 27 | –122.5836 | 49.19917 |
| 23 | KAS | Kasiks | BC | 36 | –129.4006 | 54.29306 |
| 24 | KEN | Kenai | Alaska | 26 | –150.5346 | 60.46024 |
| 25 | KEO | Keogh | BC | 41 | –127.3483 | 50.67750 |
| 26 | KWE | Kwethluck | Alaska | 26 | –161.0985 | 60.49574 |
| 27 | LAC | Lachmach | BC | 29 | –129.9794 | 54.30167 |
| 28 | LEM | Lemieux | Thompson | 43 | –120.2006 | 51.42417 |
| 29 | LOU | Louis | Thompson | 44 | –120.1283 | 51.13750 |
| 30 | MAM | Mamquam | BC | 40 | –123.1544 | 49.73333 |
| 31 | MCG | McGarvey | California | 45 | –123.9959 | 41.50229 |
| 32 | MCK | McKinley | Thompson | 35 | –121.0697 | 52.28889 |
| 33 | MSL | Mile Slough | Alaska | 20 | –149.1235 | 64.50224 |
| 34 | NAV | NavFac | Alaska | 25 | –176.6211 | 51.89991 |
| 35 | NEE | Neechanz | BC | 40 | –126.6869 | 51.64583 |
| 36 | NIC | Nicomekl | BC | 43 | –122.8694 | 49.05806 |
| 37 | NOO | Nooksack | BC | 37 | –122.5789 | 48.78056 |
| 38 | NOR | Norrisj | BC | 45 | –122.1353 | 49.17306 |
| 39 | OON | Oona | BC | 44 | –130.2561 | 53.94722 |
| 40 | PIG | Pig | Thompson | 25 | –119.8867 | 51.59828 |
| 41 | PUN | Puntledge | BC | 39 | –124.9947 | 49.69583 |
| 42 | QUA | Quatse | BC | 44 | –127.4800 | 50.69972 |
| 43 | ROS | Rosewall | BC | 44 | –124.7786 | 49.46667 |
| 44 | SAN | Sangan | HaidaGwaii | 40 | –131.9969 | 54.02722 |
| 45 | SCO | Scott | California | 35 | –122.2126 | 37.15240 |
| 46 | SER | Serpentine | BC | 34 | –122.8500 | 49.08306 |
| 47 | SHA | Sharber | California | 45 | –123.5652 | 40.89614 |
| 48 | SHO | Shovelnose | BC | 24 | –123.3433 | 50.06833 |
| 49 | SIL | Silverdale | BC | 19 | –122.3583 | 49.13306 |
| 50 | SIT | Situc | Alaska | 17 | –139.5703 | 59.44760 |
| 51 | SNA | Snake | Alaska | 29 | –165.5339 | 64.52440 |
| 52 | SOO | Sooke | BC | 37 | –123.6989 | 48.38389 |
| 53 | STA | Stave | BC | 50 | –122.4236 | 49.17167 |
| 54 | TLE | Tiell | HaidaGwaii | 17 | –131.9328 | 53.61861 |
| 55 | UPP | Upper little | California | 22 | –123.7912 | 39.27374 |
| 56 | UPT | Upper pit | BC | 19 | –122.7678 | 49.22861 |
| 57 | WAU | Waukwaas | BC | 40 | –127.4181 | 50.58806 |
| 58 | YAK | Yakoun | HaidaGwaii | 45 | –132.2058 | 53.65694 |
