## Supplementary material for "Demographic history shaped geographical patterns of deleterious mutation load in a broadly distributed Pacific Salmon": SupTab: TableS02.pdf

| Population | Region | $\beta_{ST}$ | 2.5%CI | 97.5%CI |
|---|---|---|---|---|
| CLA | Cascadia | –0.227 | –0.246 | –0.205 |
| UPP | California | –0.160 | –0.184 | –0.137 |
| MCG | California | –0.138 | –0.161 | –0.117 |
| UPT | BC | –0.137 | –0.153 | –0.121 |
| GNA | Cascadia | –0.076 | –0.094 | –0.058 |
| BNV | Cascadia | –0.069 | –0.087 | –0.051 |
| SHA | California | –0.066 | –0.092 | –0.041 |
| ELE | Cascadia | –0.058 | –0.076 | –0.042 |
| CWL | Cascadia | –0.046 | –0.066 | –0.028 |
| HOP | Cascadia | –0.013 | –0.030 | 0.005 |
| CAM | California | –0.007 | –0.030 | 0.018 |
| COW | BC | 0.025 | 0.011 | 0.039 |
| ROS | BC | 0.027 | 0.014 | 0.040 |
| SCO | California | 0.032 | 0.006 | 0.056 |
| QUA | BC | 0.045 | 0.031 | 0.056 |
| SOO | BC | 0.048 | 0.033 | 0.062 |
| PUN | BC | 0.049 | 0.036 | 0.060 |
| SER | BC | 0.056 | 0.041 | 0.071 |
| NIC | BC | 0.059 | 0.045 | 0.075 |
| CHW | BC | 0.060 | 0.046 | 0.073 |
| SIL | BC | 0.062 | 0.050 | 0.077 |
| MAM | BC | 0.064 | 0.051 | 0.076 |
| NOO | BC | 0.068 | 0.056 | 0.080 |
| WAU | BC | 0.072 | 0.057 | 0.086 |
| KAN | BC | 0.074 | 0.061 | 0.088 |
| SHO | BC | 0.075 | 0.062 | 0.089 |
| CHE | BC | 0.085 | 0.072 | 0.098 |
| NAV | Alaska | 0.087 | 0.071 | 0.106 |
| INC | BC | 0.094 | 0.079 | 0.108 |

| Population | Region | $\beta_{ST}$ | 2.5%CI | 97.5%CI |
|---|---|---|---|---|
| STA | BC | 0.095 | 0.083 | 0.108 |
| NOR | BC | 0.100 | 0.087 | 0.114 |
| KEO | BC | 0.106 | 0.093 | 0.117 |
| SIT | Alaska | 0.118 | 0.101 | 0.135 |
| ALO | BC | 0.120 | 0.108 | 0.132 |
| NEE | BC | 0.123 | 0.110 | 0.136 |
| BEL | BC | 0.128 | 0.112 | 0.144 |
| LAC | BC | 0.130 | 0.110 | 0.148 |
| YAK | HaidaGwaii | 0.159 | 0.144 | 0.173 |
| BIR | Thompson | 0.159 | 0.143 | 0.175 |
| KAS | BC | 0.167 | 0.153 | 0.181 |
| TLE | HaidaGwaii | 0.184 | 0.164 | 0.202 |
| SAN | HaidaGwaii | 0.184 | 0.167 | 0.200 |
| LEM | Thompson | 0.203 | 0.187 | 0.220 |
| ALA | Alaska | 0.208 | 0.188 | 0.227 |
| KWE | Alaska | 0.215 | 0.197 | 0.233 |
| GAT | BC | 0.215 | 0.199 | 0.231 |
| LOU | Thompson | 0.222 | 0.203 | 0.238 |
| KEN | Alaska | 0.224 | 0.205 | 0.242 |
| PIG | Thompson | 0.226 | 0.208 | 0.244 |
| ARC | Alaska | 0.226 | 0.206 | 0.244 |
| OON | BC | 0.235 | 0.214 | 0.257 |
| SNA | Alaska | 0.237 | 0.218 | 0.256 |
| AVO | Thompson | 0.242 | 0.226 | 0.259 |
| EAG | Thompson | 0.256 | 0.239 | 0.274 |
| MCK | BC | 0.256 | 0.238 | 0.273 |
| BON | Thompson | 0.259 | 0.242 | 0.278 |
| ALB | Thompson | 0.260 | 0.243 | 0.277 |
| MSL | Alaska | 0.416 | 0.395 | 0.437 |
