## Supplementary material for "Demographic history shaped geographical patterns of deleterious mutation load in a broadly distributed Pacific Salmon": SupTab: TableS03.pdf

| | pop1 | pop2 | model | AIC | deltaAIC | AICweights |
|---|---|---|---|---|---|---|
| 1 | BC | ALASKA | IM2N2m | 19844.205 | 0.000000 | 1.00000000 |
| 2 | BC | ALASKA | SC2N2m | 19993.683 | 149.477506 | 0.00000000 |
| 3 | BC | ALASKA | SC2N | 20091.367 | 247.161939 | 0.00000000 |
| 4 | BC | ALASKA | SC2m | 20249.553 | 405.347978 | 0.00000000 |
| 5 | BC | ALASKA | AM2m | 20280.495 | 436.289599 | 0.00000000 |
| 6 | BC | ALASKA | AM2N | 20550.417 | 706.211535 | 0.00000000 |
| 7 | BC | ALASKA | IM2N | 20588.718 | 744.513153 | 0.00000000 |
| 8 | BC | ALASKA | AM2N2m | 20927.582 | 1083.376859 | 0.00000000 |
| 9 | BC | ALASKA | IM2m | 20929.750 | 1085.544749 | 0.00000000 |
| 10 | BC | ALASKA | SC | 22844.723 | 3000.517754 | 0.00000000 |
| 11 | BC | ALASKA | AM | 24851.359 | 5007.154141 | 0.00000000 |
| 12 | BC | ALASKA | IM | 25565.566 | 5721.361238 | 0.00000000 |
| 13 | BC | ALASKA | SI2N | 29210.867 | 9366.661437 | 0.00000000 |
| 14 | BC | ALASKA | SI | 33802.872 | 13958.667201 | 0.00000000 |
| 15 | BC | THOMPSON | SC2N2m | 48069.467 | 0.000000 | 1.00000000 |
| 16 | BC | THOMPSON | SC2N | 51486.097 | 3416.629662 | 0.00000000 |
| 17 | BC | THOMPSON | SC2m | 51965.687 | 3896.220077 | 0.00000000 |
| 18 | BC | THOMPSON | IM2m | 53196.737 | 5127.269611 | 0.00000000 |
| 19 | BC | THOMPSON | AM2m | 53640.295 | 5570.827845 | 0.00000000 |
| 20 | BC | THOMPSON | IM2N | 53903.781 | 5834.313790 | 0.00000000 |
| 21 | BC | THOMPSON | AM2N | 55703.905 | 7634.437556 | 0.00000000 |
| 22 | BC | THOMPSON | AM2N2m | 56079.188 | 8009.720625 | 0.00000000 |
| 23 | BC | THOMPSON | IM2N2m | 58290.569 | 10221.102262 | 0.00000000 |
| 24 | BC | THOMPSON | SC | 70948.524 | 22879.056714 | 0.00000000 |
| 25 | BC | THOMPSON | IM | 76803.617 | 28734.149385 | 0.00000000 |
| 26 | BC | THOMPSON | AM | 76830.858 | 28761.391111 | 0.00000000 |
| 27 | BC | THOMPSON | SI2N | 83301.127 | 35231.660055 | 0.00000000 |
| 28 | BC | THOMPSON | SI | 106938.771 | 58869.303472 | 0.00000000 |
| 29 | CALIF1 | ALASKA | SC2N2m | 25572.807 | 0.000000 | 1.00000000 |
| 30 | CALIF1 | ALASKA | SC2N | 26456.256 | 883.448486 | 0.00000000 |
| 31 | CALIF1 | ALASKA | SC2m | 26533.032 | 960.224885 | 0.00000000 |
| 32 | CALIF1 | ALASKA | IM2m | 27280.087 | 1707.279151 | 0.00000000 |
| 33 | CALIF1 | ALASKA | AM2m | 27284.626 | 1711.818491 | 0.00000000 |
| 34 | CALIF1 | ALASKA | IM2N2m | 27547.074 | 1974.266136 | 0.00000000 |
| 35 | CALIF1 | ALASKA | IM2N | 28698.612 | 3125.804978 | 0.00000000 |
| 36 | CALIF1 | ALASKA | AM2N | 28707.275 | 3134.467809 | 0.00000000 |
| 37 | CALIF1 | ALASKA | AM2N2m | 29872.345 | 4299.537777 | 0.00000000 |
| 38 | CALIF1 | ALASKA | SC | 34102.388 | 8529.580673 | 0.00000000 |
| 39 | CALIF1 | ALASKA | SI2N | 36272.656 | 10699.848933 | 0.00000000 |
| 40 | CALIF1 | ALASKA | AM | 36332.367 | 10759.559548 | 0.00000000 |
| 41 | CALIF1 | ALASKA | IM | 36353.087 | 10780.279944 | 0.00000000 |
| 42 | CALIF1 | ALASKA | SI | 47957.956 | 22385.148200 | 0.00000000 |
| 43 | CALIF1 | THOMPSON | SC2N2m | 26987.504 | 0.000000 | 1.00000000 |
| 44 | CALIF1 | THOMPSON | IM2N2m | 27110.088 | 122.583641 | 0.00000000 |
| 45 | CALIF1 | THOMPSON | SC2N | 27132.774 | 145.270056 | 0.00000000 |
| 46 | CALIF1 | THOMPSON | SC2m | 28096.560 | 1109.055254 | 0.00000000 |
| 47 | CALIF1 | THOMPSON | IM2m | 28160.801 | 1173.296770 | 0.00000000 |
| 48 | CALIF1 | THOMPSON | AM2m | 28162.469 | 1174.964863 | 0.00000000 |
| 49 | CALIF1 | THOMPSON | AM2N | 30373.179 | 3385.674512 | 0.00000000 |
| 50 | CALIF1 | THOMPSON | IM2N | 30544.577 | 3557.072798 | 0.00000000 |
| 51 | CALIF1 | THOMPSON | AM2N2m | 31515.947 | 4528.443058 | 0.00000000 |
| 52 | CALIF1 | THOMPSON | SI2N | 37239.259 | 10251.754603 | 0.00000000 |
| 53 | CALIF1 | THOMPSON | SC | 38732.426 | 11744.922134 | 0.00000000 |
| 54 | CALIF1 | THOMPSON | AM | 43498.313 | 16510.809219 | 0.00000000 |
| 55 | CALIF1 | THOMPSON | IM | 43499.244 | 16511.739942 | 0.00000000 |
| 56 | CALIF1 | THOMPSON | SI | 58549.183 | 31561.678243 | 0.00000000 |
| 57 | CALIF2 | ALASKA | IM2N2m | 6433.225 | 0.000000 | 1.00000000 |
| 58 | CALIF2 | ALASKA | IM2m | 6494.858 | 61.633439 | 0.00000000 |
| 59 | CALIF2 | ALASKA | AM2m | 6507.062 | 73.836541 | 0.00000000 |
| 60 | CALIF2 | ALASKA | SC2m | 6507.635 | 74.409678 | 0.00000000 |
| 61 | CALIF2 | ALASKA | SC2N2m | 6640.349 | 207.123993 | 0.00000000 |
| 62 | CALIF2 | ALASKA | SC2N | 7073.012 | 639.786562 | 0.00000000 |
| 63 | CALIF2 | ALASKA | AM2N2m | 7170.000 | 736.774889 | 0.00000000 |
| 64 | CALIF2 | ALASKA | AM2N | 7724.173 | 1290.947796 | 0.00000000 |
| 65 | CALIF2 | ALASKA | IM2N | 7774.306 | 1341.080879 | 0.00000000 |
| 66 | CALIF2 | ALASKA | SI2N | 9006.647 | 2573.421854 | 0.00000000 |
| 67 | CALIF2 | ALASKA | SC | 10418.343 | 3985.117487 | 0.00000000 |
| 68 | CALIF2 | ALASKA | IM | 11314.702 | 4881.477437 | 0.00000000 |
| 69 | CALIF2 | ALASKA | AM | 11317.293 | 4884.068230 | 0.00000000 |
| 70 | CALIF2 | ALASKA | SI | 16285.398 | 9852.172722 | 0.00000000 |
| 71 | CALIF2 | THOMSPON | SC2N | 8298.807 | 0.000000 | 0.55182679 |
| 72 | CALIF2 | THOMSPON | SC2m | 8299.914 | 1.107383 | 0.31720319 |
| 73 | CALIF2 | THOMSPON | SC2N2m | 8302.665 | 3.858624 | 0.08015181 |
| 74 | CALIF2 | THOMSPON | AM2m | 8303.577 | 4.769959 | 0.05081822 |
| 75 | CALIF2 | THOMSPON | IM2m | 8446.370 | 147.563572 | 0.00000000 |
| 76 | CALIF2 | THOMSPON | AM2N | 9101.888 | 803.081513 | 0.00000000 |
| 77 | CALIF2 | THOMSPON | IM2N | 9301.264 | 1002.457303 | 0.00000000 |
| 78 | CALIF2 | THOMSPON | AM2N2m | 9669.669 | 1370.862027 | 0.00000000 |
| 79 | CALIF2 | THOMSPON | IM2N2m | 10271.232 | 1972.424888 | 0.00000000 |
| 80 | CALIF2 | THOMSPON | SC | 12167.780 | 3868.972767 | 0.00000000 |
| 81 | CALIF2 | THOMSPON | SI2N | 12472.883 | 4174.076179 | 0.00000000 |
| 82 | CALIF2 | THOMSPON | AM | 13400.245 | 5101.438004 | 0.00000000 |
| 83 | CALIF2 | THOMSPON | IM | 13454.332 | 5155.525317 | 0.00000000 |
| 84 | CALIF2 | THOMSPON | SI | 18807.596 | 10508.789620 | 0.00000000 |
| 85 | CASCADIA | ALASKA | IM2N2m | 18061.267 | 0.000000 | 1.00000000 |
| 86 | CASCADIA | ALASKA | SC2N2m | 18479.911 | 418.643488 | 0.00000000 |
| 87 | CASCADIA | ALASKA | IM2m | 18641.045 | 579.777430 | 0.00000000 |
| 88 | CASCADIA | ALASKA | SC2m | 18641.309 | 580.041240 | 0.00000000 |
| 89 | CASCADIA | ALASKA | AM2m | 18641.671 | 580.403269 | 0.00000000 |
| 90 | CASCADIA | ALASKA | AM2N | 19220.408 | 1159.140729 | 0.00000000 |
| 91 | CASCADIA | ALASKA | AM2N2m | 19559.113 | 1497.845639 | 0.00000000 |
| 92 | CASCADIA | ALASKA | SC2N | 19785.650 | 1724.382553 | 0.00000000 |
| 93 | CASCADIA | ALASKA | IM2N | 19988.890 | 1927.622431 | 0.00000000 |
| 94 | CASCADIA | ALASKA | SC | 26954.769 | 8893.501833 | 0.00000000 |
| 95 | CASCADIA | ALASKA | AM | 27843.785 | 9782.517840 | 0.00000000 |
| 96 | CASCADIA | ALASKA | IM | 28081.828 | 10020.561064 | 0.00000000 |
| 97 | CASCADIA | ALASKA | SI2N | 28622.606 | 10561.338496 | 0.00000000 |
| 98 | CASCADIA | ALASKA | SI | 38418.342 | 20357.074993 | 0.00000000 |
| 99 | CASCADIA | THOMPSON | SC2N2m | 45178.874 | 0.000000 | 1.00000000 |
| 100 | CASCADIA | THOMPSON | SC2m | 45770.743 | 591.869244 | 0.00000000 |
| 101 | CASCADIA | THOMPSON | IM2N2m | 45915.586 | 736.712185 | 0.00000000 |
| 102 | CASCADIA | THOMPSON | SC2N | 48529.476 | 3350.601914 | 0.00000000 |
| 103 | CASCADIA | THOMPSON | IM2m | 49366.029 | 4187.155208 | 0.00000000 |
| 104 | CASCADIA | THOMPSON | AM2m | 49750.269 | 4571.394987 | 0.00000000 |
| 105 | CASCADIA | THOMPSON | IM2N | 49918.979 | 4740.105519 | 0.00000000 |
| 106 | CASCADIA | THOMPSON | AM2N | 49999.779 | 4820.905250 | 0.00000000 |
| 107 | CASCADIA | THOMPSON | AM2N2m | 51598.557 | 6419.683017 | 0.00000000 |
| 108 | CASCADIA | THOMPSON | SC | 68639.241 | 23460.367732 | 0.00000000 |
| 109 | CASCADIA | THOMPSON | IM | 73214.641 | 28035.767308 | 0.00000000 |
| 110 | CASCADIA | THOMPSON | AM | 73217.413 | 28038.538967 | 0.00000000 |
| 111 | CASCADIA | THOMPSON | SI2N | 76358.501 | 31179.627001 | 0.00000000 |
| 112 | CASCADIA | THOMPSON | SI | 106549.480 | 61370.606596 | 0.00000000 |
| 113 | THOMPSON | ALASKA | SC2N2m | 33229.449 | 0.000000 | 1.00000000 |
| 114 | THOMPSON | ALASKA | SC2m | 33320.660 | 91.211487 | 0.00000000 |
| 115 | THOMPSON | ALASKA | IM2N2m | 33493.620 | 264.171214 | 0.00000000 |
| 116 | THOMPSON | ALASKA | AM2m | 33933.419 | 703.969832 | 0.00000000 |
| 117 | THOMPSON | ALASKA | IM2m | 34164.219 | 934.770307 | 0.00000000 |
| 118 | THOMPSON | ALASKA | AM2N2m | 35430.417 | 2200.967973 | 0.00000000 |
| 119 | THOMPSON | ALASKA | AM2N | 35618.409 | 2388.960248 | 0.00000000 |
| 120 | THOMPSON | ALASKA | SC2N | 35729.053 | 2499.603747 | 0.00000000 |
| 121 | THOMPSON | ALASKA | IM2N | 37487.554 | 4258.105555 | 0.00000000 |
| 122 | THOMPSON | ALASKA | SI2N | 51347.927 | 18118.477617 | 0.00000000 |
| 123 | THOMPSON | ALASKA | SC | 51809.634 | 18580.185332 | 0.00000000 |
| 124 | THOMPSON | ALASKA | AM | 57590.643 | 24361.194597 | 0.00000000 |
| 125 | THOMPSON | ALASKA | IM | 57596.897 | 24367.447626 | 0.00000000 |
| 126 | THOMPSON | ALASKA | SI | 75673.317 | 42443.867737 | 0.00000000 |
