## Supplementary material for "Demographic history shaped geographical patterns of deleterious mutation load in a broadly distributed Pacific Salmon": SupTab: TableS04.pdf

| | BC.vs.Alaska | BC.vs.Thompson | Cascadia.vs.Thompson | California1.vs.Alaska | California2.vs.Alaska | Cascadia.vs.Alaska | Thompson.vs.Alaska | California1.vs.Thompson | California2.vs.Thompson |
|---|---|---|---|---|---|---|---|---|---|
| *parameters* | IM2N2mG | SC2N2mG | SC2N2mG | SC2N2mG | IM2N2mG | IM2N2mG | SC2N2mG | SC2N2mG | SC2N2mG |
| *Nref* | 14515 | 14053 | 14602 | 14117 | 13032 | 14868,21 | 14498 | 6342 | 15882 |
| *hrf* | 0,100 | 0,063 | 0,108 | 0,085 | 0,989 | 0,36787015 | 5,88E−02 | 0,09853226 | 4,55E−02 |
| *P* | 0,650 | 0,894 | 0,688 | 0,501 | 0,159 | 0,05678748 | 0,71 | 0,59227584 | 5,57E−01 |
| *Q* | 0,205 | 0,497 | 0,496 | 0,438 | 0,013 | 0,49960248 | 4,97E−01 | 0,49851057 | 4,85E−01 |
| *Ne1* | 2063 | 110358 | 212220 | 41563 | 42510 | 240992 | 60467 | 44535 | 26896 |
| *Ne2* | 1437636 | 192787 | 18663 | 100090 | 259892 | 75767 | 147862 | 62606 | 42386 |
| *b1* | 12,27056 | 0,10027 | 0,10185 | 0,09006 | 0,07092 | 0,28000 | 1,18E−01 | 0,10004 | 1,00E−01 |
| *b2* | 0,02627 | 0,04231 | 0,24501 | 0,08489 | 0,98881 | 0,18000 | 4,21E−02 | 0,15580 | 9,36E+01 |
| *m1* | 0,0004660 | 0,0018288 | 0,0001807 | 0,0014686 | 0,0017725 | 0,0000146 | 1,798E−03 | 0,0001648 | 1,88E−03 |
| *m²* | 0,0000305 | 0,0013328 | 0,0019866 | 0,0005068 | 0,0006309 | 0,0000286 | 9,008E−04 | 0,0001080 | 0,0015 |
| *me1* | 0,0000464 | 0,00000906 | 0,00001029 | 0,00003566 | 0,00002267 | 0,00022756 | 2,053E−05 | 0,00000329 | 3,62E−05 |
| *me2* | 0,000000627 | 0,0000000000553 | 0,00003062 | 0,0000000000232 | 0,00000317 | 0,00097647 | 5,855E−07 | 0,00000129 | 3,08E−06 |
| *Tsplit* | 129453 | 157472 | 168333 | 100553 | 81848 | 158434 | 140394 | 160900 | 93627 |
| *Tsc* | NA | 14886 | 20151 | 15122 | NA | NA | 10782 | 10796 | 9245 |
| *N1b1* | 25309 | 11066 | 21615 | 3743 | 3015 | 67065 | 7159 | 4455 | 2703 |
| *N2B2* | 37772 | 8157 | 4573 | 8496 | 256983 | 13596 | 6227 | 9754 | 3966117 |
