## Supplementary material for "Demographic history shaped geographical patterns of deleterious mutation load in a broadly distributed Pacific Salmon": SupTab: TableS05.pdf

| Populations Comparison | model | Nref | hrf | P | Q | Ne1 | Ne2 | b1 | b2 | m1 | $m^2$ | me1 | me2 | Tsplit | Tsc |
|---|---|---|---|---|---|---|---|---|---|---|---|---|---|---|---|
| **BC vs Alaska** | **IM2N2mG** | **14515** | **1.0E-01** | **6.5E-01** | **2.1E-01** | **2063** | **1437636** | **12.27** | **0.03** | **0.000466** | **0.000031** | **0.000046** | **0.000001** | **129453** | **NA** |
| | | CI2.5 | 0.0E+00 | 0.0E+00 | 0.0E+00 | 1555 | 175680 | 2.02 | 0.00 | 0.000000 | 0.000000 | 0.000000 | 0.000000 | 32506 | NA |
| | | CI97.5 | 6.2E-01 | 1.0E+00 | 5.8E-01 | 2570 | 2699592 | 22.52 | 0.08 | 0.000954 | 0.000208 | 0.000222 | 0.000005 | 226400 | NA |
| **BC vs Thompson** | **SC2N2mG** | **30725** | **6.3E-02** | **8.9E-01** | **5.0E-01** | **241284** | **421506** | **0.10** | **0.04** | **0.000836** | **0.000610** | **0.000004** | **0.000000** | **344294** | **32547** |
| | | CI2.5 | 4.4E-03 | 7.7E-01 | 0.0E+00 | 144119.8 | 267016.9 | 0.00 | 0.01 | 0.000599 | 0.000515 | 0.000004 | 0.000000 | 132797 | 30698 |
| | | CI97.5 | 1.2E-01 | 1.0E+00 | 1.0E+00 | 338449 | 575995 | 0.22 | 0.08 | 0.001074 | 0.000704 | 0.000004 | 0.000002 | 555791 | 34397 |
| **Cascadia vs Thompson** | **SC2N2mG** | **14602** | **1.1E-01** | **6.9E-01** | **5.0E-01** | **212220** | **18663** | **0.10** | **0.25** | **0.000083** | **0.000909** | **0.000005** | **0.000014** | **168333** | **20151** |
| | | CI2.5 | 0.0E+00 | 0.0E+00 | 0.0E+00 | 0.0 | 0.0 | 0.00 | 0.00 | 0.000000 | 0.000000 | 0.000000 | 0.000000 | 0 | 0 |
| | | CI97.5 | 6.3E-01 | 1.0E+00 | 1.0E+00 | 1384178.7 | 203225.4 | 0.59 | 1.82 | 0.000638 | 0.014131 | 0.000305 | 0.000777 | 1305228 | 160550 |
| **California1 vs Alaska** | **SC2N2mG** | **14117** | **8.5E-02** | **5.0E-01** | **4.4E-01** | **41563** | **100090** | **0.09** | **0.08** | **0.001469** | **0.000507** | **0.000036** | **0.000000** | **100553** | **15122** |
| | | CI2.5 | 4.7E-02 | 5.0E-01 | 4.4E-01 | 24776 | 91134 | 0.00 | 0.00 | 0.000461 | 0.000025 | 0.000000 | 0.000000 | 0 | 1424 |
| | | CI97.5 | 1.2E-01 | 5.0E-01 | 4.4E-01 | 58349.5 | 109045.1 | 0.23 | 0.23 | 0.002476 | 0.000989 | 0.000087 | 0.000015 | 277697 | 28821 |
| **California2 vs Alaska** | **IM2N2mG** | **13032.0** | **9.9E-01** | **1.6E-01** | **1.3E-02** | **42510.0** | **259892.1** | **0.07** | **0.99** | **0.001772** | **0.000631** | **0.000023** | **0.000003** | **81848** | **NA** |
| | | CI2.5 | 9.7E-01 | 7.2E-02 | 0.0E+00 | 0 | 216187 | 0.00 | 0.98 | 0.000779 | 0.000274 | 0.000007 | 0.000002 | 70505 | NA |
| | | CI97.5 | 1.0E+00 | 2.5E-01 | 3.7E-02 | 86723 | 303597 | 0.24 | 0.99 | 0.002766 | 0.000987 | 0.000038 | 0.000004 | 93192 | NA |
| **Cascadia vs Alaska** | **IM2N2mG** | **14868.00** | **3.7E-01** | **5.7E-02** | **5.0E-01** | **240992** | **75767** | **0.28** | **0.18** | **0.000015** | **0.000029** | **0.000228** | **0.000976** | **158434** | **NA** |
| | | CI2.5 | -1.8E-03 | 0.0E+00 | 3.4E-01 | 0 | 69710 | 0.10 | 0.07 | 0.000008 | 0.000015 | 0.000118 | 0.000874 | 109924 | NA |
| | | CI97.5 | 7.4E-01 | 6.2E-01 | 6.6E-01 | 482169 | 81825 | 0.45 | 0.29 | 0.000021 | 0.000042 | 0.000337 | 0.001079 | 206943 | NA |
| Thompson vs Alaska | **SC2N2mG** | **14498** | **5.9E-02** | **7.1E-01** | **5.0E-01** | **60467** | **147862** | **0.12** | **0.04** | **0.001798** | **0.000901** | **0.000021** | **0.000001** | **140394** | **10782** |
| | | CI2.5 | 5.3E-02 | 1.2E-01 | 0.0E+00 | 0 | 0 | 0.00 | 0.00 | 0.001466 | 0.000752 | 0.000000 | 0.000000 | 121041 | 9494 |
| | | CI97.5 | 6.4E-02 | 1.0E+00 | 1.0E+00 | 206436 | 362758 | 0.61 | 0.15 | 0.002131 | 0.001050 | 0.000107 | 0.000039 | 159747 | 12070 |
| **California1 vs Thompson** | **SC2N2mG** | **6342** | **9.9E-02** | **5.9E-01** | **5.0E-01** | **44535** | **62606** | **0.10** | **0.16** | **0.002332** | **0.001529** | **0.000047** | **0.000018** | **160900** | **10796** |
| | | CI2.5 | 0.0E+00 | 0.0E+00 | 0.0E+00 | 0 | 0 | 0.00 | 0.00 | 0.000000 | 0.000000 | 0.000000 | 0.000000 | 4954 | 0 |
| | | CI97.5 | 2.7E-01 | 1.0E+00 | 1.0E+00 | 91658 | 128139 | 0.30 | 0.57 | 0.009028 | 0.004718 | 0.000204 | 0.000048 | 316846 | 22808 |
| **California2 vs Thompson** | **SC2N2mG** | **15882** | **4.5E-02** | **5.6E-01** | **4.8E-01** | **26896** | **42386** | **93.57** | **93.57** | **0.001882** | **0.001542** | **0.000036** | **0.000003** | **93627** | **9245** |
| | | CI2.5 | 4.1E-02 | 3.9E-01 | 7.3E-02 | 25693.2 | 34562.0 | 93.56 | 27.97 | 0.001116 | 0.001016 | 0.000017 | 0.000002 | 83855 | 8572 |
| | | CI97.5 | 5.0E-02 | 7.3E-01 | 9.0E-01 | 28098.7 | 50209.6 | 93.58 | 159.17 | 0.002647 | 0.002067 | 0.000055 | 0.000004 | 103398 | 9919 |
