## Supplementary material for "Demographic history shaped geographical patterns of deleterious mutation load in a broadly distributed Pacific Salmon": SupTab: TableS06.pdf

| Populations Comparison | best model | Nref | hrf | P | Q | Ne1 | Ne2 | b1 | b2 | m1 | $m^2$ | me1 | me2 | Tsplit | Tsc |
|---|---|---|---|---|---|---|---|---|---|---|---|---|---|---|---|
| | | 14515 | .10 | .65 | .21 | 2,063 | 1,437,636 | 12.27 | .03 | .000466 | .000031 | .000046 | .000001 | 129,453 | NA |
| **BC vs Alaska** | **IM2N2mG** | CI2.5 | .00 | .57 | .15 | | 1,286,048 | 2.94 | .00 | .000418332700 | .000000000000 | .000024582580 | .000000000000 | 110,054 | NA |
| | | CI97.5 | .20 | .73 | .26 | 287,391 | 1,589,224 | 21.61 | 5.95 | .000514 | .000076 | .000068 | .000033 | 148,852 | NA |
| | | 14053 | .06 | .89 | .50 | 110,358 | 192,787 | .10 | .04 | .001829 | .001333 | .000009 | .000000 | 157,472 | 14,886 |
| **BC vs Thompson** | **SC2N2mG** | CI2.5 | .00 | .85 | .46 | 53,566 | 74,508 | .00 | .00 | .000000000000 | .000435190300 | .000000000000 | .000000000000 | 130,845 | 11,581 |
| | | CI97.5 | .15 | .94 | .54 | 167,150 | 311,066 | 7.47 | 6.41 | .003941 | .002230 | .001107 | .000524 | 184,099 | 18,191 |
| | | 14602 | .11 | .69 | .50 | 212,220 | 18,663 | .10 | .25 | .000181 | .001987 | .000010 | .000031 | 168,333 | 20,151 |
| **Cascadia vs Thompson** | **SC2N2mG** | CI2.5 | .02 | .65 | .46 | 22,122 | .00 | .00 | .00 | .000164319448 | .001935638088 | .000000000000 | .000002977441 | .00 | .00 |
| | | CI97.5 | .19 | .73 | .54 | 402,318 | 1,044,390 | 3.98 | 4.97 | .000197 | .002038 | .000025 | .000058 | 356,301 | 89,641 |
| | | 14117 | .09 | .50 | .44 | 41,563 | 100,090 | .09 | .08 | .001469 | .000507 | .000036 | .000000 | 100,553 | 15,122 |
| **California1 vs Alaska** | **SC2N2mG** | CI2.5 | .00 | .47 | .39 | .00 | 50,195 | .00 | .00 | .000916094700 | .000029385760 | .000000000000 | .000000000000 | 89,785 | 11,676 |
| | | CI97.5 | .17 | .53 | .49 | 94,639 | 149,985 | 8.09 | 3.87 | .002021 | .000984 | .000596 | .000442 | 111,321 | 18,568 |
| | | 13032 | .99 | .16 | .01 | 42,510 | 259,892 | .07 | .99 | .001773 | .000631 | .000023 | .000003 | 81,848 | NA |
| **California2 vs Alaska** | **IM2N2mG** | CI2.5 | .89 | .09 | .00 | .00 | 175,841 | .00 | .00 | .000000000000 | .000000000000 | .000000000000 | .000000000000 | 69,904 | NA |
| | | CI97.5 | 1.09 | .23 | .06 | 97,535 | 343,943 | 12.13 | 9.28 | .053773 | .051028 | .000360 | .000482 | 93,792 | NA |
| | | **14868.21** | .37 | .06 | .50 | 240,992 | 75,767 | .28 | .18 | .000015 | .000029 | .000228 | .000976 | 158,434 | NA |
| **Cascadia vs Alaska** | **IM2N2mG** | CI2.5 | .29 | .00 | .00 | .00 | .00 | .00 | .00 | .000000 | .000000 | .000206 | .000951 | 109,819 | NA |
| | | CI97.5 | .45 | .14 | .55 | 822,684 | 226,204 | 8.23 | 10.83 | .000075 | .000089 | .000249 | .001002 | 207,049 | NA |
| | | **14498** | .06 | .71 | .50 | 60,467 | 147,862 | .12 | .04 | .001798 | .000901 | .000021 | .000001 | 140,394 | 10,782 |
| Thompson vs Alaska | **SC2N2mG** | CI2.5 | .00 | .67 | .45 | 57,955 | 144,899 | .00 | .00 | .001049764436 | .000159933306 | -.000222958392 | -.000315721400 | 98,995 | 7,738 |
| | | CI97.5 | .12 | .75 | .55 | 62,979 | 150,825 | 8.41 | 2.24 | .002546 | .001642 | .000264 | .000317 | 181,793 | 13,826 |
| | | 6342 | .10 | .59 | .50 | 44,535 | 62,606 | .10 | .16 | .000165 | .000108 | .000003 | .000001 | 160,900 | 10,796 |
| **California1 vs Thompson** | **SC2N2mG** | CI2.5 | .03 | .56 | .45 | 42,494 | .00 | .00 | .00 | .000000000000 | .000000000000 | .000000000000 | .000000000000 | 94,540 | .00 |
| | | CI97.5 | .17 | .63 | .55 | 46,576 | 348,605 | 6.09 | 6.08 | .001004 | .000559 | .000427 | .000310 | 227,260 | 22,388 |
| | | **15882** | .05 | .56 | .49 | 26,896 | 42,386 | .10 | 93.60 | .001880 | .001542 | .000036 | .000003 | 93,627 | 9,245 |
| **California2 vs Thompson** | **SC2N2mG** | CI2.5 | .00 | .52 | .45 | 24,514 | .00 | .00 | 88.16 | .000858386548 | .001165775713 | .000000000000 | .000000000000 | 40,071 | 1,451 |
| | | CI97.5 | .13 | .60 | .52 | 29,278 | 105,663 | 1.30 | 99.04 | .002902 | .001917 | .011106 | .010710 | 147,183 | 17,039 |
