## Supplementary material for "Demographic history shaped geographical patterns of deleterious mutation load in a broadly distributed Pacific Salmon": SupTab: TableS08.pdf

| | POP_ID | Tajima | piNpiS |
|---|---|---|---|
| 1 | ALA | 0.187365468 | 0.381261 |
| 2 | ALB | 0.466564035 | 0.325531 |
| 3 | ALO | 0.229149035 | 0.322075 |
| 4 | ARC | 0.260862436 | 0.345108 |
| 5 | AVO | 0.463035332 | 0.329969 |
| 6 | BEL | 0.447603283 | 0.293750 |
| 7 | BIR | 0.365854892 | 0.289335 |
| 8 | BNV | −0.079362001 | 0.372620 |
| 9 | BON | 0.362099913 | 0.354392 |
| 10 | CAM | 0.464932330 | 0.283914 |
| 11 | CHE | 0.090980224 | 0.317567 |
| 12 | CHW | 0.214782274 | 0.274725 |
| 13 | CLA | −0.372686148 | 0.500000 |
| 14 | COW | −0.054353484 | 0.315625 |
| 15 | CWL | 0.192948387 | 0.422680 |
| 16 | EAG | 0.576448901 | 0.284422 |
| 17 | ELE | 0.233001858 | 0.353256 |
| 18 | GAT | 0.352461858 | 0.372819 |
| 19 | GNA | 0.144992381 | 0.320000 |
| 20 | HOP | 0.357628028 | 0.314035 |
| 21 | INC | 0.277634020 | 0.301102 |
| 22 | KAN | 0.147261684 | 0.303529 |
| 23 | KAS | 0.419060320 | 0.304195 |
| 24 | KEN | 0.203383346 | 0.409246 |
| 25 | KEO | 0.251448982 | 0.311008 |
| 26 | KWE | 0.244772151 | 0.331601 |
| 27 | LAC | 0.381204949 | 0.278434 |
| 28 | LEM | 0.207810418 | 0.373368 |
| 29 | LOU | −0.145970051 | 0.406574 |

| | POP_ID | Tajima | piNpiS |
|---|---|---|---|
| 30 | MAM | 0.402383857 | 0.267736 |
| 31 | MCG | 0.458720823 | 0.288062 |
| 32 | MCK | −0.007417193 | 0.348555 |
| 33 | MSL | 0.350958071 | 0.342482 |
| 34 | NAV | −0.037038323 | 0.459682 |
| 35 | NEE | 0.345438523 | 0.300408 |
| 36 | NIC | 0.393628610 | 0.283748 |
| 37 | NOO | 0.387893622 | 0.292490 |
| 38 | NOR | −0.238776701 | 0.396441 |
| 39 | OON | 0.427071987 | 0.313803 |
| 40 | PIG | 0.578726614 | 0.287169 |
| 41 | PUN | 0.260415676 | 0.281618 |
| 42 | QUA | 0.229889028 | 0.290983 |
| 43 | ROS | 0.447902494 | 0.269681 |
| 44 | SAN | 0.115213495 | 0.400444 |
| 45 | SCO | 0.521973292 | 0.314669 |
| 46 | SER | 0.420390766 | 0.279170 |
| 47 | SHA | 0.459435869 | 0.311838 |
| 48 | SHO | 0.281279225 | 0.297364 |
| 49 | SIL | 0.266100283 | 0.281350 |
| 50 | SIT | 0.138373100 | 0.390262 |
| 51 | SNA | 0.347318341 | 0.377973 |
| 52 | SOO | 0.337848349 | 0.295138 |
| 53 | STA | 0.383963662 | 0.293548 |
| 54 | TLE | 0.422858263 | 0.303370 |
| 55 | UPP | 0.471853778 | 0.282271 |
| 56 | UPT | −0.892286728 | 0.581892 |
| 57 | WAU | −0.111326790 | 0.384168 |
| 58 | YAK | −0.111598949 | 0.402393 |
