## Supplementary material for "Demographic history shaped geographical patterns of deleterious mutation load in a broadly distributed Pacific Salmon": SupTab: TableS09.pdf

| | DAF of deleterious mutations | Count of deleterious mutation by population | Number of homozygous mutation by individuals | Number of heterozygous mutation by sample | Total number of deleterious mutation by sample |
|---|---|---|---|---|---|
| *Cascadia* | 0.102 | 127.86 | 3.23 | 14.43 | 17.65 |
| *California* | 0.196 | 66.20 | 4.88 | 14.02 | 18.89 |
| *BC* | 0.1301 | 96.26 | 2.86 | 13.85 | 16.70 |
| *HaidaGwaii* | 0.164 | 81.66 | 4.70 | 13.54 | 18.24 |
| *Thompson* | 0.200 | 62.37 | 3.75 | 12.20 | 15.95 |
| *Alaska* | 0.169 | 80.875 | 4.77 | 12.16 | 16.94 |
