## Supplementary material for "Demographic history shaped geographical patterns of deleterious mutation load in a broadly distributed Pacific Salmon": SupTab: TableS10.pdf

| region1 | region2 | W | P.values |
|---|---|---|---|
| cascadia | BC | 829630 | 1.3616e−01 |
| cascadia | california | 865050 | 1.0000e−06 |
| cascadia | haida gwaii | 863390 | 4.1100e−06 |
| cascadia | thompson | 898590 | 2.0000e−16 |
| california | alaska | 829970 | 1.2000e−01 |
| BC | california | 849220 | 5.0000e−04 |
| BC | haida gwaii | 846390 | 1.4000e−03 |
| BC | thompson | 882880 | 1.0000e−12 |
| BC | alaska | 810580 | 9.3500e−01 |
| california | haida gwaii | 773990 | 5.0000e−04 |
| california | thompson | 844170 | 2.7430e−03 |
| california | alaska | 809090 | 8.0600e−01 |
| haida gwaii | thompson | 739730 | 1.1000e−12 |
| haida gwaii | alaska | 775970 | 1.1000e−03 |
| thompson | alaska | 846820 | 9.9000e−05 |
