## Supplementary material for "Demographic history shaped geographical patterns of deleterious mutation load in a broadly distributed Pacific Salmon": SupTab: TableS11.pdf

| | | | homozygous derived | | total load | |
|---|---|---|---|---|---|---|
| | **region1** | **region2** | W | P−values | W | P−values |
| 1 | cascadia | BC | 5320.5 | 0.4152 | 5288 | 0.4806 |
| 2 | cascadia | california | 3117.5 | 2.53E−06 | 3776 | 2.70E−03 |
| 3 | cascadia | haida gwaii | 3785.5 | 2.20E−03 | 3978.5 | 0.0122 |
| 4 | cascadia | thompson | 3993 | 1.12E−02 | 5560 | 0.17 |
| 5 | cascadia | alaska | 3121.5 | 2.57E−06 | 5691.5 | 0.09 |
| 6 | BC | california | 2822 | 4.90E−08 | 3379 | 6.90E−05 |
| 7 | BC | haida gwaii | 3461.5 | 1.00E−04 | 3589.5 | 5.00E−04 |
| 8 | BC | thompson | 3651 | 6.60E−04 | 5260 | 0.52 |
| 9 | BC | alaska | 2809 | 3.90E−08 | 5434.5 | 0.287 |
| 10 | california | haida gwaii | 5802.5 | 0.044 | 5234.5 | 0.565 |
| 11 | california | thompson | 6059 | 7.80E−03 | 6784 | 1.22E−05 |
| 12 | california | alaska | 5057.5 | 8.51E−01 | 6878.5 | 4.13E−06 |
| 13 | haida gwaii | thompson | 5252 | 5.26E−01 | 7270 | 2.63E−08 |
| 14 | haida gwaii | alaska | 4244.5 | 5.78E−02 | 6554 | 1.00E−04 |
| 15 | thompson | alaska | 3974.5 | 9.80E−03 | 4427.5 | 0.65 |
